# Characterizing gene tree conflict in plastome-inferred phylogenies

**DOI:** 10.1101/512079

**Authors:** Joseph F. Walker, Gregory W. Stull, Nathanael Walker-Hale, Oscar M. Vargas, Drew A. Larson

## Abstract

**Premise of the study:** Evolutionary relationships among plants have been inferred primarily using chloroplast data. To date, no study has comprehensively examined the plastome for gene tree conflict.

**Methods:** Using a broad sampling of angiosperm plastomes, we characterized gene tree conflict among plastid genes at various time scales and explore correlates to conflict (e.g., evolutionary rate, gene length, molecule type).

**Key results:** We uncover notable gene tree conflict against a backdrop of largely uninformative genes. We find gene length is the strongest correlate to concordance, and that nucleotides outperform amino acids. Of the most commonly used markers, *matK* greatly outperforms *rbcL*; however, the rarely used gene *rpoC2* is the top-performing gene in every analysis. We find that *rpoC2* reconstructs angiosperm phylogeny as well as the entire concatenated set of protein-coding chloroplast genes.

**Conclusions:** Our results suggest that longer genes are superior for phylogeny reconstruction. The alleviation of some conflict through the use of nucleotides suggests that systematic error is likely the root of most of the observed conflict, but further research on biological conflict within plastome is warranted given the documented cases of heteroplasmic recombination. We suggest *rpoC2* as a useful marker for reconstructing angiosperm phylogeny, reducing the effort and expense of assembling and analyzing entire plastomes.

---

Chloroplast data have been the most prominent source of information for plant phylogenetics, largely due to the ease with which chloroplast genes can be sequenced, assembled, and analyzed (Palmer, 1985; Taberlet et al., 1991). The majority of broad-scale phylogenetic studies on plants have used chloroplast genes (e.g., Chase et al., 1993, Soltis et al., 2000, 2011), and the resulting phylogenies have been used for countless other comparative studies examining ancestral states, historical biogeography, and other evolutionary patterns. While older studies relied mostly on targeted genes such as *rbcL* and *matK*, recent advances in DNA sequencing have drastically increased the ease and affordability of whole-chloroplast genome (i.e., plastome) sequencing (Moore et al. 2006; Cronn et al., 2008, 2012; Stull et al., 2013; Uribe-Convers, et al. 2014), increasing the number of studies employing plastome-scale data for phylogenetic and comparative analyses (e.g., Jansen et al., 2007; Moore et al., 2007, 2010; Ruhfel et al., 2014; Stull et al., 2015; Gitzendanner et al., 2018). Nonetheless, the utility of plastid genes, as well as the entire plastome, is ultimately determined by the extent to which they reflect ‘true’ evolutionary relationships (i.e., the ‘species tree’) of the lineages in question (Doyle, 1992).

Phylogenomic conflict (i.e., the presence of conflicting relationships among gene trees in a genomic dataset) is now recognized as a nearly ubiquitous feature of nuclear phylogenomic studies (Smith et al., 2015). Most gene tree conflict is attributed to biological causes such as incomplete lineage sorting, hybridization, and gene duplication and loss (Maddison, 1997; Galtier and Daubin, 2008; Smith et al., 2015; Walker et al., 2017; Vargas et al., 2017). The genes within the plastome, however, are generally thought to be free of such biological sources of conflict. This is because the plastome is typically uniparentally inherited (maternally in angiosperms, paternally in conifers: Mogensen, 1996) and undergoes a unique form of recombination that is not expected to result in conflicting gene histories within a single genome (Palmer, 1983; Bendich, 2004; Walker et al. 2015).

However, in angiosperms, nonmaternal inheritance is relatively common (e.g., Smith, 1989; McCauley et al., 2007), and several surveys of pollen in flowering plants (Corriveau and Coleman, 1988; Zhang et al., 2003) have documented plastid DNA in up to 18% of the species examined, indicating potential for biparental inheritance and heteroplasmy (i.e., the presence of two or more different plastomes in a single organism, cell, or organelle). Heteroplasmy, which has been documented in multiple angiosperm species (e.g., Johnson and Palmer, 1988; Lee et al., 1988; Hansen et al., 2007; Carbonell-Caballero et al., 2015), creates an opportunity for heteroplasmic recombination, which could result in gene tree conflict in plastomes-inferred phylogenies. Heteroplasmic recombination (both intra- and interspecific) has been invoked by multiple studies to explain discordance in various clades (Huang et al., 2001; Marshall et al., 2001; Erixxon and Oxelman, 2008; Bouillé et al., 2011), but only a few recent studies have documented clear evidence of this phenonmenon (Sullivan et al., 2017; Sancho et al., 2018). Beyond heteroplasmy, sharing of genes between the chloroplast and nuclear genomes remains another potential source of biological conflict (Martin et al. 1998; Martin 2003 Stegemann et al., 2003). Although biological conflict in the plastome generally seems rare, the true extent of intra-plastome conflict is poorly known. Quantifying its extent is of high importance given that the vast majority of studies assume no conflict as an operating principle (Wolfe and Randle, 2004).

Aside from biological sources of conflict, there also remain significant potential sources of systematic conflict that have been poorly explored across the plastome (e.g., Burleigh and Mathews 2007a,b). Chloroplast data are used at various time scales, and the accumulation of substitutions over long periods of evolutionary time increases the probability of encountering systematic error due to saturation (Rodríguez-Ezpeleta et al., 2007; Philippe et al., 2011). Conflict has been demonstrated among different functional groups of genes (Liu et al., 2012), among different regions of the plastome (Walker et al., 2014), as well as among individual genes (e.g., Shepherd et al., 2008, Foster et al. 2018). The rate of chloroplast evolution as a whole has been examined (and compared with the nuclear and mitochondrial genomes; Wolfe, 1987), and rate variation within the chloroplast—especially across the three major regions of the genome, i.e., the long single-copy (LSC) region, the short single-copy (SSC) region, and the inverted repeats (IRa, IRb)—has been explored to help determine the markers useful for phylogenetic inference at different time scales (e.g., Graham and Olmstead, 2000; Shaw et al., 2005, 2007, 2014). However, no study has comprehensively examined gene tree conflict within the plastome to better characterize the extent and sources of conflict, and to identify the plastid genes most concordant with our current understanding of angiosperm phylogeny (e.g., Soltis et al., 2011; Wickett et al., 2014; Zeng et al., 2014; Gitzendanner et al., 2018; but see Logacheva et al. [2007] for a preliminary investigation of the concordance of individual plastid genes with angiosperm phylogeny).

Here we use phylogenomic tools to characterize the extent of conflict among plastid genes as a function of evolutionary rate, rate variation among species, sequence length, and data type (i.e., nucleotides vs. amino acids) at varying time scales across angiosperms. Our results show that the plastome—at all levels—contains notable gene tree conflict, with the number of conflicting genes at each node often comparable to the number of concordant genes; however, the majority of plastid genes are uninformative for most nodes when considering support. Information content (gene length and molecule type, i.e., nucleotides vs amino acids) was the strongest correlate with concordance, suggesting that most observed gene-tree conflict is a consequence of spurious inferences from insufficient information. However, many nodes across angiosperm phylogeny show at least several strongly supported conflicting genes, indicating the need for future investigations into the causes of intraplastome conflict (e.g., inappropriate models, heteroplasmic recombination, horizontal gene transfer). We also document the performance of individual genes at recapitulating angiosperm phylogeny, finding the seldom-used gene *rpoC2* to outperform commonly used genes (e.g., *rbcL*, *matK*) in all cases, consistent with previous work highlighting the utility of this gene (e.g., Logacheva et al., 2007). Our results provide an important glimpse into the extent and sources of intraplastome conflict.

## MATERIALS AND METHODS

### Data acquisition and sampling

Complete plastome coding data (both nucleotide and amino acid) were downloaded from NCBI for 53 taxa: 51 angiosperm ingroups and two gymnosperm outgroups (*Ginkgo Biloba* and *Podocarpus lambertii*; Table S1). Our sampling scheme was designed to capture all major angiosperm lineages (e.g., Soltis et al., 2011), while also including denser sampling for nested clades in Asterales. This allowed us to evaluate the extent of gene tree conflict at different evolutionary levels/time scales, from species-level relationships in *Diplostephium* (Asteraceae) to the ordinal-level relationships defining the backbone of angiosperm phylogeny.

### Data preparation, alignment, and phylogenetic inference

All scripts developed for this study may be found on GitHub (https://github.com/jfwalker/ChloroplastPhylogenomics). Orthology was determined based upon the annotations of protein-coding genes on Genbank; this resulted in almost complete gene occupancy apart from instances of gene loss or reported pseudogenization. We did not use non-coding regions, as proper orthology at deep time scales can be difficult to assess due to gene re-arrangements and inversions; this is in part why most deep phylogenetic analyses of angiosperms using plastomes have excluding non-coding data (e.g., Moore et al., 2010; Ruhfel et al., 2014).

The amino acid and nucleotide data were aligned using Fast Statistical Alignment (FSA; Bradley *et al*., 2009) with the default settings for peptide and the setting “--noanchored” for nucleotide. FSA has been shown to be one of the top-performing alignment programs (Redelings, 2014), and does not rely upon a guide tree for sequence alignment, alleviating downstream bias. A maximum likelihood (ML) tree was then inferred for each gene using RAxMLv.8.2.4 (Stamatakis, 2014), with the PROTGAMMAAUTO and GTR+G models of evolution used for the amino acid and nucleotide data, respectively. For each dataset, we conducted 200 rapid bootstrap replicates. The alignments were also concatenated into supermatrices and partitioned by gene using the phyx program pxcat (Brown et al. 2017). The nucleotide and amino acid supermatrices were then each used to infer ‘plastome’ trees using the GTR+G and the PROTGAMMAAUTO models, respectively, as implemented in RAxML. To complement the model inference performed by RAxML from the AUTO feature, we also used IQ-TREE’s (Nguyen et al. 2014) built-in model selection process (Kalyaanamoorthy et al. 2017) on the partitioned data. We did not partition by codon position because the lengths of most of the plastid genes, especially when divided into three smaller partitions, would have been insufficient to inform an evolutionary model.

### Reference phylogenies

For the conflict analyses, described below, we created several reference trees against which the gene trees were mapped. Primarily, we used a topology based on Soltis et al. (2011) and—for species-level relationships within Asteraceae—Vargas et al. (2017); we refer to this is our ‘accepted tree’, or AT. We also created a reference tree based on the recent plastome phylogeny of Gitzendanner et al. (2018), which we refer to as the ‘Gitzendanner tree’, or GT. The GT is largely identical to the AT, but several deep (and commonly contentions) nodes differ: e.g., the placement of Buxales and ordinal relationships in lamiids (here, we only sampled Lamiales, Gentianales, and Solanales). In the latter case, all three possible combinations of lamiids taxa were assessed (discussed further below in Assesment of Supported Conflict). Although it is difficult to determine which of these trees better represents species relationships of angiosperms, they both serve as useful frameworks for revealing conflict among genes. Here, we primarily focus analyses using the AT, but conflict analyses using the GT were also conduct to ensure that our overall results were consistent across reference trees.

### Analysis of conflict

All gene trees were rooted on the outgroups using the phyx program pxrr (Brown et al. 2017), in a ranked fashion (“-r”) in the order *Podocarpus lambertii* then *Ginkgo biloba*; this was performed to account for any missing genes in either of the taxa. Conflict in the data was identified using the bipartition method as implemented in phypartsv.0.0.1 (Smith et al. 2015), with the gene trees from each data set (amino acid and nucleotide) mapped against the AT and GT references trees. The concordance analyses were performed using both a support cutoff (at 70% bootstrap support, i.e., moderate support) and no support cutoff. When the support cutoff is used, any gene tree node with under 70 bootstrap support is regarded as uninformative for the reference node in question (i.e., it is uncertain whether the gene tree node is in conflict or concordance with the reference node); when no support cutoff is used, the gene tree node is evaluated as conflicting or concordant regardless of the support value. However, in both cases, a gene is considered uninformative if a taxon relevant to a particular node/relationship is missing from the gene dataset. For example, consider the reference tree ((A,B),C); if a gene tree is missing taxon B, but contains the relationship (A,C), then the gene tree is concordant with the node ((A,B),C) but uninformative regarding the node (A,B). See Smith et al. (2015) for an in-depth explanation of this method.

Levels of conflict were examined across the entire tree and within particular time intervals of the evolutionary history of angiosperms. For the latter case, the ~150 Ma of crown angiosperm evolution were divided into five time intervals (30 Ma each): 150 to 121 mya, 120 to 91 mya, 90 to 61 mya, 60 to 31 mya, and 30 to 0 mya. The nodes across the tree were binned into the time intervals based on their inferred ages (ages > 20 Ma from Magallón *et al*., 2015; ages < 20 Ma from Vargas *et al*., 2017 and Roquet *et al*., 2009). At each time interval, the proportion of concordant nodes for each gene was calculated (for all the nodes falling within that time interval). This allowed us to assess the level(s) of divergence at which each gene is most informative. We also determined concordance levels of the ‘plastome’ trees (against the reference) within each of these time intervals.

### Assessments of Individual Plastid Genes

Using the phyx program pxlstr (Brown *et al.* 2017), we calculated summary statistics for each gene alignment and corresponding gene tree: number of included species, alignment length, tree length (a measure of gene evolutionary rate), and root-to-tip variance (a measure of rate variation across the phylogeny). Alignment length and tree length represent different measures of a gene’s information content. Levels of concordance of each gene tree with the reference trees were then assessed by tabulating the number of nodes concordant between the gene tree and the given reference tree (e.g., the AT). The number of concordant nodes (in Fig. 2 this is treated as a proportion of total nodes available to support and in Fig. 3 this is based on total nodes) was used as a measure of the gene’s ability to accurately reconstruct angiosperm phylogeny.

**Figure 1.**
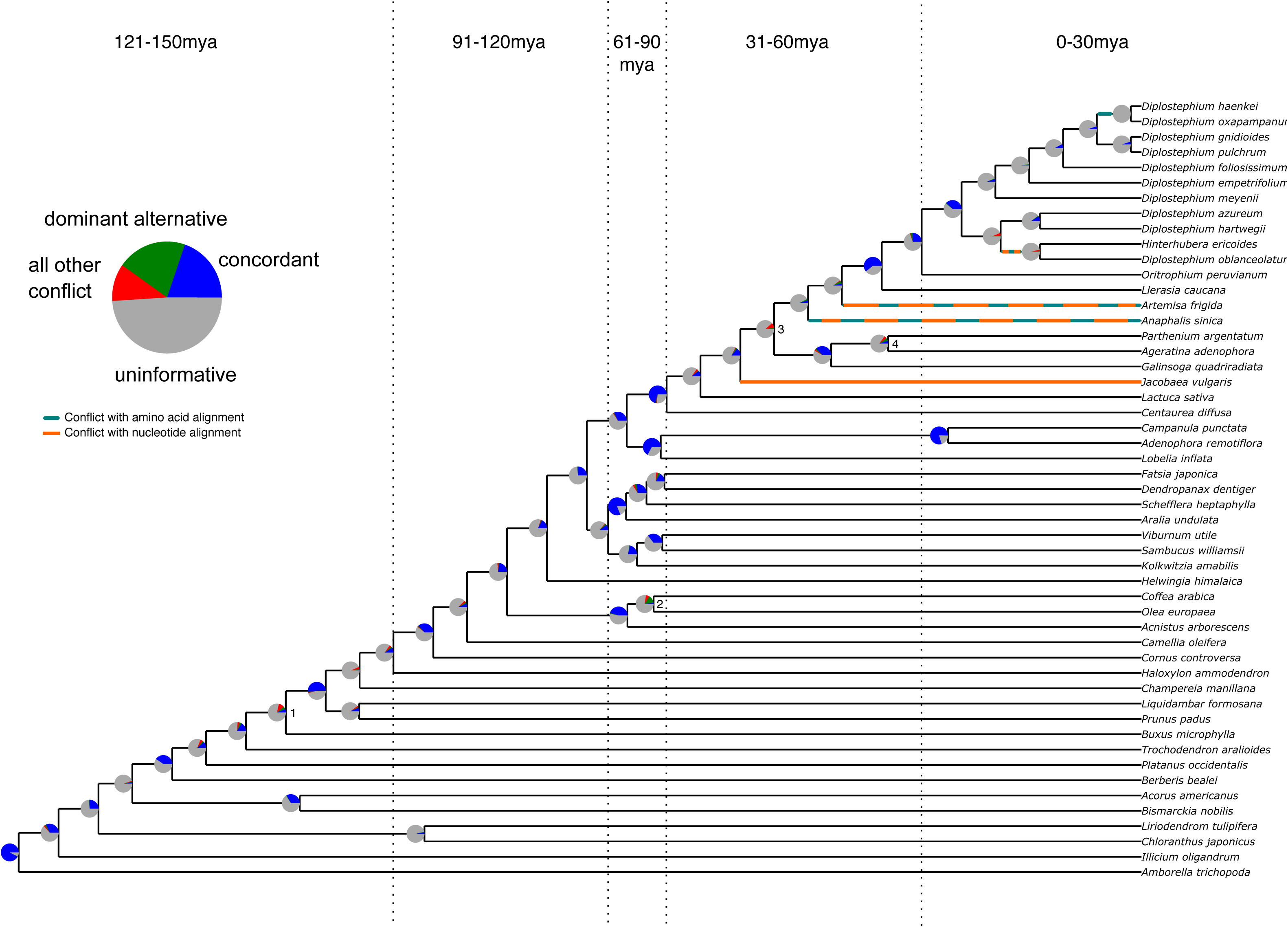
Summary of chloroplast conflict against the reference phylogeny of angiosperms. Green and orange lines indicate where the amino acid- and nucleotide-inferred plastome trees conflict with the reference phylogeny. Pie charts depict the amount of gene tree conflict observed in the nucleotide analysis, with the blue, red, green, and gray slices representing, respectively, the proportion of gene trees concordant, conflicting (supporting a single main alternative topology), conflicting (supporting various alternative topologies), and uninformative (BS < 70 or missing taxon) at each node in the species tree. The dashed lines represent 30 myr time intervals (positioned based on Magallon *et al.* 2015 and Vargas *et al.* 2017) used to bin nodes for examinations of conflict at different levels of divergence.

**Figure 2.**
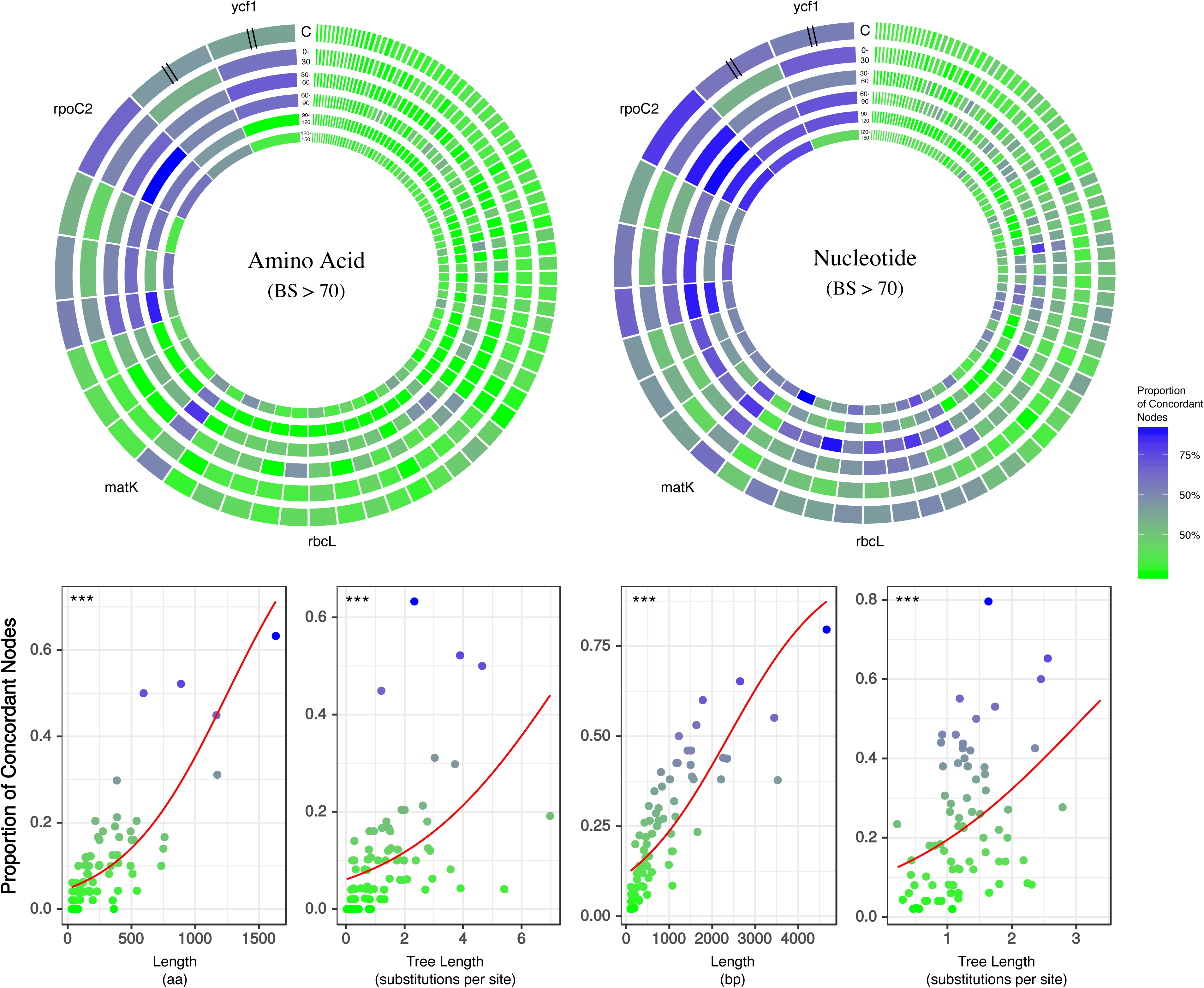
Gene tree concordance/conflict at varying time scales. Each diagram represents a different molecule type and shows the proportion of concordance each gene exhibits at the five time slices shown in Fig. 1: (1) 150−120 mya, (2) 120−90 mya, (3) 90−60 mya, (4) 60−30 mya and (5) 30−0 mya. The individual genes are scaled by length of alignment; however, *ycf1* and *ycf2* are cut to approximately the length of *rpoC2* due to their abnormally long alignments. The plots along the bottom show the relationships between gene concordance levels and various attributes of the genes (e.g., alignment length, tree length).

**Figure 3.**
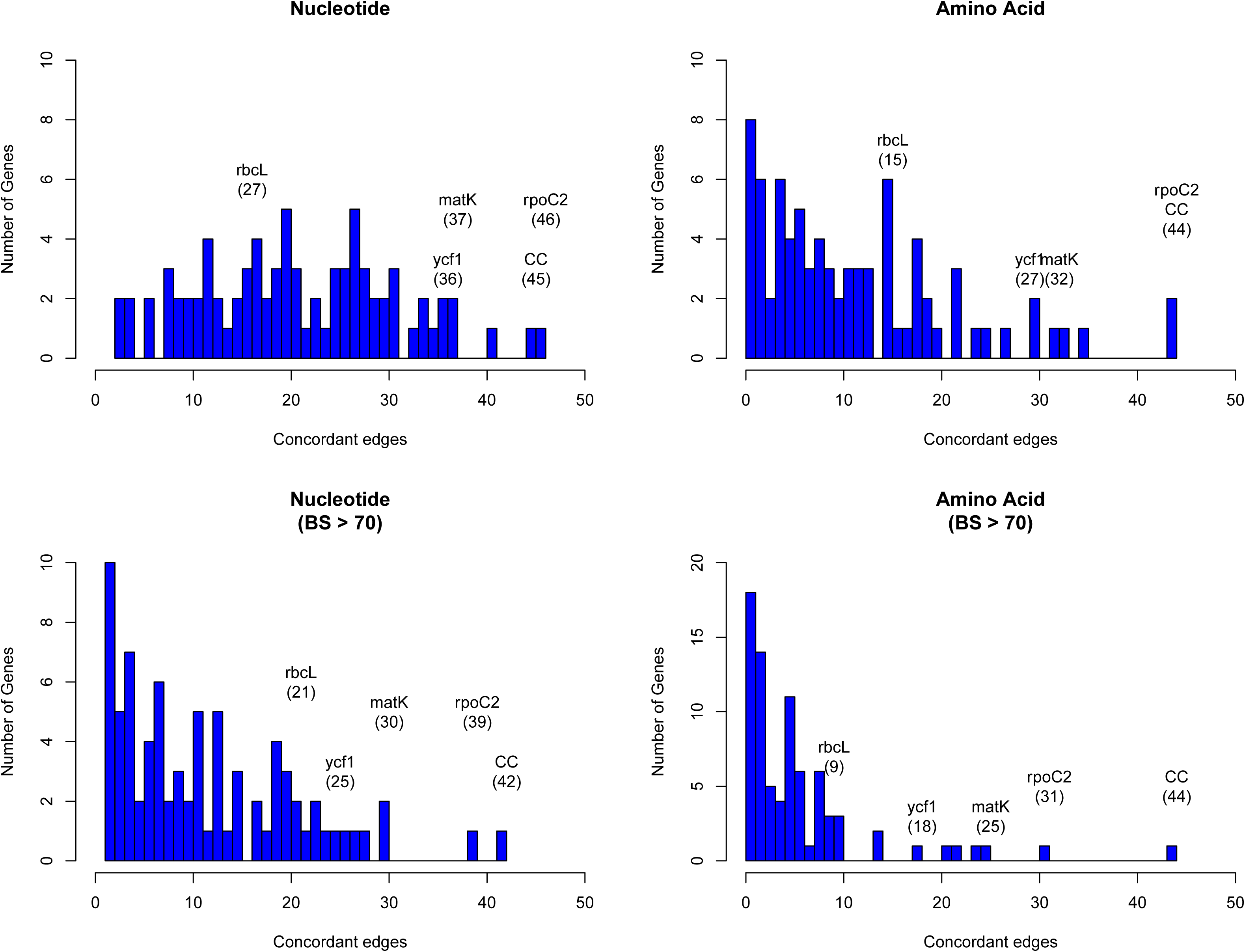
Histograms depicting number of concordant edges each gene tree contains compared to the reference phylogeny (i.e., the ‘accepted tree, AT). Each molecule type is plotted twice (once integrating BS support, shown on the bottom row; another time not integrating BS support, shown on the top row), resulting in four total histograms. The x-axes place each gene based on the number of concordant nodes it shares with the AT; the y-axes show the number of genes with different counts of concordant nodes. Commonly used markers (*matK*, *ndhF*, *rbcL*, and *ycf1*) are labeled on the graph, along with the most concordant gene (*rpoC2*) and the number of concordant nodes for the complete chloroplast (CC) compared to the AT.

### Predictors of Concordance

We examined the statistical relationships between gene tree concordance (using concordance data from the AT-based conflict analysis) and alignment length, tree length, and root-to-tip variance. Because the performance of each gene consists of an aggregate sample of trials (with each node being a trial with outcomes of either concordance or discordance), we analyzed the relationships between gene performance (concordance levels) and alignment length, tree length, and root-to-tip variance using logistic regression of aggregate binomial trials with the function glm() in R (R Core Team, 2018). Binomial models were generally characterized by high residual deviance, and we thus allowed for overdispersion by fitting quasibinomial logistic regressions (using ‘family = quasibinomial()’ in R). All code used for these analyses is available on GitHub (https://github.com/jfwalker/ChloroplastPhylogenomics).

We modelled gene performance as a function of length, tree length, and root-to-tip variation, and as a function of each predictor individually. Because it is possible that apparent relationships between alignment length and concordance may reflect signal from gene information content per alignment site, we also modelled gene performance as a function of length and tree length (as a proxy of gene information content, noted above), to assess the relationship between alignment length and gene performance after controlling for variation associated with gene information content. This has the added benefit of controlling for possible multicollinearity introduced by the covariation between tree length and root-to-tip variation.

Investigation of model fits on full datasets revealed that several observations were highly influential based on leverage and Cook’s distance values. Therefore, we also conducted investigations on reduced datasets to investigate the influence of these observations. In amino acid datasets, we excluded *rpl22* and *rpl32*, which were probably influential due to their high tree length values, and *ycf1* and *ycf2*, which were probably influential based on their long alignments. In nucleotide datasets, we excluded *clpP*, *rps15*, *ycf1* and *ycf2*. Combined analyses of alignment length and tree length were not subject to influence driven by high root-to-tip variance values, and hence reduced datasets had fewer genes removed. In this case, we excluded only *ycf1* and *ycf2.* Regression results were summarized in tables using the R package Stargazer (Hlavac, 2018).

### Saturation Analyses

We also performed saturation analyses on all the chloroplast genes to determine if they were capable of inferring deep divergence times (Phillipe and Forterre, 1999). The saturation analysis was performed on each codon position and thus the data were realigned by codons for this analysis. The amino acid alignments were used to guide the codon alignments using the program pxaa2cdn, from the phyx package. Four of the genes did not appear to match the amino acid sequence in length (i.e., the gene was not 1/3^rd^ the length of the amino acid); thus these were left out of this analysis, since the codons could not be properly aligned. This discrepancy is likely due to errors in the GenBank submission (the amino acid and nucleotide data do not perfectly correspond). Saturation was assessed by determining the observed number of differences between sequences compared to the inferred number of substitutions. This analysis was performed using the “dist.dna” and “dist.corrected” functions in the R package “ape” (Paradis et al. 2017), with the JC69 model of evolution used for the correction. The analysis was conducted on the entire gene and on each codon position separately.

### Comparison of genomic regions

We assessed the utility of the three major plastome regions—the Long Single Copy (LSC) region, Short Single Copy (SSC), and the Inverted Repeat (IR) region—for reconstructing angiosperm phylogeny in two ways. First, we constructed ML phylogenies (as described above for each of the ‘plastome’ analyses) for each genomic region using the concatenated set of genes comprising each region; with the resulting trees, we then calculated (for each genomic region) the number of nodes concordant with the AT. Second, using the concordance levels of each individual gene (described above), we created a plastome diagram (with genes arranged according to their genomic position) showing the concordance levels of each gene at the five different time scales discussed above (Fig. S1); this permits a qualitative visual assessment of the general concordance levels of each genomic region at each time slice.

### Assessment of supported conflict

We compared four contentious regions of the AT to a series of alternative relationships. The tested alternative relationships were present in either the GT (i.e., the plastome phylogeny of Gitzendanner et al., 2018), our plastome phylogenies (with >70 BS for the node in question; we refer to this tree below as the ‘plastome tree’, or PT), and/or three or more of the gene trees from our nucleotide dataset (with >70 BS for the node in question). These alternative relationships pertain to: (1) the placements of *Buxus* and *Trochodendron*; (2) relationships among *Acnistus* (Solanales), *Coffea* (Gentianales), and *Olea* (Lamiales); (3) the placements of *Jacobaea* and *Artemisia* in Asteraceae; and (4) the placements of *Galinsoga*, *Ageratina*, and *Parthenium* in Asteraceae. To test these alternative relationships, we used a modified version of the Maxmimum Gene Wise Edge (MGWE) method (Walker et al. 2018), where instead of a defined “TREE SET”, we used a constraint tree for each alternative hypothesis. The likelihood for every gene was calculated across each constraint, thereby forcing the relationship in question to be the same. Similar to MGWE, this allowed for conflict across the rest of the topology; however, this provides a greater amount of tree space to be explored (similar to Smith et al., 2018). This method creates likelihood scores solely based on each alternative relationship, making it robust to gene tree conflict outside of the node in question. The modification of the MGWE method has been implemented in the EdgeTest.py program of the package PHylogenetic Analysis Into Lineages (https://github.com/jfwalker/PHAIL).

## RESULTS

### Patterns of conflicting chloroplast signal

The sampling for this experiment allowed us to examine conflict at multiple timescales, with roughly 10 divergences occurring in each 30 Ma bin. Our ‘plastome’ trees, inferred from the concatenated gene sets (nucleotide and amino acid), were highly concordant with the AT (Fig. 1). Without considering support, the gene trees showed notable levels of conflict across the different analyses. This pattern is similar to that seen in nuclear phylogenomic datasets (Smith et al., 2015). When considering support, the majority of the plastid genes were uninformative for practically all nodes in the phylogeny (Fig. 1; Tables S2, S3); i.e., they had bootstrap support below 70 (moderate support) for that particular relationship (whether in conflict or concordance).

There was no obvious relationship between the amount of gene tree conflict and evolutionary scale (i.e., conflict was relatively evenly distributed across shallow and deeper nodes/time scales; Figs. 1, 2). Although the greatest degree of gene tree concordance with the AT appeared in the nodes with inferred ages between 90–61 mya (ages based on Magallón et al., 2015), these nodes typically still contained at least 50% uninformative gene trees (Fig. 1). Genome location also should weak correspondence to conflict/concordance (Fig. S1; Table S4); discussed further below. Rather than timescale and genome location, data type (nucleotide vs amino acid) had a much greater impact on the prevalence of conflict (Figs. 2, 3), with the amino acid dataset generally showing higher levels of gene tree conflict. When factoring in support (BS 70 cutoff), the amino acid data set showed even less concordance with the AT (as more genes were considered uninformative due to low BS support); the integration of support also decreased concordance of the nucleotide data set with the AT, but proportionally less. See Figs. S3 and S4 for conflict analyses showing the amino acid and nucleotide gene trees mapped onto the AT (the nucleotide results from Fig. S4 are also shown in Fig. 1).

To identify how many different models of amino acid evolution underlie the genes in the plastome, we tested each gene against the candidate set of amino acid models in IQ-TREE and RAxML. We found that a wide range of evolutionary models best fit our data—rather than just a single model for the entire concatenated set of genes. Many of the models were not designed specifically for plastome data, and cpRev, which was designed for plastome data, was only the best fit for 19 of the 79 genes based upon the IQ-TREE model test (Table S2).

To examine relationships between gene characteristics and levels of concordance/conflict, we calculated the following statistics for each gene: alignment length (a measure of gene information content), tree length (a measure of evolutionary rate), and root-to-tip variance (a measure of variation in evolutionary rate across the tree). This information is presented for each gene across both data types in the supplementary information (Tables S2 and S3). We used logistic multiple regression to test relationships between gene performance and characteristics (Tables S5–S12). We found that alignment length had a significant positive multiplicative relationship with odds of concordance across both datasets (Fig. 2, Tables S5-S8). Tree length and root-to-tip variance had significant positive and negative multiplicative relationships, respectively, in bot datasets (Tables S5, S6). Notably, excluding highly influential observations with outlying predictor values rendered all three predictors significant but did not affect the direction of most relationships Tables S7, S8). In models including only alignment length and tree length, both predictors had significant positive multiplicative relationships with odds of concordance even when the other was included in the model (Fig. 2, Tables S9-S12).

### Genomic patterns of concordance/conflict

In terms of number of nodes concordant with the AT, the LSC and SSC regions were roughly comparable, with both outperforming the IR whether considering bootstrap or not (Table S4). This pattern held across time periods, with the LSC and SSC regions having more concordant nodes than the IR at every time slice. The tree lengths of the LSC and SSC regions (1.82 and 2.05, respectively) were also considerably larger than that of the IR (0.98). The alignment lengths of the LSC region, SSC region, and IR were 73422 bp, 10395 bp, and 19314 bp respectively. The genome diagram of concordance (Fig. S1) does not show any striking patterns among the different genomic regions; however, it is notable that the majority of the LSC region is discordant (or uninformative) with the exception of a few highly informative genes (namely, *rpoC2* and *matK*).

### Performance of individual plastid genes

Across all analyses, *rpoC2* showed the highest levels of concordance with the AT (Fig. 2, Tables S2 and S3); in general, it performed at least as well as the 79 concatenated genes in reconstructing the AT. The commonly used genes *ndhF* and *matK* generally scored among the best-performing plastid genes (in terms of number of concordant nodes), while *rbcL*, the other most commonly used gene, performing relatively poorly (Fig. 2). The *matK* alignment is ~250 bp longer than the *rbcL* alignment; the best performing gene, *rpoC2* (alignment length 4660 bp), is one of the longest plastid genes. However, the notably long region *ycf1*—which encodes for ~5,400 bp (Dong et al., 2015)—did not perform as well as *rpoC2*. In this study, the alignment length of *ycf1* was 21,696 bp (vs. 4660 bp for *rpoC2*). In several cases it performed toward the top; however, it was never the top-performing gene in terms of number of concordant nodes (Tables S2 and S3). Notably, despite the high levels of observed conflict overall, we found that every node of the AT was supported by at least one gene. Thus, to varying degrees, all relationships of the AT are found within the plastome gene tree set.

### Saturation analyses-

None of the genes analyzed showed significant signatures of saturation (Fig. S2). The uncorrected genetic distances and the corrected genetic distances appeared to be roughly the same, and this trend was true for the first, second, and third codon position. Four genes (*ndhD*, *psbL*, *rpl16* and *rpl2*) were unable to be properly analyzed for saturation as the nucleotide data uploaded to GenBank were missing the codons for 2 to 3 amino acids in some taxa resulting in an improper codon alignment.

### Analysis of well-supported conflict

We found that, out of the four major contentious relationships in the AT, the edge-based analysis only supports one as the most likely (Table 1) compared to the alternative relationships found in the other trees examined (i.e., the GT, the plastome trees, and the gene trees). In the case of *Buxus* (node 1 in Fig. 1), this analysis supports *Buxus* as sister to Trochodendraceae (Table 1), which is the topology found in the GT and four gene trees: *rbcL*, *petL*, *rps2*, *rps14* (Table 2). In the case of lamiids relationships (node 2), the data support *Olea* (Lamiales) sister to *Acnistus* (Solanales) + *Coffea* (Gentianales), which is the relationship present in the GT and 15 gene trees (Tables 1, 2). For node 3 (the placements of *Jacobaea* and *Artemisa*), the data support the AT topology, yet no individual gene trees provided >70 BS support for this relationship; however, *rpoC2* matched the GT topology with strong support (Tables 1 and 2). Lastly, for node 4 (the placement of *Galinsoga*), the data collectively support *Parthenium argentatum* sister to *Galinsoga* (our plastome tree, PT), with two gene trees (petA, psaJ) providing strong support for this topology; however, 16 genes (Table 2) supported a dominant alternative (*Ageratina* sister to *Galinsoga quadriradiata*).

**Table 1.** Likelihood scores for alternative resolutions of contentious nodes based on edge-based analyses using a modification of the Maximum Gene Wise Edge (MGWE) method (Walker et al. 2018). The best-supported relationship (highest likelihood score) for each case is presented in **bold**.

| Contentious relationships | GT | AT | CC | Alternative topology (>3 genes supporting) |
| --- | --- | --- | --- | --- |
| *Buxus* placement (node 10) | **-618242.522753** | -618371.605232 | -618371.605232 | **-618242.522753** |
| Lamiid relationships (node 19) | **-618203.494625** | -618235.710231 | **-618203.494625** | **-618203.494625** |
| *Jacobaea* and *Artemisia* placements (node 34) | -618506.967905 | **-618506.967905** | N/A (BS < 70) | N/A |
| *Galinsoga* placement (node 36) | NA | -618308.350169 | **-618174.41765** | -618319.713582 |

**Table 2.** Individual genes supporting (with >70 BS) the alternative topologies examined for each contentious relationship.

| Contentious relationships | GT | AT | CC | Alternative topology (>3 genes supporting) |
| --- | --- | --- | --- | --- |
| *Buxus* placement (node 10) | rbcL, petL, rps2, rps14 | ndhD, rps4, matK, psbB, rpoC2 | ndhD, rps4, matK, psbB, rpoC2 | rbcL, petL, rps2, rps14 |
| Lamiid relationships (node 19) | ndhK, rbcL, rps11, rps7, ycf4, rpoC2, rps4, ndhA, ndhJ, ccsA, rps2, petB, matK, ndhI, atpB | psbB, petA | ndhK, rbcL, rps11, rps7, ycf4, rpoC2, rps4, ndhA, ndhJ, ccsA, rps2, petB, matK, ndhI, atpB | ndhK, rbcL, rps11, rps7, ycf4, rpoC2, rps4, ndhA, ndhJ, ccsA, rps2, petB, matK, ndhI, atpB |
| *Jacobaea* and *Artemisia*<br>placements (node 34) | rpoC2 | None | N/A | N/A |
| *Galinsoga* placement<br>(node 36) | N/A | ndhC, rbcL,<br>matK, ndhE | petA, psaJ | ndhF, petB, psbC,<br>ndhI, psaB, ycf2,<br>psbB, ycf4, petD,<br>rpl16, ndhA, rpoC2<br>atpB, rpoB, ccsA,<br>atpF |

## DISCUSSION

Following the typical assumptions of chloroplast inheritance, we would expect all genes in the plastomes to have the same evolutionary history. We would also expect all plastid genes to show similar patterns of conflict when compared to non-plastid inferred phylogenies. Furthermore, the amino acid and nucleotide plastid data we used should show the same conflicting/concordant relationships against given reference trees (e.g., the AT or GT). However, our results, discussed below, frequently conflict with these common assumptions about chloroplast inheritance and evolutionary history.

### Conflicting topologies inferred from the chloroplast genome

In general, the ‘plastome’ topologies inferred from nucleotide and amino acid alignments showed high levels of concordance with the AT (Fig. 1; Figs. S3 and S4). While the genes within the chloroplast genome are largely uninformative for most nodes of the phylogeny, a number of genes exhibited well-supported conflict (Fig. 1). In general, there appears to be no relationship between evolutionary scale and amount of gene tree conflict: i.e., conflict generally does not appear to correlate with divergence time in angiosperms. Instead, the extent of conflict/concordance had a stronger relationship with the molecule type analyzed. The amino-acid dataset showed the highest levels of gene tree conflict (Figs. 3, S3), and the nucleotide dataset had about half the amount of gene tree conflict found in the amino acid data (Figs. 3, S3, S4). It is difficult to determine the causes of the observed instances of strongly supported conflict, which can be found at most nodes in the phylogeny (Figs. 1, S3, and S4). The edge based analysis’ support for the GT over the PT suggest some systematic error occurred when resolving the species relationships using all genes. However, no analyses we performed fully explain the level of conflict we saw and we suggest that the possibility of biological conflict deserves further exploration, especially given the recent documented instances of inter-plastome recombination (Sancho *et al*., 2018; Sullivan *et al*., 2017) and chloroplast-nuclear genomic exchange (Martin *et al*. 1998; Martin 2003).

The superior performance of (coding) nucleotide data compared to amino acid data possibly stems from the relatively greater information content of nucleotides (i.e., longer alignments). Assuming there is not a significant amount of missing/indel data (with the exception of *ycf1*), longer alignments should result in better-informed models, aided by both parsimony-informative and -uninformative characters (Yang, 1998). However, inherent differences in amino acid and nucleotide models might also explain differences in performance. The nucleotide data was run using the GTR model, where the substitution rates among bases are individually estimated; however, for the empirical amino acid substitution models investigated here, substitution rates are pre-estimated, as the number of estimated parameters for changes among 20 states is extremely large. Additionally, although the plastome has been treated as a single molecule for designing amino acid models of evolution (Adachi *et al.* 2000), a wide variety of amino acid models (some of which were designed for viruses, such as flu or HIV) were inferred to be the best for different plastid genes (Table S2). This might be the result of the different methods implemented in RAxML vs. IQ-TREE for model testing, the different available models, or the lack of sufficient information (because of gene length) to inform the model. However, the most important point is that, based on the amount of information present in each gene, the chloroplast is inferred to evolve under significantly different models of evolution. Given the highly pectinate structure of the AT, phylogenetic inference in this case should rely heavily on the model for the likelihood calculations. While in some cases (e.g., the shortest plastid genes) amounts genetic information might be inherently insufficient, in others, improvements in amino acid modeling might lead to great improvements in phylogenetic inference; such has been suggested for animal mitochondrial data (Richards, 2018).

We expect that at deeper time scales, nucleotides (of coding regions) may begin to experience saturation and thus information loss due to increased noise, at which point amino acids (with 20 states) would begin to outperform nucleotides. However, the time scale of angiosperm evolution does not appear great enough to result in nucleotide saturation (at least for the genes sampled here, given that no genes appeared to exhibit significant levels of saturation; Fig. S2), indicating that nucleotides are the most informative molecule for phylogenetic analysis of plastomes across angiosperms. Future work, with a broader plastome sampling across green plants, will be necessary to determine the evolutionary scale at which amino acids become more informative than nucleotides for phylogenetic inference.

### Analyses of well-supported conflict

The use of a reference topology (AT) afforded us the ability to examine how individual genes within the plastomes agree/conflict with our current, generally accepted hypothesis of angiosperm phylogeny. However, several contentious areas of angiosperm phylogeny remain unresolved, with different topologies recovered across different analyses (e.g., the AT, GT, and PT show small differences in several parts of the tree). Thus we used edge-based analyses to compare these alternative topologies.

The placement of *Buxus* (Buxales) is a salient example, with different analyses resolving different placements. Our edge-based analyses found *Buxus* sister to *Trochodendron*/Trochdendrales (i.e., the GT topology) to have the highest likelihood score (Table 1), and this placement was recovered in four plastid genes (*rbcL*, *petL*, *rps2*, *rps14*) with strong support. However, five plastid genes recovered, with strong support, *Buxus* and *Trochodendron* successively sister to the core eudicots (i.e., the AT topology): *ndhD*, *rps4*, *matK*, *psbB*, and *rpoC2*. Relationships among core lamiid orders are another intriguing example of gene tree conflict. Core lamiid relationships have been notoriously problematic (Refulio-Rodriguez and Olmstead, 2014; Stull et al., 2015), and previous phylogenetic studies have recovered every possible combination of the constituent orders (Boraginales, Gentianales, Lamiales, Solanales). While our sampling does not include Boraginaceae, we tested alternative relationships among the three remaining orders. The relationship (Lamiales(Gentianales,Solanales)), which is present in the GT, received the highest likelihood support and is found in 15 gene trees (Tables 1 and 2). However, the alternatives were each recovered with strong support by two plastid genes: *psaA* and *atpH* (Gentianales sister to the rest), and *psbB* and *petA* (Solanales sister to the rest). Asteraceae also exhibited several examples of strongly supported conflict (Tables 1 and 2), but sampling differences across the trees examined make comparisons more difficult.

These examples above are not meant to represent an exhaustive examination of conflicting relationships within angiosperms. Instead, they are intended to highlight several instances of strongly supported conflict within the plastome. Based on the present analyses, it is difficult to say whether this conflict is biological (e.g.., genes with different evolutionary histories in a single genome) or systematic (e.g., modeling error) in nature. But these results nevertheless have important implications for plastid phylogenomics, suggesting that concatenated analyses should be performed with caution given the notable presence of gene-tree conflict. The majority of nodes show at least one strongly supported conflicting gene, and many nodes have roughly equivalent numbers of conflicting and concordant genes against the reference tree (Fig. 1). Concatenated analyses with extensive underlying conflict can yield problematic results in terms of both topology and branch lengths, and occasionally, even with large datasets, a few genes can drive the entire analysis toward the wrong topology (Brown and Thomson, 2017; Shen et al. 2017; Walker et al. 2018).

### Utility of individual plastid genes for future studies

Previous studies have laid a strong framework for determining the utility of chloroplast regions at various phylogenetic scales. For example, work by Shaw et al. (2005, 2007, 2014) highlighted non-coding DNA regions useful for shallow evolutionary studies, while Graham and Olmstead (2000) explored protein-coding genes useful for reconstructing deep relationships in angiosperms. Here, we expand upon previous work by using a novel phylogenomic approach, allowing us examining the concordance of individual protein-coding plastid genes with all nodes of the accepted angiosperm phylogeny (AT; as well as slight variations thereof, e.g., the GT). We paid special attention to *matK* and *rbcL*, given their historical significance for plant systematics (e.g., Donoghue *et al.*, 1992; Chase *et al.*, 1993; Hilu *et al.*, 2003). We find that *rbcL* performs relatively well in recapitulating the TT—however, *matK* performs considerably better (i.e., it generally has more nodes concordant with the TT; Tables S2 and S3). This is likely due to a strong positive correlation between alignment length and number of concordant nodes, as noted above (Fig. 2); *matK* has a longer alignment/gene length than *rbcL*.

The gene *ycf1* has been found to be a useful marker in phylogenetics (e.g., Neubig *et al.*, 2008; Neubig and Abbot, 2010; Thomson *et al.*, in press) and barcoding (Dong et al. 2015), and here we find that it generally performs above average. The alignment of *ycf1* is abnormally long, and this is likely due to its position spanning the boundary of the IR and the SSC, an area known to fluctuate greatly in size. This variability likely contributes to the value of *ycf1* as a marker for ‘species-level’ phylogenetics and barcoding. However, the performance of *ycf1* does not scale with its alignment length. In terms of concordance, we find that *matK* performs roughly equally as well if not slightly better than *ycf1* (Tables S2 and S3). This might in part be a consequence of *ycf1* being missing/lacking annotation from some species, preventing us from analyzing its concordance/conflict with certain nodes. Nevertheless, our results add to the body of evidence supporting *ycf1* as a generally useful plastid region. However, we found *rpoC2* to outperform all other plastid regions in every case (Tables S2 and S3), and its alignment length (4660 bp) is ~1/5^th^ the length of *ycf1*, easing the computational burden of using alignment tools such as FSA (which would struggle with a region as long as ycf1), as well as divergence dating and tree-building programs such as BEAST (Redelings and Suchard 2006; Drummond and Rambaut 2007).

In our analyses, when BS support is not considered, *rpoC2* performed at least as well if not better than using the concatenation of all chloroplast genes (Tables S2 and S3). When support is considered (Tables S2 and S3), *rpoC2* still remains the best-performing gene, but it performs slightly worse than the concatenation of all chloroplast genes (in terms of number of supported nodes concordant with the TT). The utility of *rpoC2* likely stems from its notable length, resulting in a wealth of useful phylogenetic information. In light of our results, *rpoC2* should be a highly attractive coding region for future studies, as it generally recapitulates the plastome phylogeny while allowing more proper branch length inferences (given that conflicting signal among multiple genes can result in problematic branch length estimates: Mendes & Hahn, 2016). Recent work (Smith et al., 2018) indicates that filtering for a smaller set of highly informative genes might yield more accurate results in various phylogenetic applications (e.g., divergence dating). Given this, *rpoC2* might be particularly useful for comparative analyses requiring accurate branch length estimates (i.e., the majority of comparative analyses). Use of *rpoC2* alone (instead of the entire plastome) would also allow for more complex, computationally expensive models to be implemented. Furthermore, focused sequencing of *rpoC2* would increase compatibility of datasets from different studies, facilitating subsequent comprehensive, synthetic analyses. Although the performance *rpoC2* may be dataset dependent, our results support its utility at multiple levels, in terms of both time scales and sampling. It is important to note, however, that we did include non-coding regions in our study, which would likely outperform coding regions at shallow phylogenetic levels. But, as we noted, it is generally too difficult to use non-coding regions across broad phylogenetic scales (e.g., angiosperms) because of complications related to homology assessment and alignment.

### Genomic patterns of concordance/conflict

Several previous studies (e.g., Jian *et al*., 2008; Moore *et al.*, 2011) have highlighted the Inverted Repeat (IR) as a valuable plastid region for deep-level phylogenetic analyses, attributing its utility to its relatively slow rate of evolution, resulting in less homoplasy and minimal saturation. However, our results contradict this notion, suggesting that the coding sequences of the IR alone perform poorly compared to the LSC and SSC coding regions for reconstructing angiosperm phylogeny (Fig. S1, Table S4). However, there are important differences between our study and earlier studies on the IR (e.g., Jian et al., 2008; Moore et al., 2011). For one, we did not include the ribosomal RNA genes, which are highly conserved; thus if the conserved nature of the IR (or at least portions of it) is the basis of its utility, then this might explain the poor performance in the current study. However, our saturation analyses (described above) did not reveal any genes to have significant saturation issues at the scale of angiosperm evolution. This calls into question the idea that the conserved genes of the IR would make it superior for reconstruction of angiosperm phylogeny. Instead, it is possible that the non-coding regions of the IR (which we did not include here) are highly informative for angiosperm phylogeny. While the non-coding regions of the LSC and SSC regions have been extensively examined for use as phylogenetic markers (Shaw *et al*., 2005, 2007, 2014), the non-coding regions of the IR have been underexplored. Among the IR genes examined here, *ycf2* (which is exceptionally long) showed the greatest levels of concordance (Fig. S1, Table S4), underscoring the idea that longer genes are generally more useful for phylogeny reconstruction.

Another important difference between our study and Moore et al. (2011) is that we only partitioned our data by gene region, while Moore et al. (2011) explored various partitioning strategies (including codon positions and different combinations of genes). It is clear that sequences within the plastomes follow various different models of molecular evolution (as shown above in the results section ***Patterns of conflicting chloroplast signal***). Exploration of more complicated partitioning schemes—which is a time-consuming process (Lanfear *et al*., 2012; Kainer and Lanfear 2015), and beyond the scope of this study—might generally improve plastome-inferred phylogenies.

## Conclusions

We find notable levels of gene-tree conflict within the plastome, at all levels of angiosperm phylogeny, highlighting the necessity of future research into the causes of plastome conflict—do some genes share different evolutionary histories, or is systematic error (e.g., poor modeling) the source of the observed conflict?—and the appropriateness of concatenating plastid genes for phylogeney reconstruction. However, while conflict is evident, the majority of plastid genes are largely uninformative for angiosperm phylogeny (Fig. 1). We find alignment length and molecular type (nucleotide vs amino acids) to be the strongest correlate with plastid gene performance—i.e., longer genes and nucleotides (which have greater information content than amino acids) generally show higher levels of concordance with our accepted hypothesis of angiosperm phylogeny. Accordingly, we found *rpoC2* (one of the longest plastid genes) to be the best-performing plastid gene, outperforming all of the more commonly used genes (e.g., *rbcL*, *ndhF*, *matK*) at reconstructing angiosperm phylogeny. This supports the notion that longer genes (with greater information content) are generally superior for phylogeny reconstruction (Yang, 1998) and should be prioritized when developing phylogenetic studies.

## ACKNOWLEDGEMENTS

We thank S. A. Smith and M. J. Moore for reading an early draft of the manuscript and proving helpful suggestions toward its improvement. JFW was supported by a Rackham Predoctoral Fellowship. G.W.S. was supported by a NSF Postdoctoral Fellowship (NSF DBI grant 1612032). O.M.V was supported by NSF FESD 1338694 and DEB 1240869. D.A.L. was supported by NSF DEB grant 1458466.

## AUTHOR CONTRIBUTIONS

This study was conceived by all of the authors. Analyses were led by J.F.W and N.W.-H, with contributions from D.A.L, O.M.V, and G.W.S. Writing was led by G.W.S., J.F.W., and N.W.-H., with contributions from O.M.V and. D.A.L.

