## Supplementary figures and images for "Characterizing gene tree conflict in plastome-inferred phylogenies"

### Fig. S1

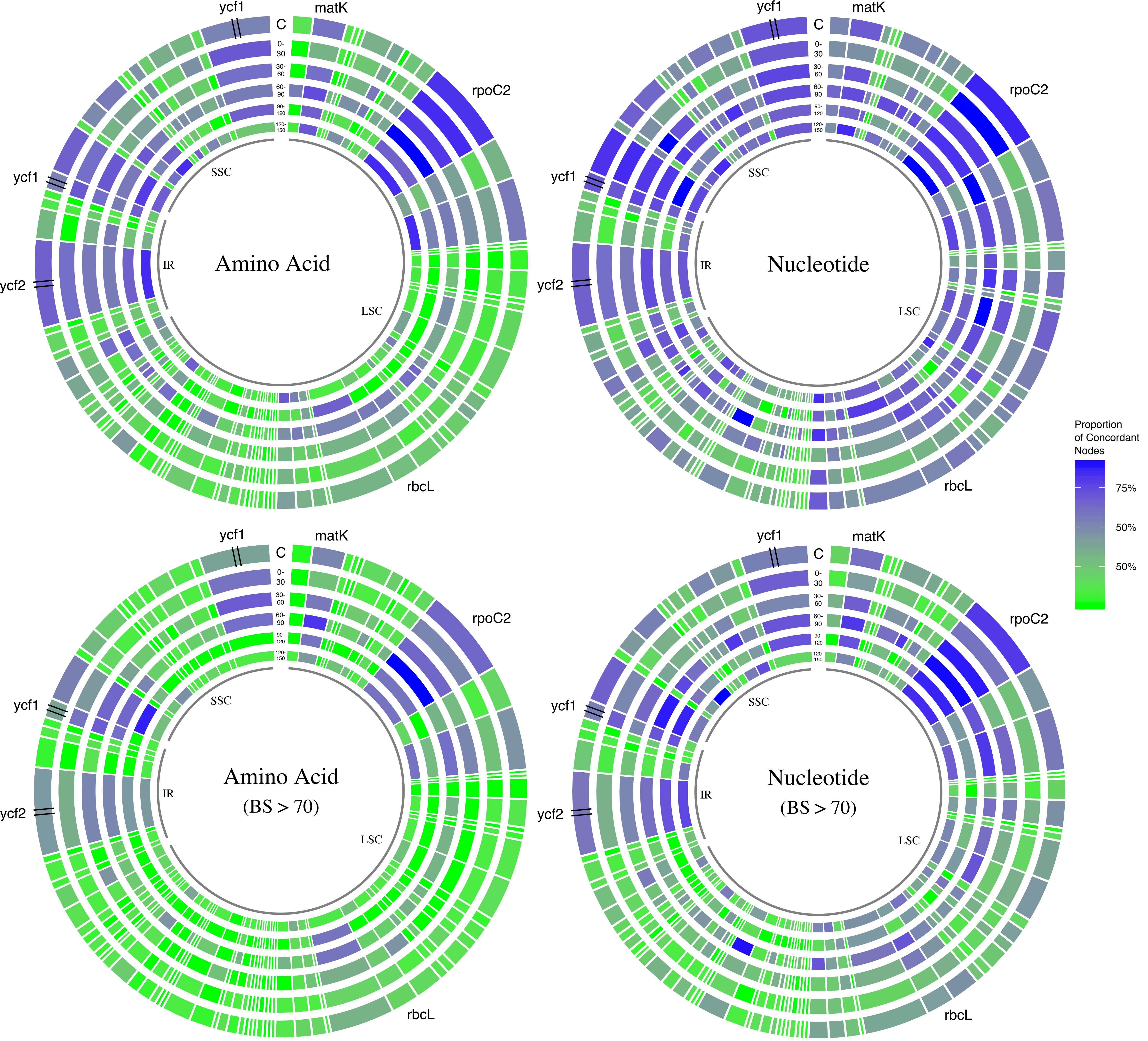

### Fig. S3

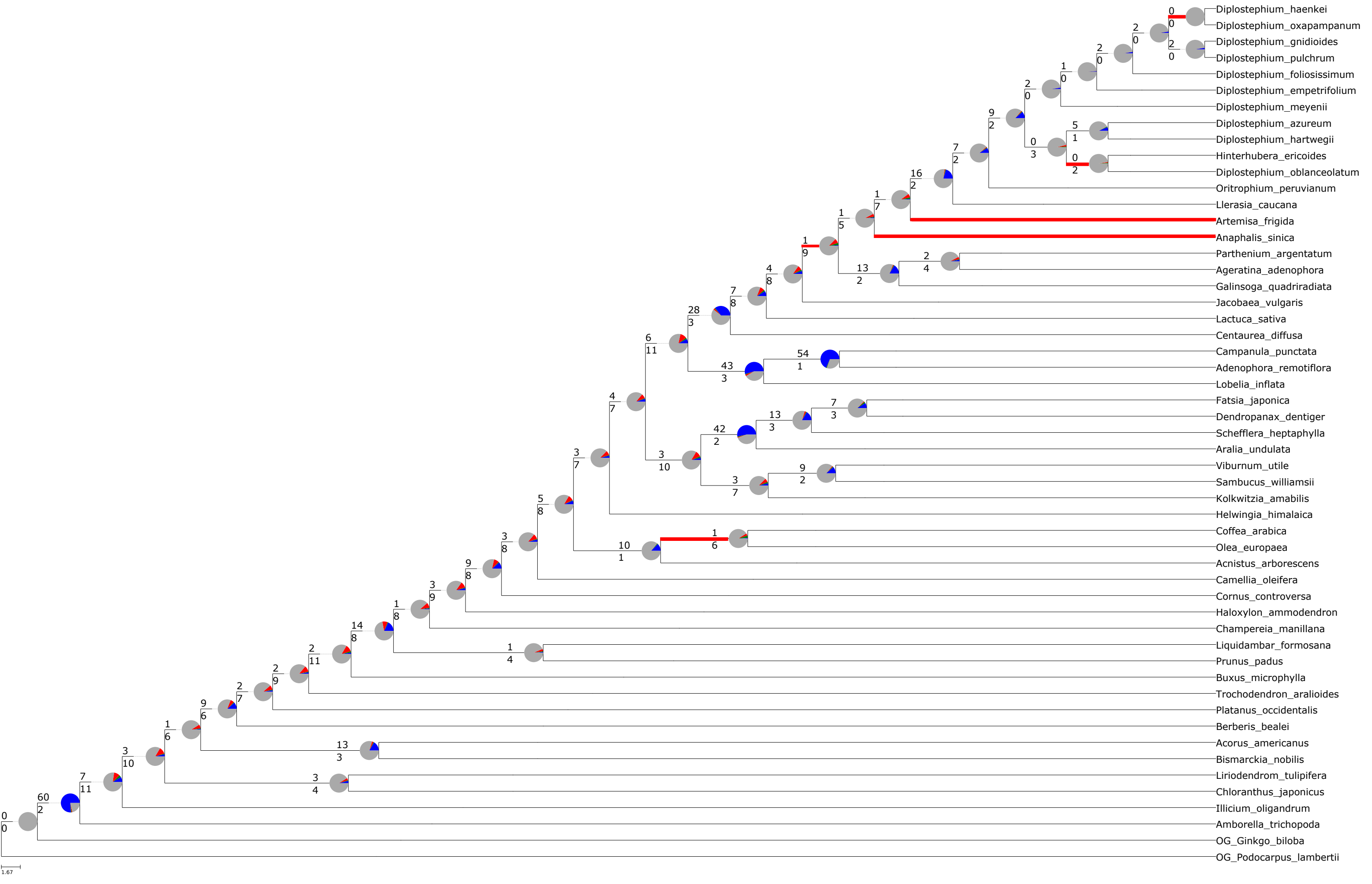

### Fig. S4

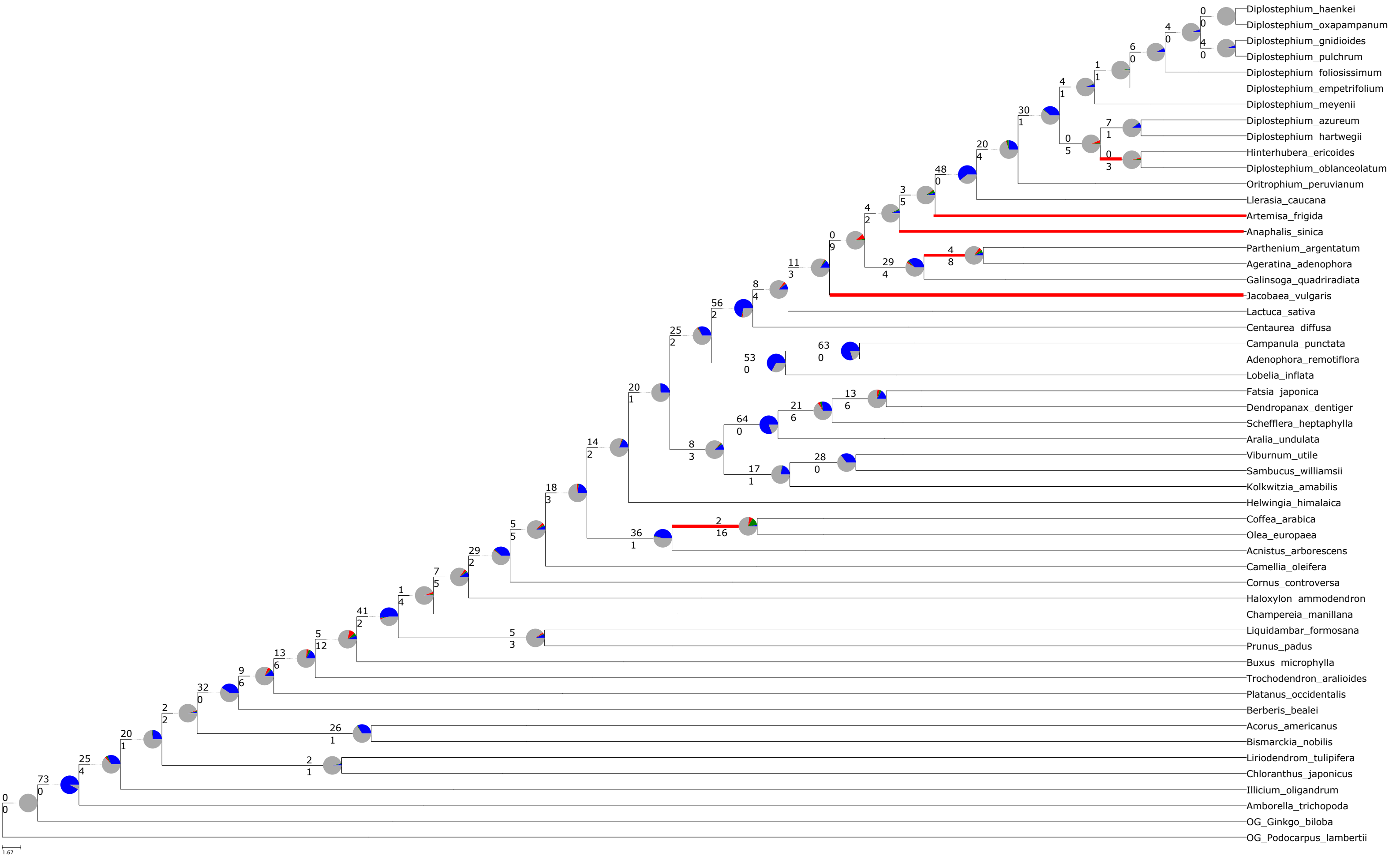
