## Supplementary material for "Characterizing gene tree conflict in plastome-inferred phylogenies": Fig. S2

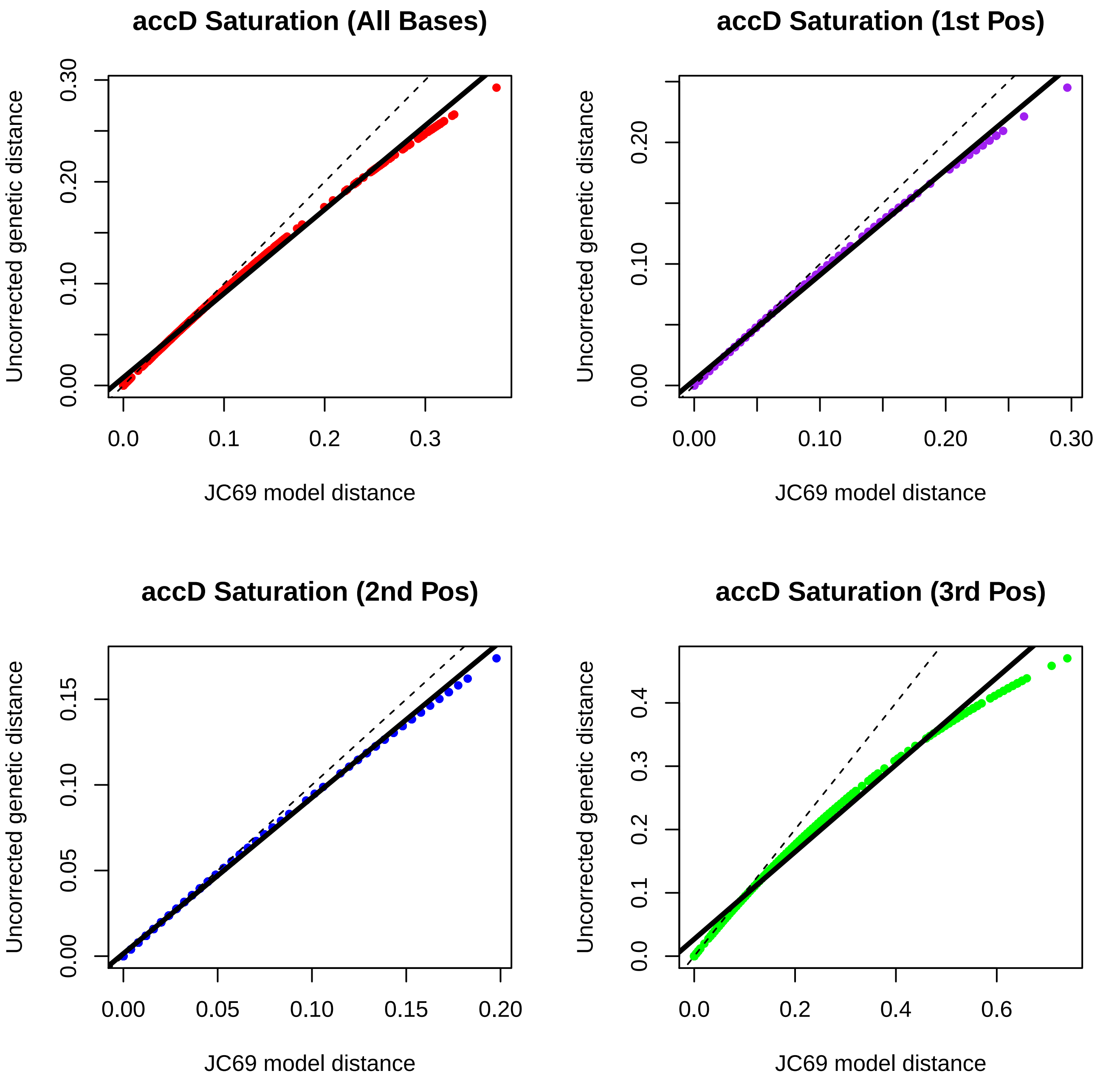
accD Saturation (All Bases)
Uncorrected genetic distance
0.00
0.10
0.20
0.30
0.0
0.1
0.2
0.3
JC69 model distance
accD Saturation (1st Pos)
Uncorrected genetic distance
0.00
0.10
0.20
0.00
0.10
0.20
0.30
JC69 model distance
accD Saturation (2nd Pos)
Uncorrected genetic distance
0.00
0.05
0.10
0.15
0.00
0.05
0.10
0.15
0.20
JC69 model distance
accD Saturation (3rd Pos)
Uncorrected genetic distance
0.0
0.1
0.2
0.3
0.4
0.0
0.2
0.4
0.6
JC69 model distance

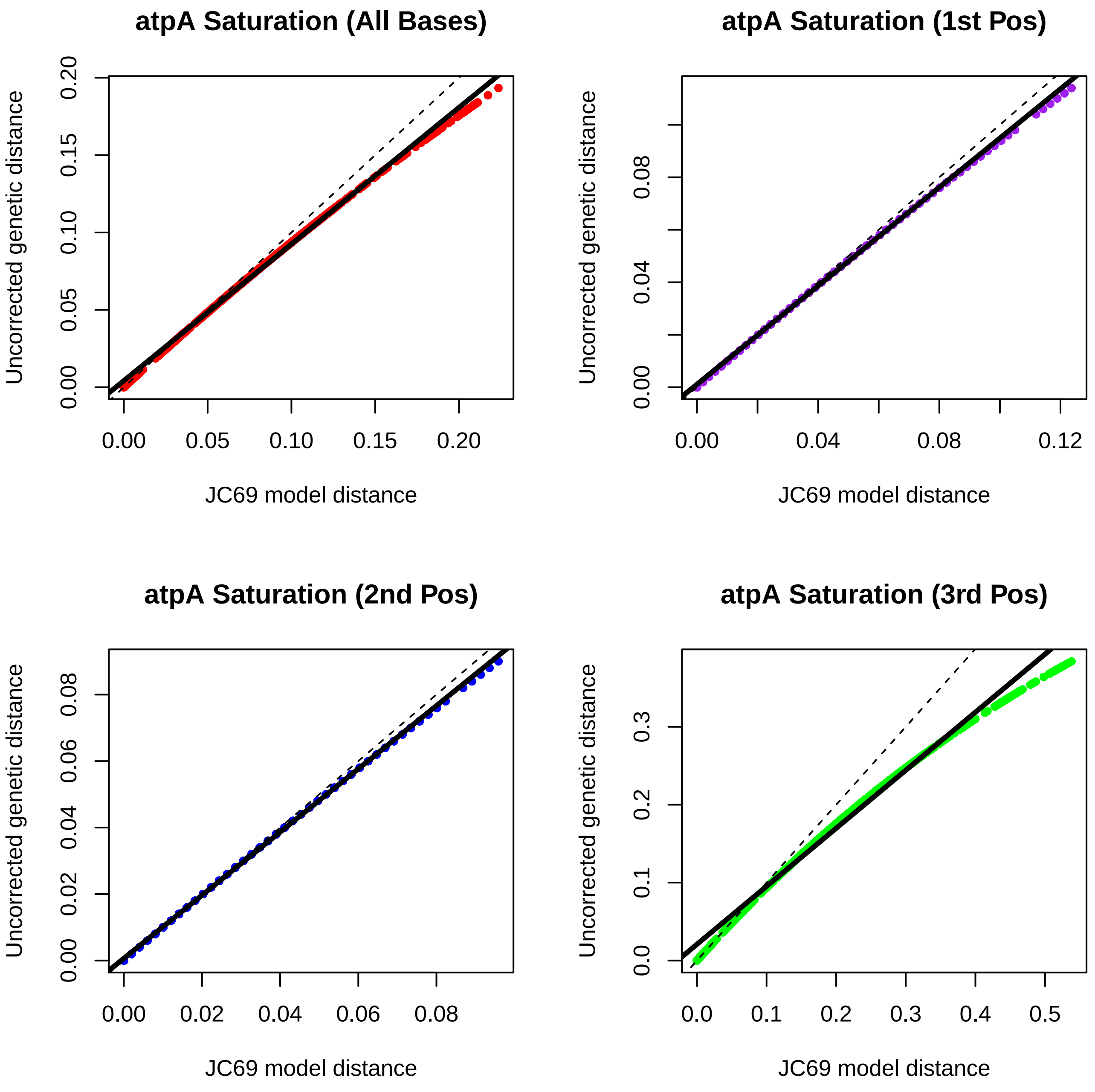
atpA Saturation (All Bases)
Uncorrected genetic distance
0.00
0.05
0.10
0.15
0.20
JC69 model distance
0.00
0.05
0.10
0.15
0.20
atpA Saturation (1st Pos)
Uncorrected genetic distance
0.00
0.04
0.08
JC69 model distance
0.00
0.04
0.08
0.12
atpA Saturation (2nd Pos)
Uncorrected genetic distance
0.00
0.02
0.04
0.06
0.08
JC69 model distance
0.00
0.02
0.04
0.06
0.08
atpA Saturation (3rd Pos)
Uncorrected genetic distance
0.0
0.1
0.2
0.3
JC69 model distance
0.0
0.1
0.2
0.3
0.4
0.5

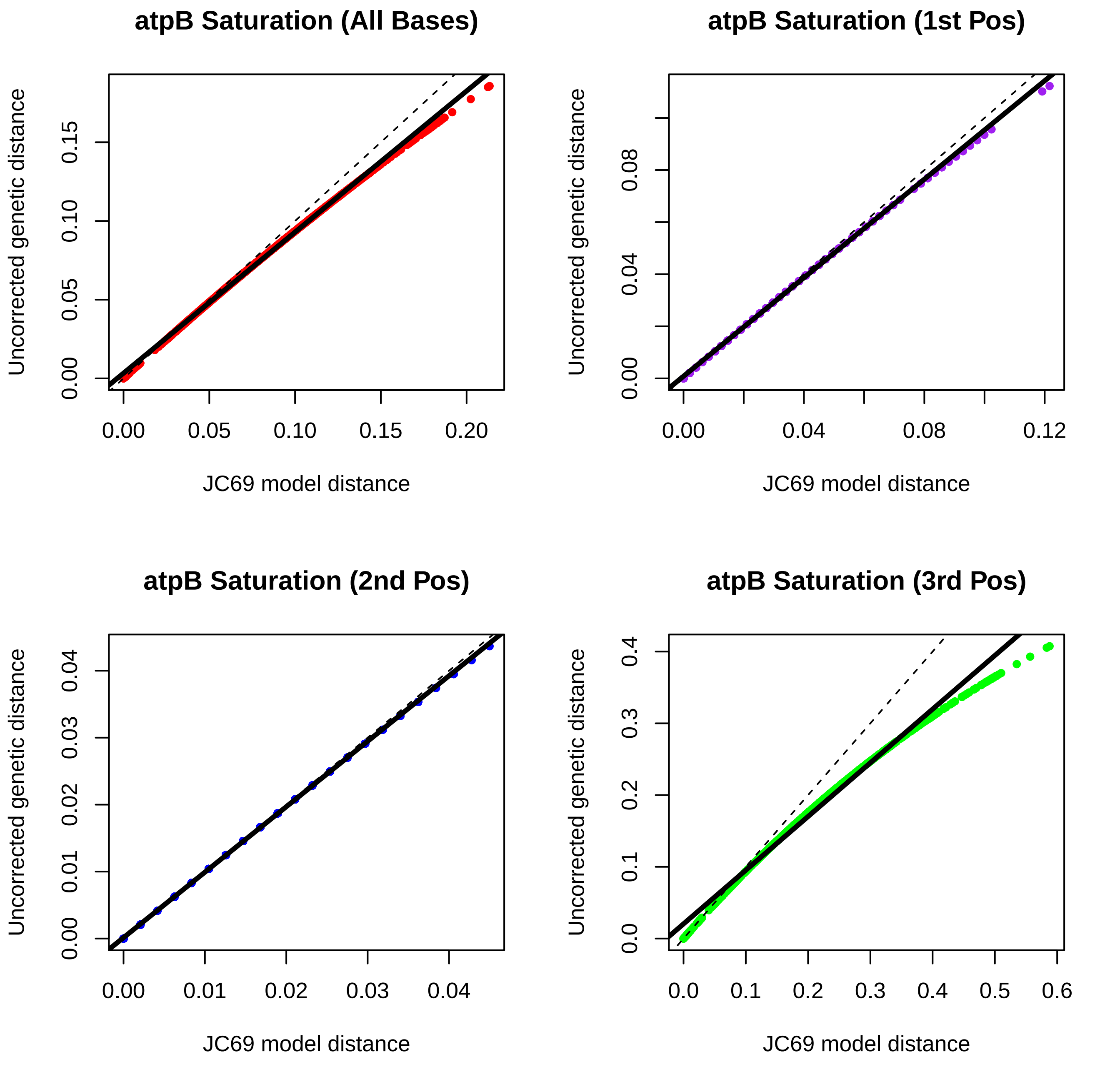
atpB Saturation (All Bases)
atpB Saturation (1st Pos)
atpB Saturation (2nd Pos)
atpB Saturation (3rd Pos)
Uncorrected genetic distance
JC69 model distance
0.00
0.05
0.10
0.15
0.20
0.04
0.08
0.12
0.01
0.02
0.03
0.0
0.1
0.2
0.3
0.4
0.5
0.6

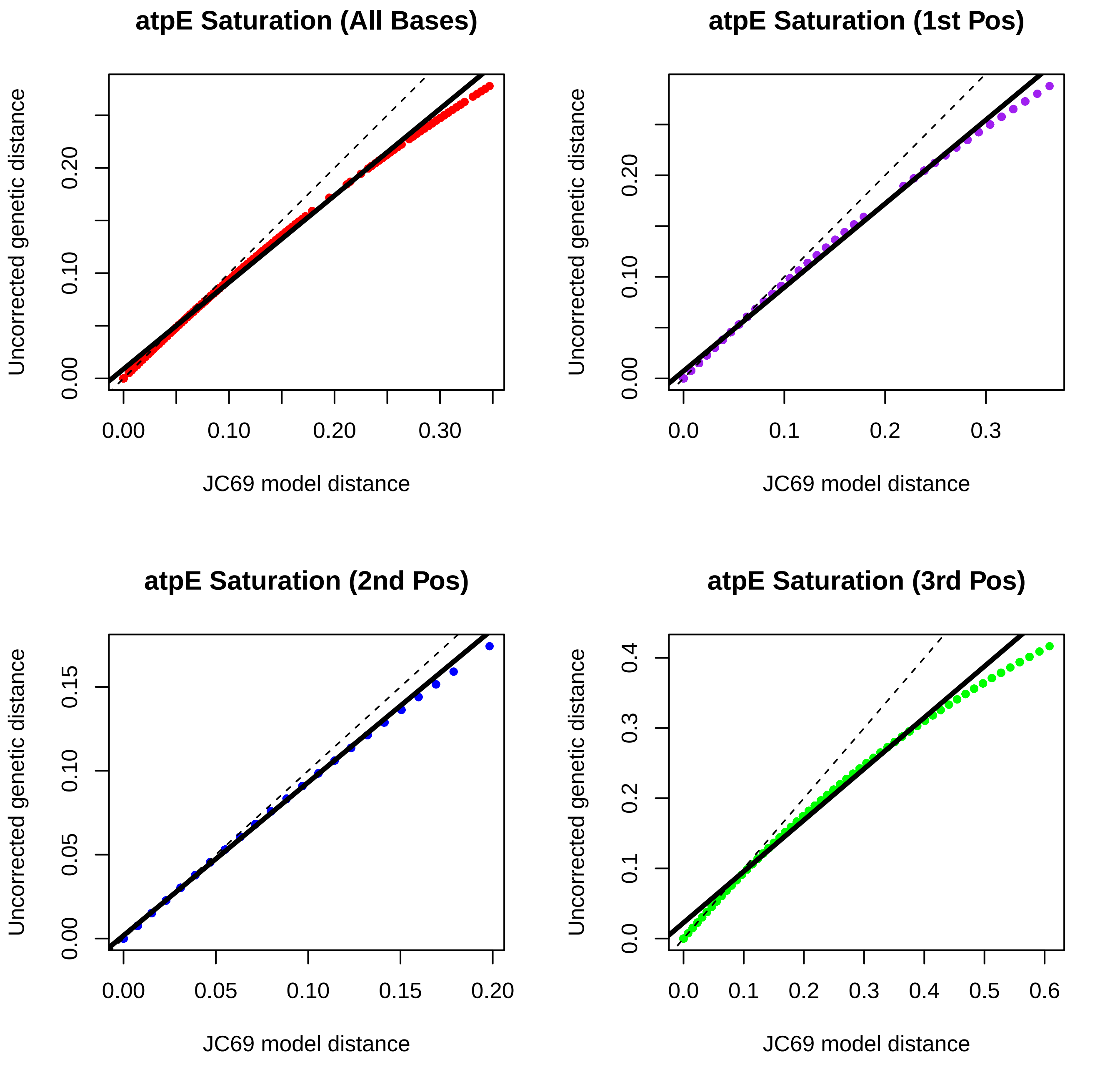

atpE Saturation (All Bases)
atpE Saturation (1st Pos)
atpE Saturation (2nd Pos)
atpE Saturation (3rd Pos)
Uncorrected genetic distance
JC69 model distance
0.00
0.10
0.20
0.30
0.0
0.1
0.2
0.3
0.05
0.15
0.4
0.5
0.6

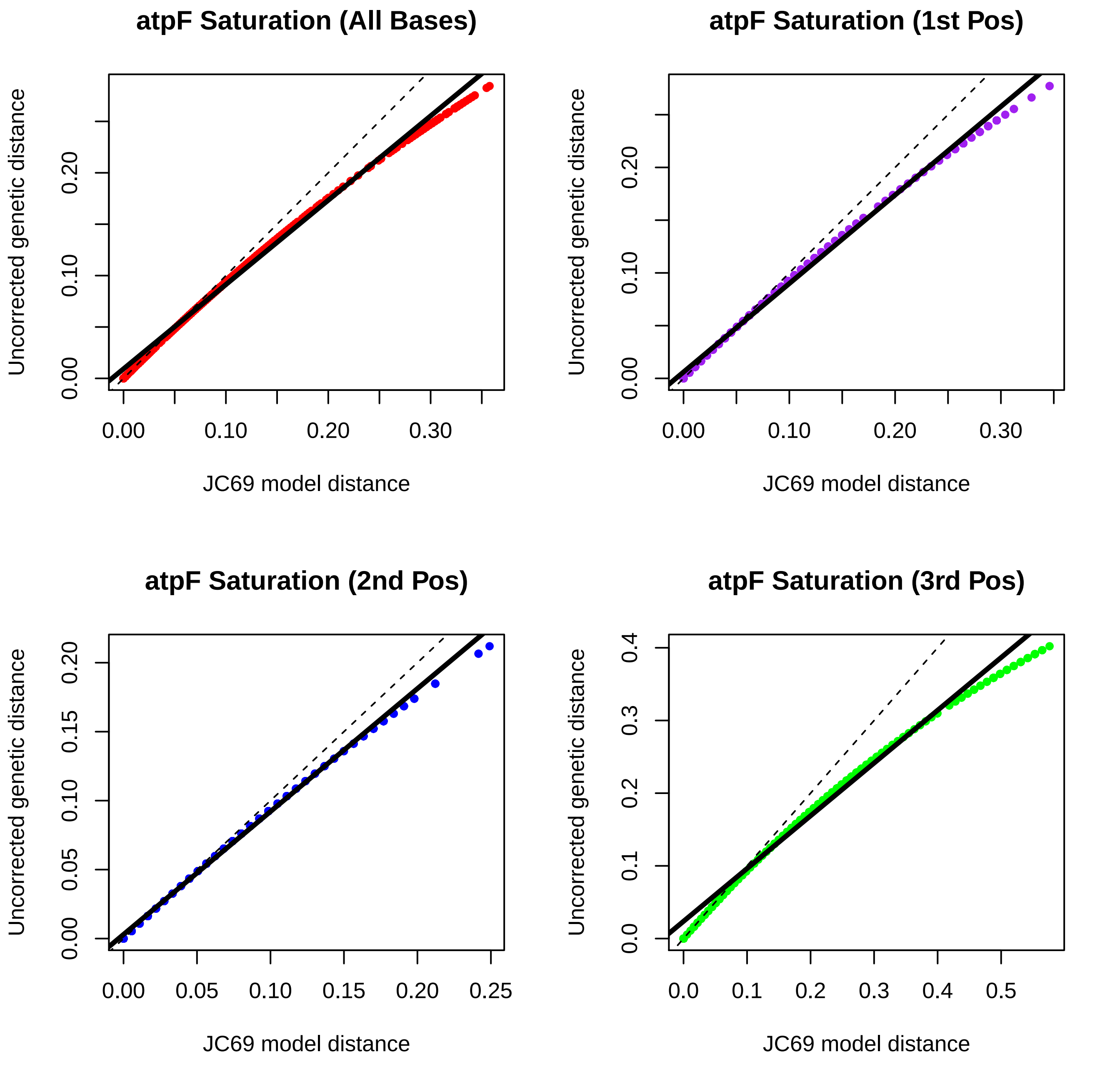

atpF Saturation (All Bases)
atpF Saturation (1st Pos)
atpF Saturation (2nd Pos)
atpF Saturation (3rd Pos)
Uncorrected genetic distance
JC69 model distance
0.00
0.10
0.20
0.30
0.05
0.15
0.25
0.0
0.1
0.2
0.3
0.4
0.5

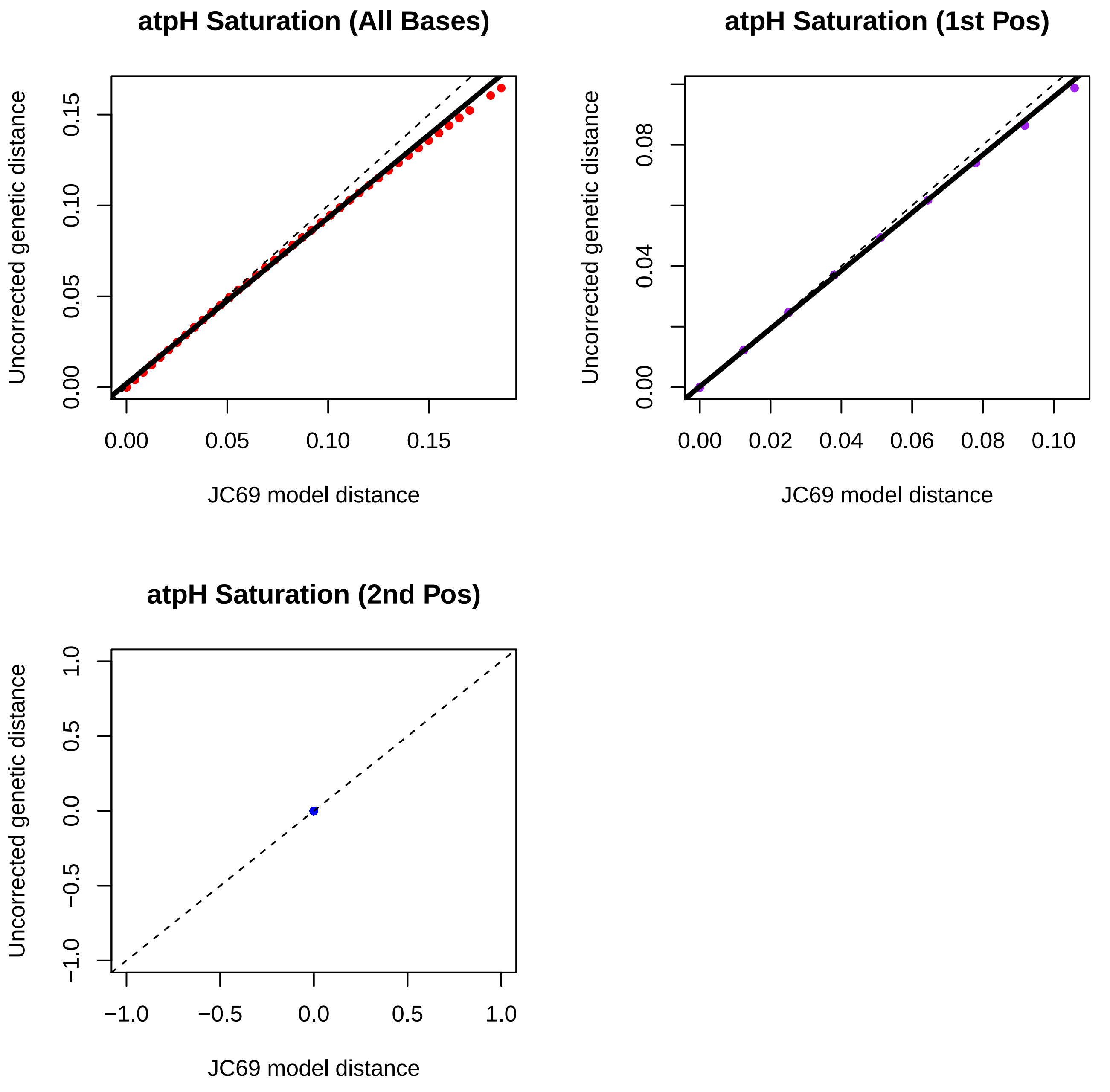
atpH Saturation (All Bases)
Uncorrected genetic distance
JC69 model distance
0.00
0.05
0.10
0.15
atpH Saturation (1st Pos)
Uncorrected genetic distance
JC69 model distance
0.00
0.04
0.08
0.00
0.02
0.04
0.06
0.08
0.10
atpH Saturation (2nd Pos)
Uncorrected genetic distance
JC69 model distance
−1.0
−0.5
0.0
0.5
1.0

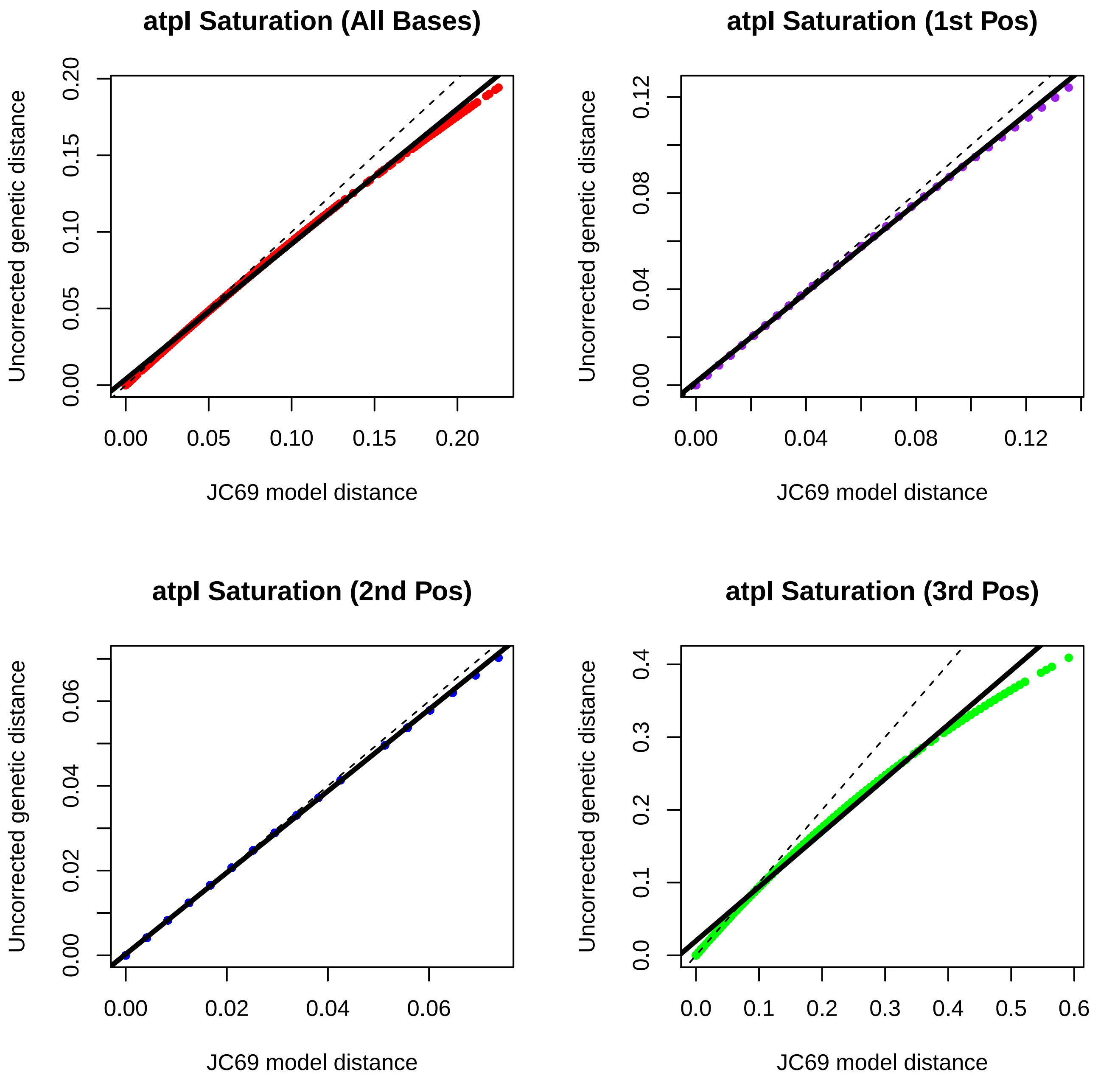
atpI Saturation (All Bases)
Uncorrected genetic distance
0.00
0.05
0.10
0.15
0.20
JC69 model distance
0.00
0.05
0.10
0.15
0.20
atpI Saturation (1st Pos)
Uncorrected genetic distance
0.00
0.04
0.08
0.12
JC69 model distance
0.00
0.04
0.08
0.12
atpI Saturation (2nd Pos)
Uncorrected genetic distance
0.00
0.02
0.04
0.06
JC69 model distance
0.00
0.02
0.04
0.06
atpI Saturation (3rd Pos)
Uncorrected genetic distance
0.0
0.1
0.2
0.3
0.4
JC69 model distance
0.0
0.1
0.2
0.3
0.4
0.5
0.6

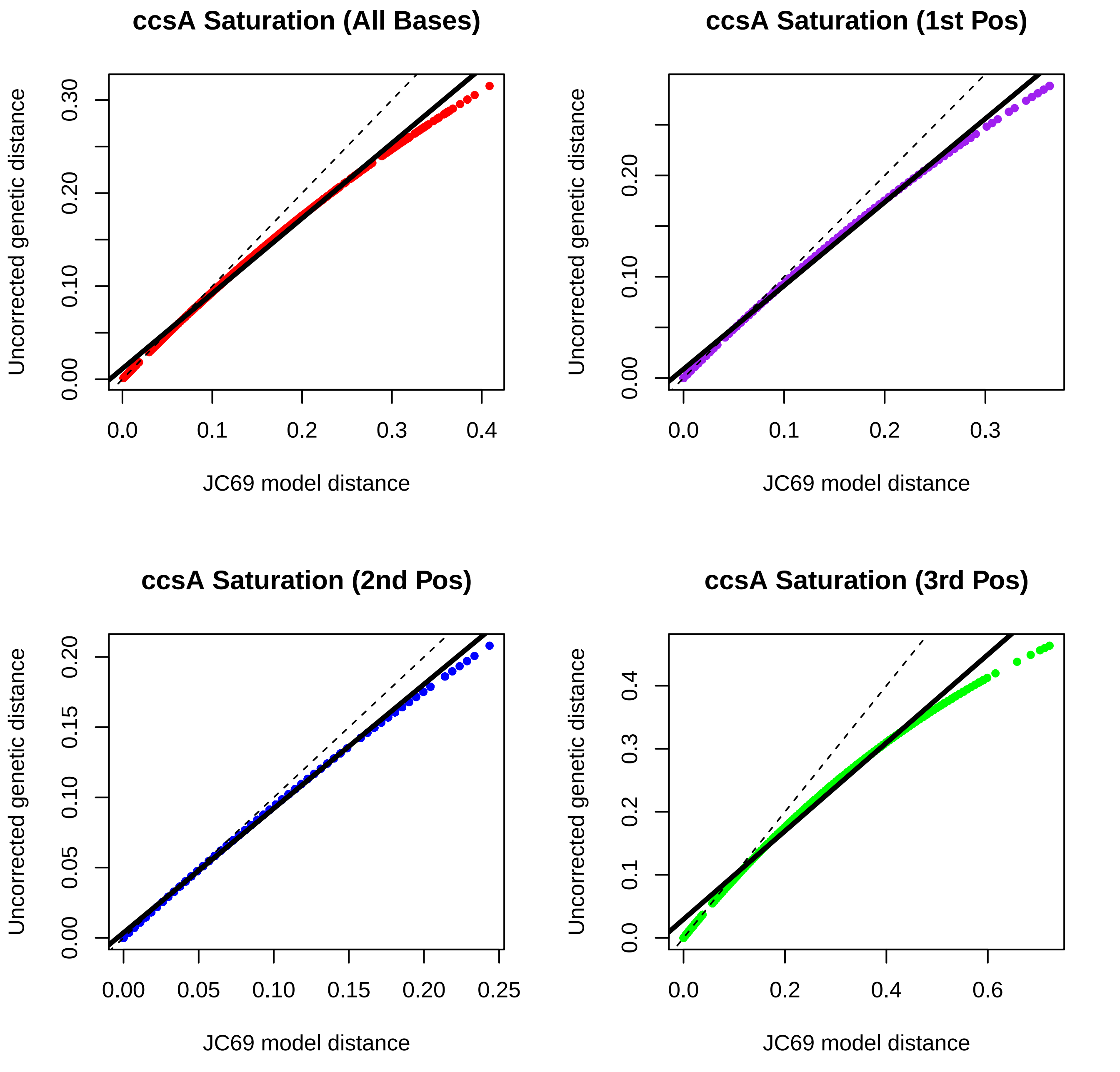
ccsA Saturation (All Bases)
ccsA Saturation (1st Pos)
ccsA Saturation (2nd Pos)
ccsA Saturation (3rd Pos)
Uncorrected genetic distance
JC69 model distance
0.00
0.10
0.20
0.30
0.0
0.1
0.2
0.3
0.4
0.05
0.15
0.25
0.6

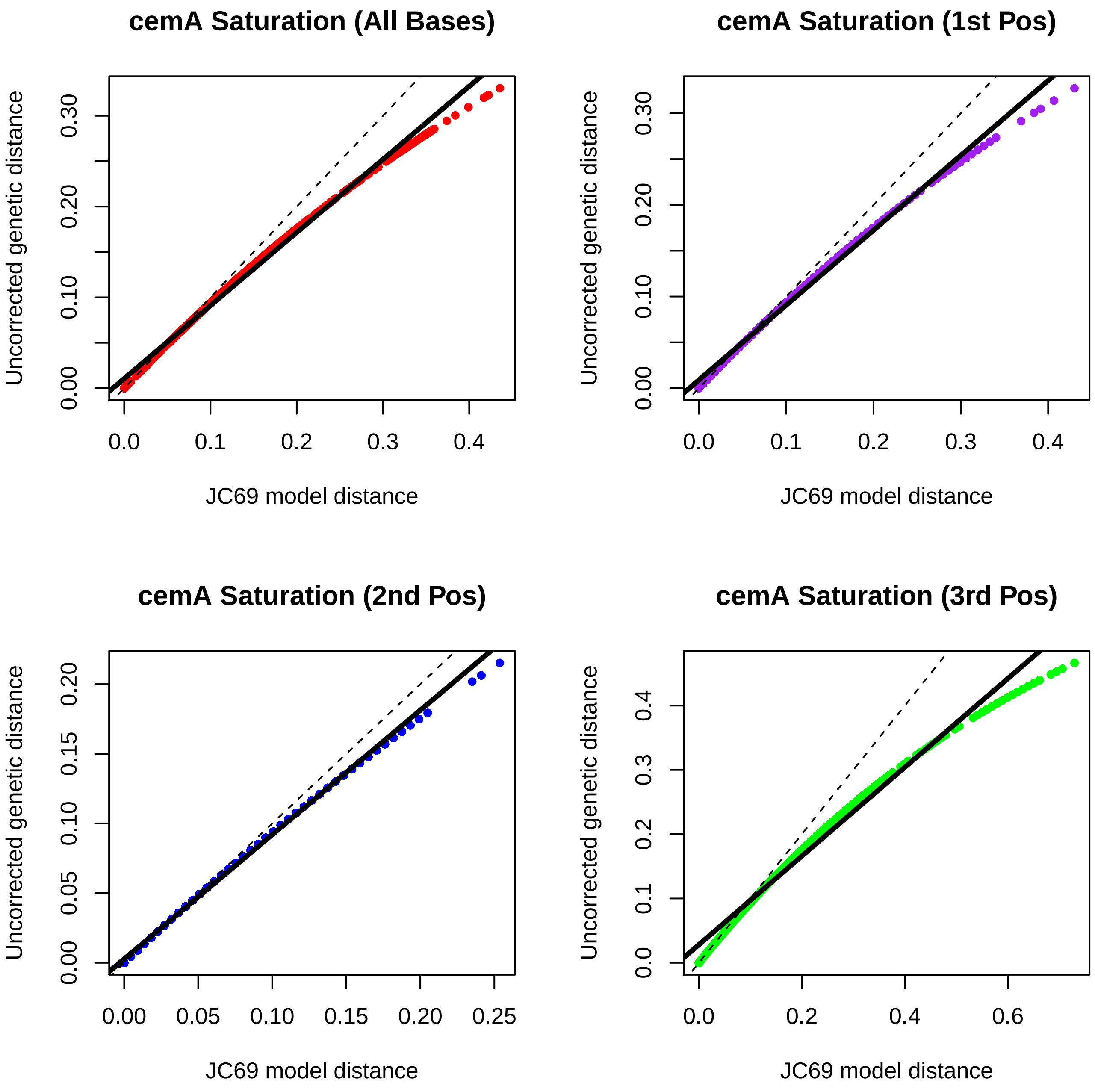

cemA Saturation (All Bases)
cemA Saturation (1st Pos)
cemA Saturation (2nd Pos)
cemA Saturation (3rd Pos)
Uncorrected genetic distance
JC69 model distance
0.00
0.10
0.20
0.30
0.0
0.1
0.2
0.3
0.4
0.05
0.15
0.25
0.6

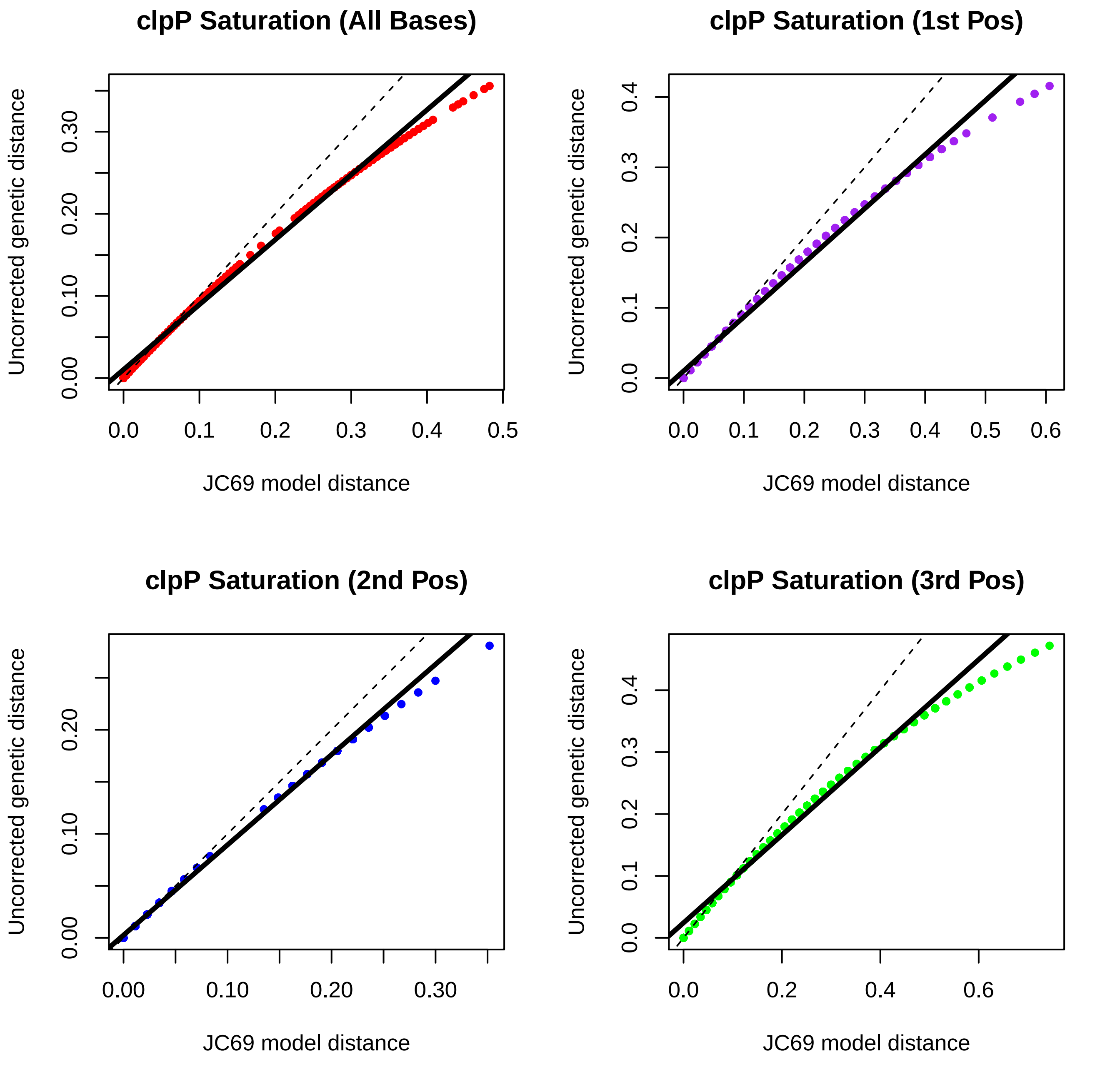
clpP Saturation (All Bases)
Uncorrected genetic distance
0.00
0.10
0.20
0.30
0.0
0.1
0.2
0.3
0.4
0.5
JC69 model distance
clpP Saturation (1st Pos)
Uncorrected genetic distance
0.0
0.1
0.2
0.3
0.4
0.0
0.1
0.2
0.3
0.4
0.5
0.6
JC69 model distance
clpP Saturation (2nd Pos)
Uncorrected genetic distance
0.00
0.10
0.20
0.00
0.10
0.20
0.30
JC69 model distance
clpP Saturation (3rd Pos)
Uncorrected genetic distance
0.0
0.1
0.2
0.3
0.4
0.0
0.2
0.4
0.6
JC69 model distance

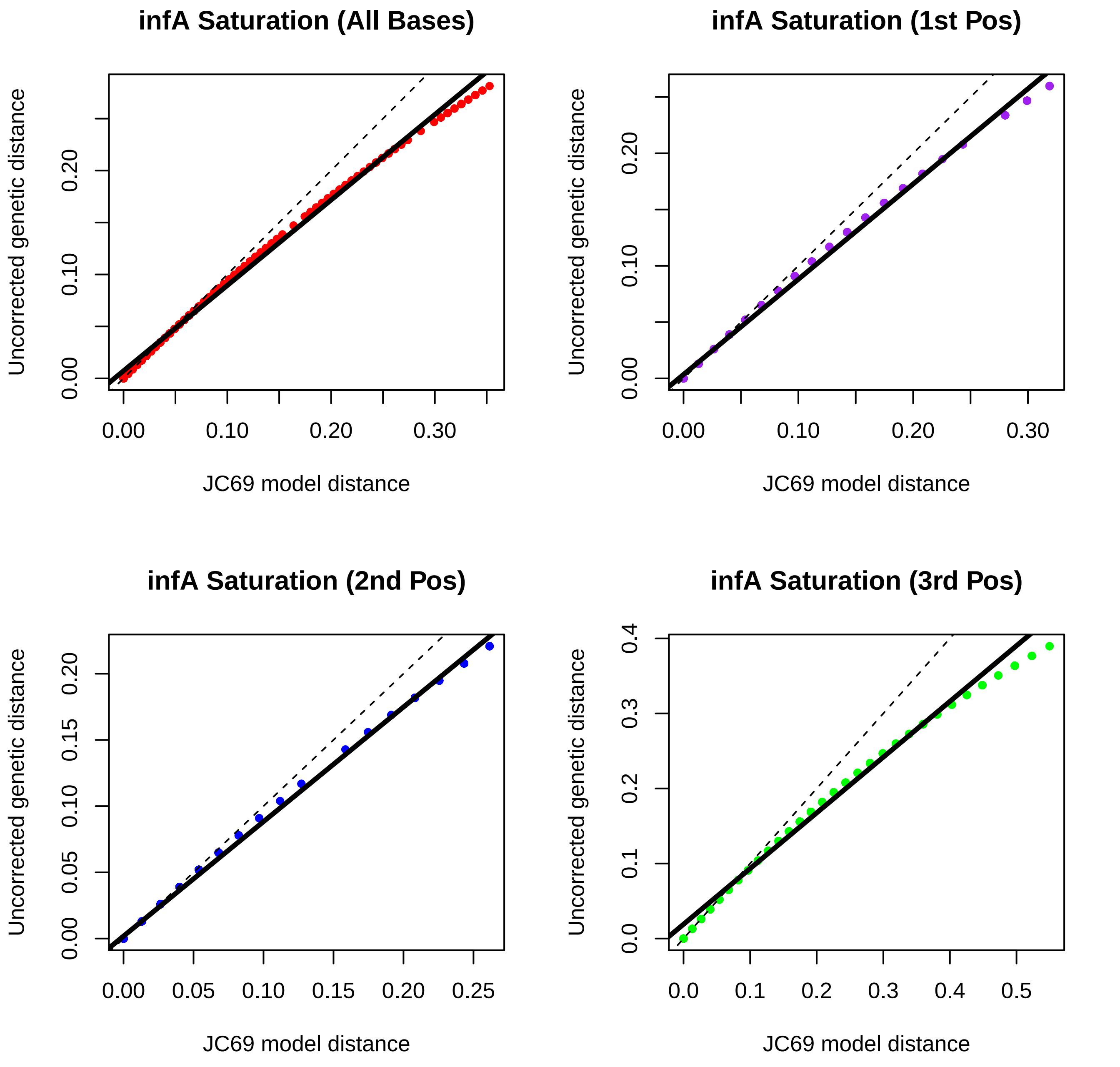
infA Saturation (All Bases)
infA Saturation (1st Pos)
infA Saturation (2nd Pos)
infA Saturation (3rd Pos)
Uncorrected genetic distance
JC69 model distance
0.00
0.10
0.20
0.30
0.05
0.15
0.25
0.0
0.1
0.2
0.3
0.4
0.5

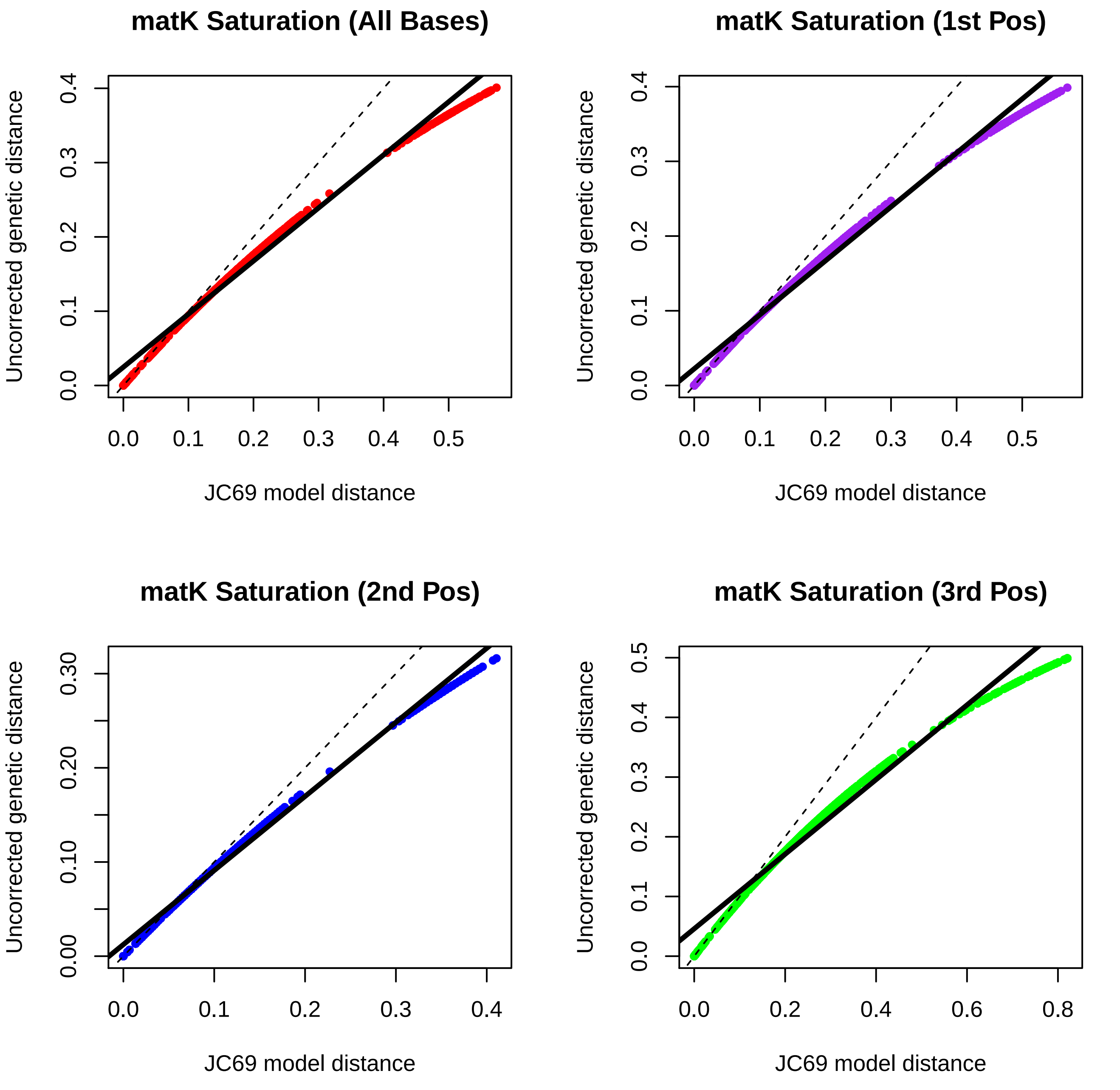

matK Saturation (All Bases)
matK Saturation (1st Pos)
matK Saturation (2nd Pos)
matK Saturation (3rd Pos)
Uncorrected genetic distance
JC69 model distance
0.0
0.1
0.2
0.3
0.4
0.5
0.00
0.10
0.20
0.30
0.6
0.8

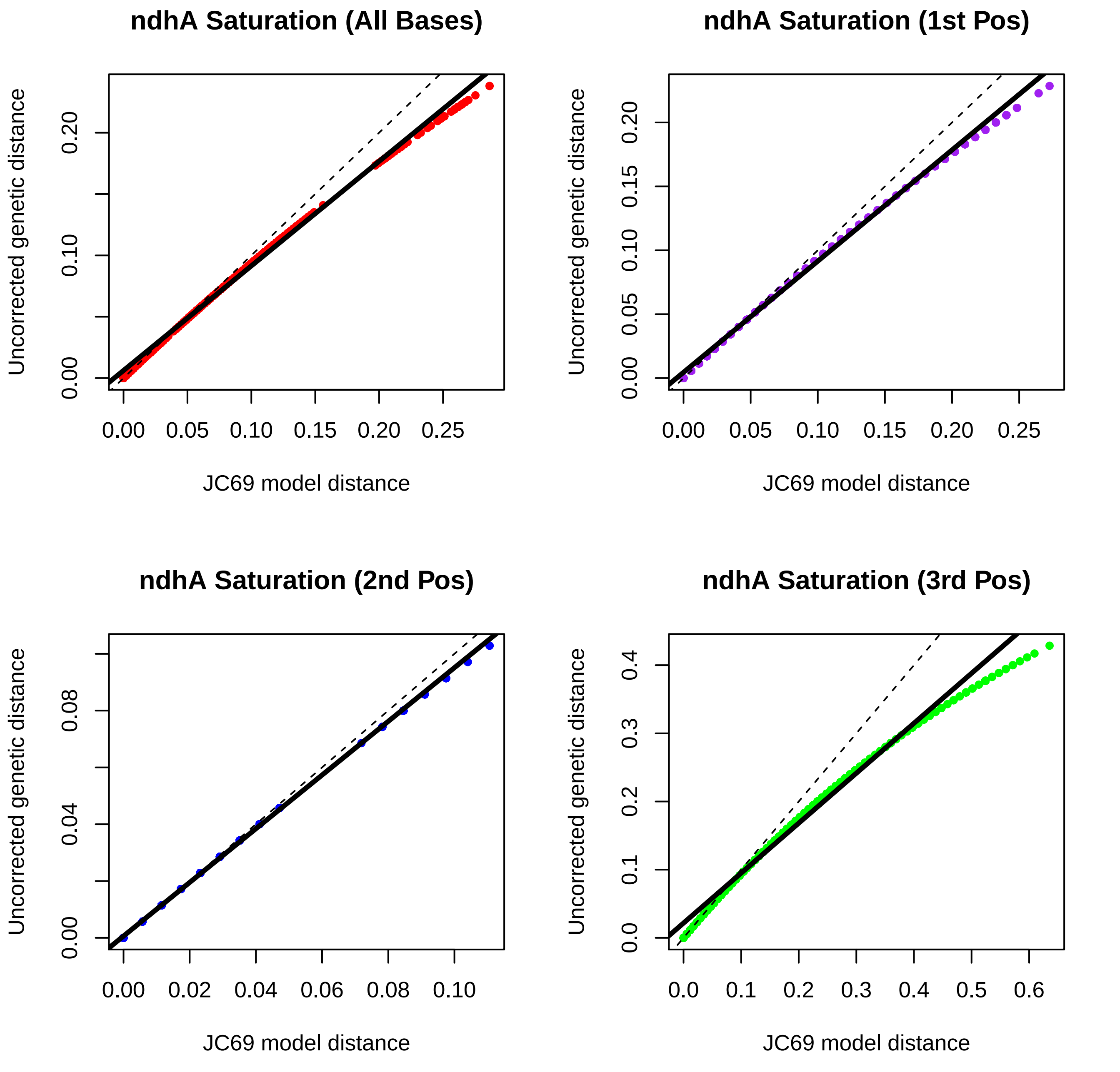

ndhA Saturation (All Bases)
ndhA Saturation (1st Pos)
ndhA Saturation (2nd Pos)
ndhA Saturation (3rd Pos)
Uncorrected genetic distance
JC69 model distance
0.00
0.05
0.10
0.15
0.20
0.25
0.02
0.04
0.06
0.08
0.0
0.1
0.2
0.3
0.4
0.5
0.6

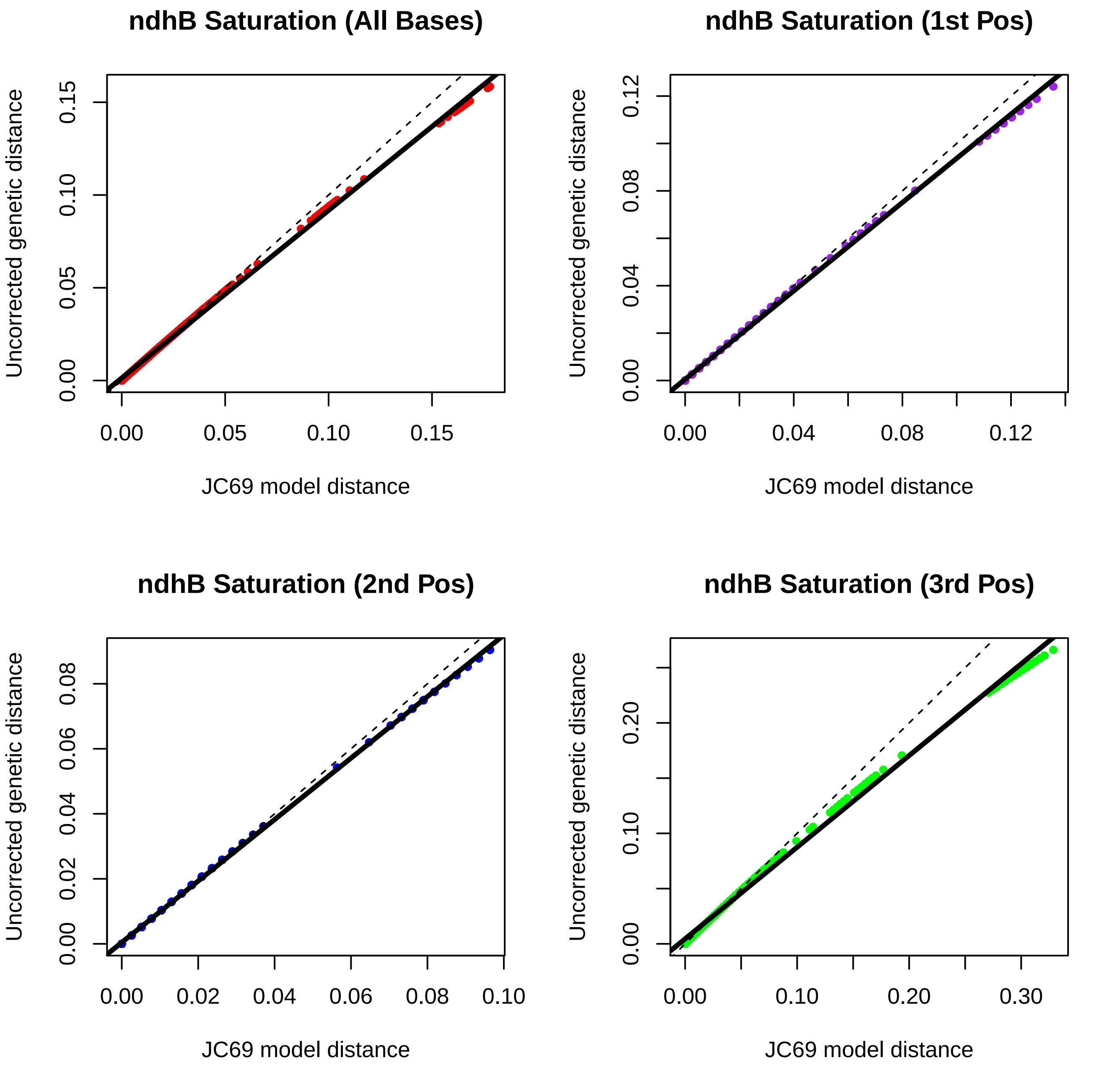
ndhB Saturation (All Bases)
ndhB Saturation (1st Pos)
ndhB Saturation (2nd Pos)
ndhB Saturation (3rd Pos)
Uncorrected genetic distance
JC69 model distance
0.00
0.05
0.10
0.15
0.04
0.08
0.12
0.02
0.06
0.20
0.30

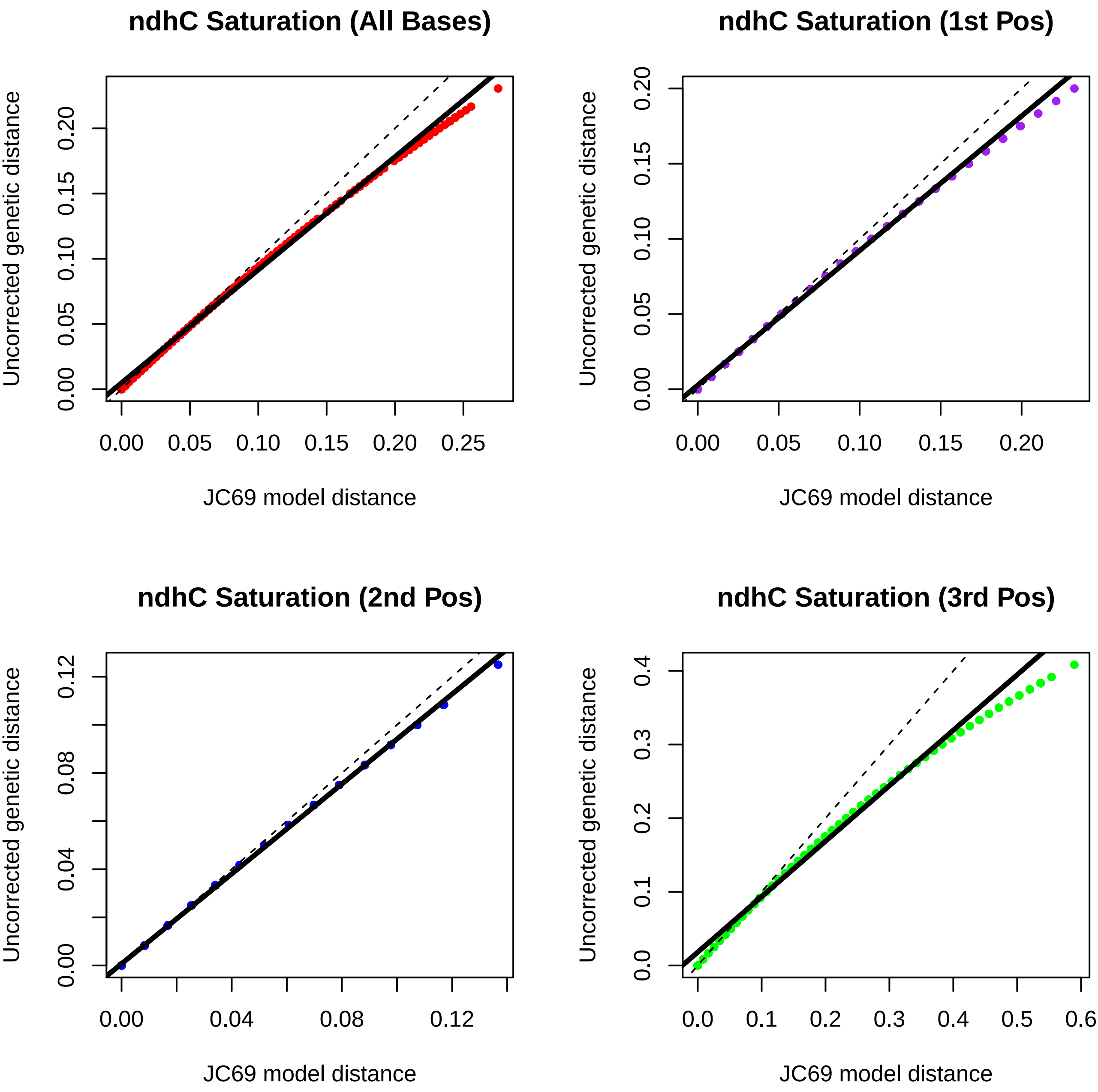
ndhC Saturation (All Bases)
ndhC Saturation (1st Pos)
ndhC Saturation (2nd Pos)
ndhC Saturation (3rd Pos)
Uncorrected genetic distance
JC69 model distance
0.00
0.05
0.10
0.15
0.20
0.25
0.04
0.08
0.12
0.0
0.1
0.2
0.3
0.4
0.5
0.6

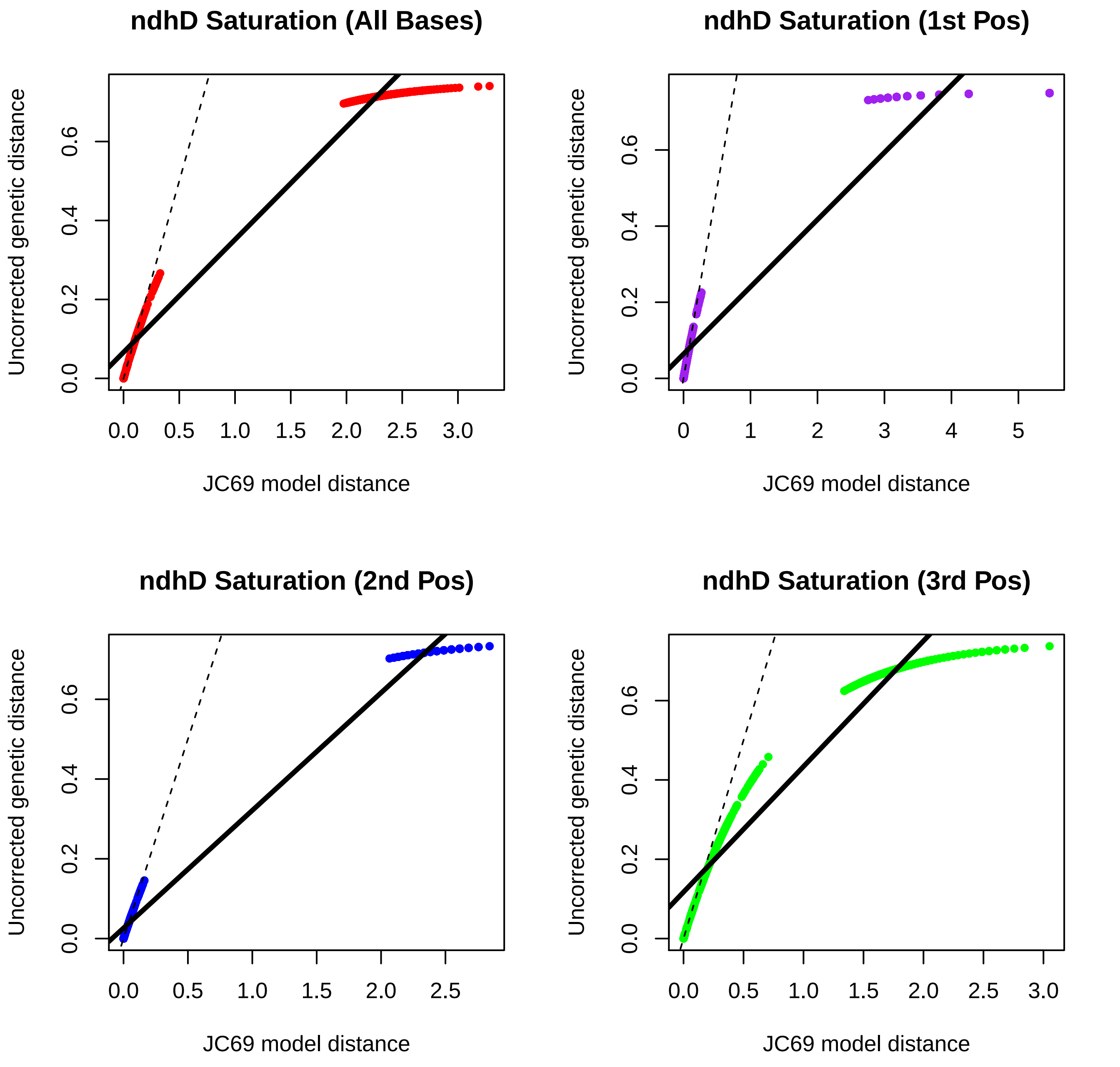

ndhD Saturation (All Bases)
ndhD Saturation (1st Pos)
ndhD Saturation (2nd Pos)
ndhD Saturation (3rd Pos)
Uncorrected genetic distance
JC69 model distance
0.0
0.2
0.4
0.6
0.5
1.0
1.5
2.0
2.5
3.0
0
1
2
3
4
5

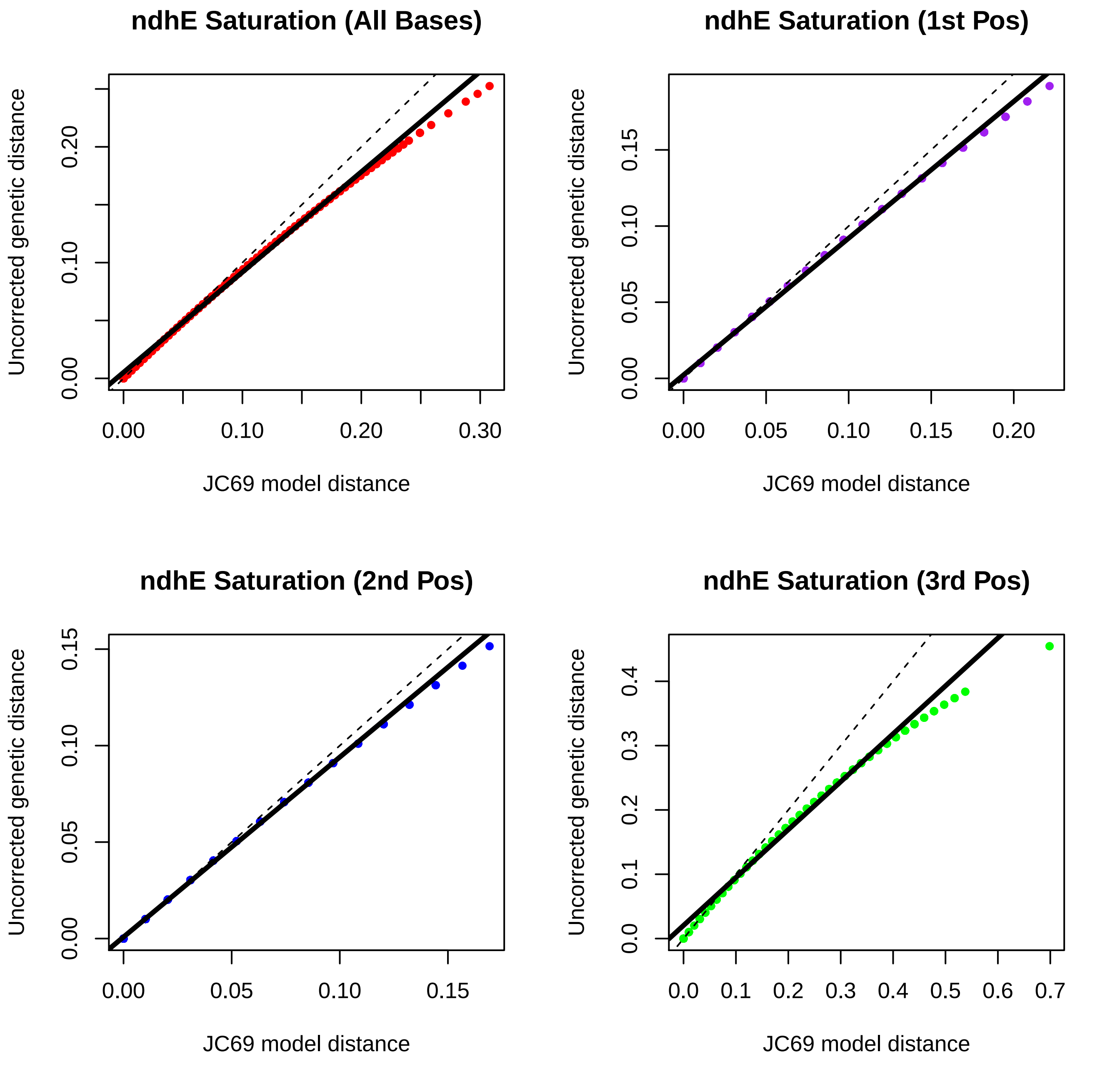
ndhE Saturation (All Bases)
Uncorrected genetic distance
JC69 model distance
0.00
0.10
0.20
0.30
ndhE Saturation (1st Pos)
Uncorrected genetic distance
JC69 model distance
0.00
0.05
0.10
0.15
0.20
ndhE Saturation (2nd Pos)
Uncorrected genetic distance
JC69 model distance
0.00
0.05
0.10
0.15
ndhE Saturation (3rd Pos)
Uncorrected genetic distance
JC69 model distance
0.0
0.1
0.2
0.3
0.4
0.5
0.6
0.7

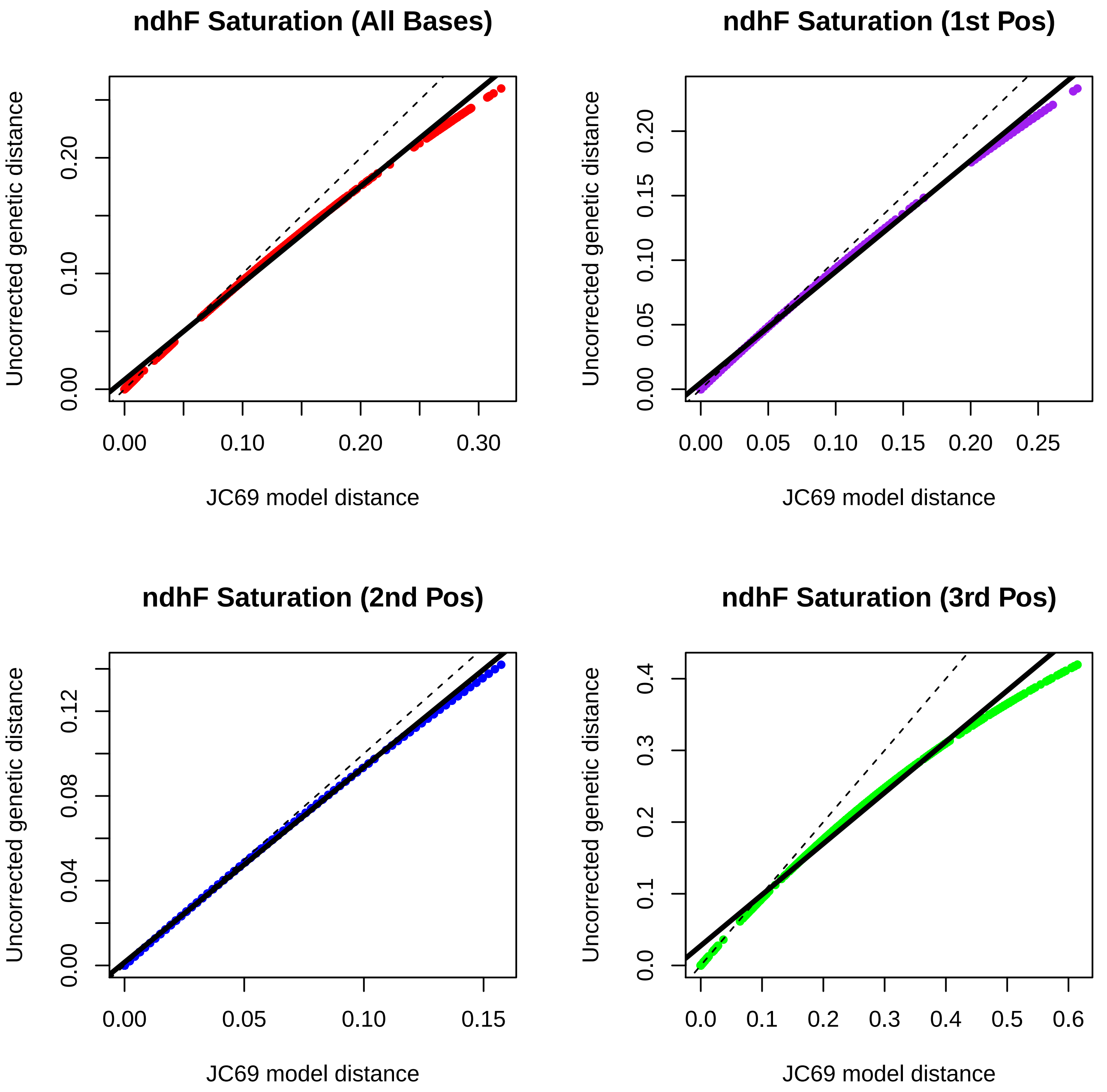
ndhF Saturation (All Bases)
Uncorrected genetic distance
0.00
0.10
0.20
0.00
0.10
0.20
0.30
JC69 model distance
ndhF Saturation (1st Pos)
Uncorrected genetic distance
0.00
0.05
0.10
0.15
0.20
0.00
0.05
0.10
0.15
0.20
0.25
JC69 model distance
ndhF Saturation (2nd Pos)
Uncorrected genetic distance
0.00
0.04
0.08
0.12
0.00
0.05
0.10
0.15
JC69 model distance
ndhF Saturation (3rd Pos)
Uncorrected genetic distance
0.0
0.1
0.2
0.3
0.4
0.0
0.1
0.2
0.3
0.4
0.5
0.6
JC69 model distance

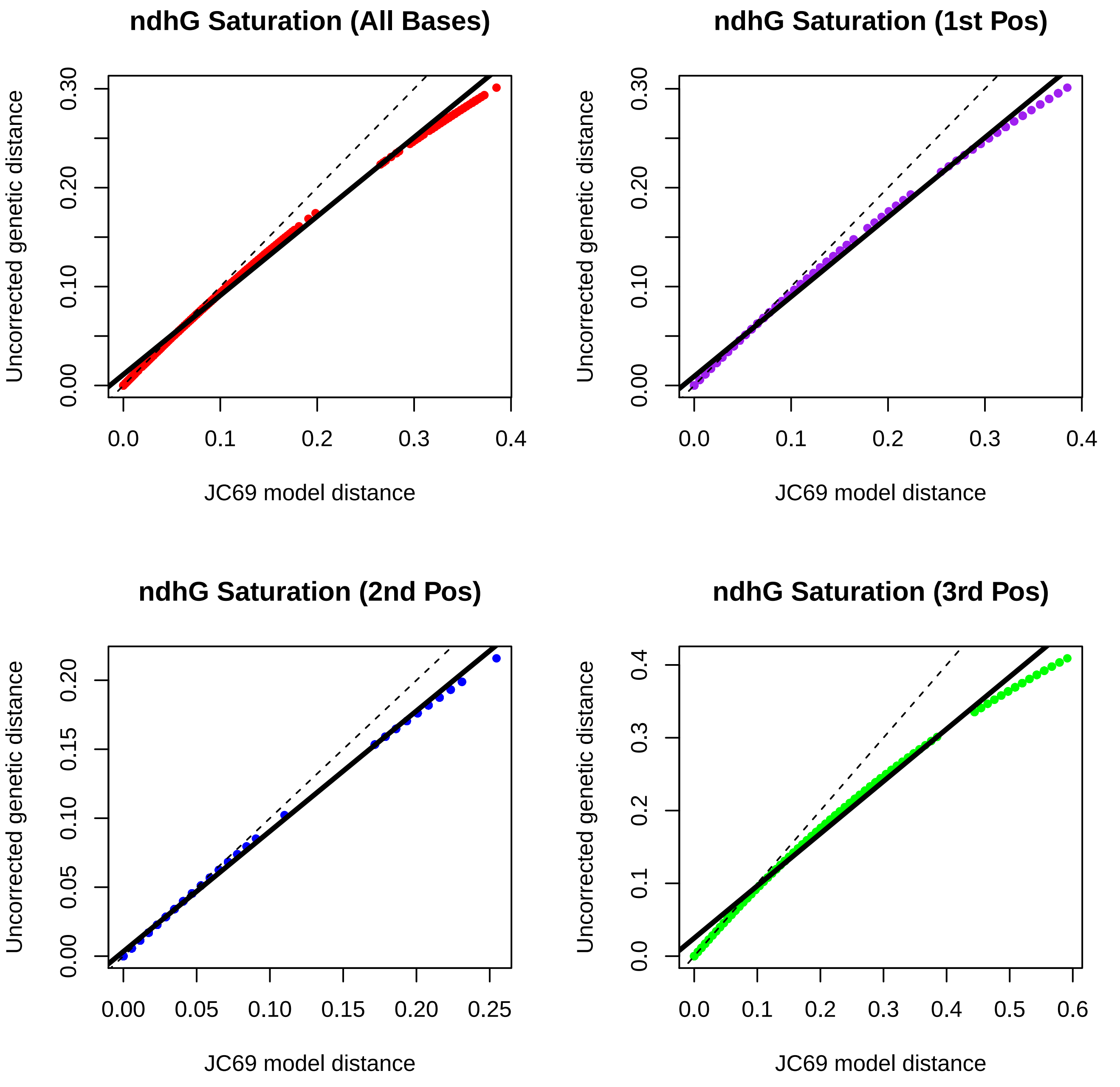
ndhG Saturation (All Bases)
ndhG Saturation (1st Pos)
ndhG Saturation (2nd Pos)
ndhG Saturation (3rd Pos)
Uncorrected genetic distance
JC69 model distance
0.00
0.10
0.20
0.30
0.0
0.1
0.2
0.3
0.4
0.05
0.15
0.25
0.5
0.6

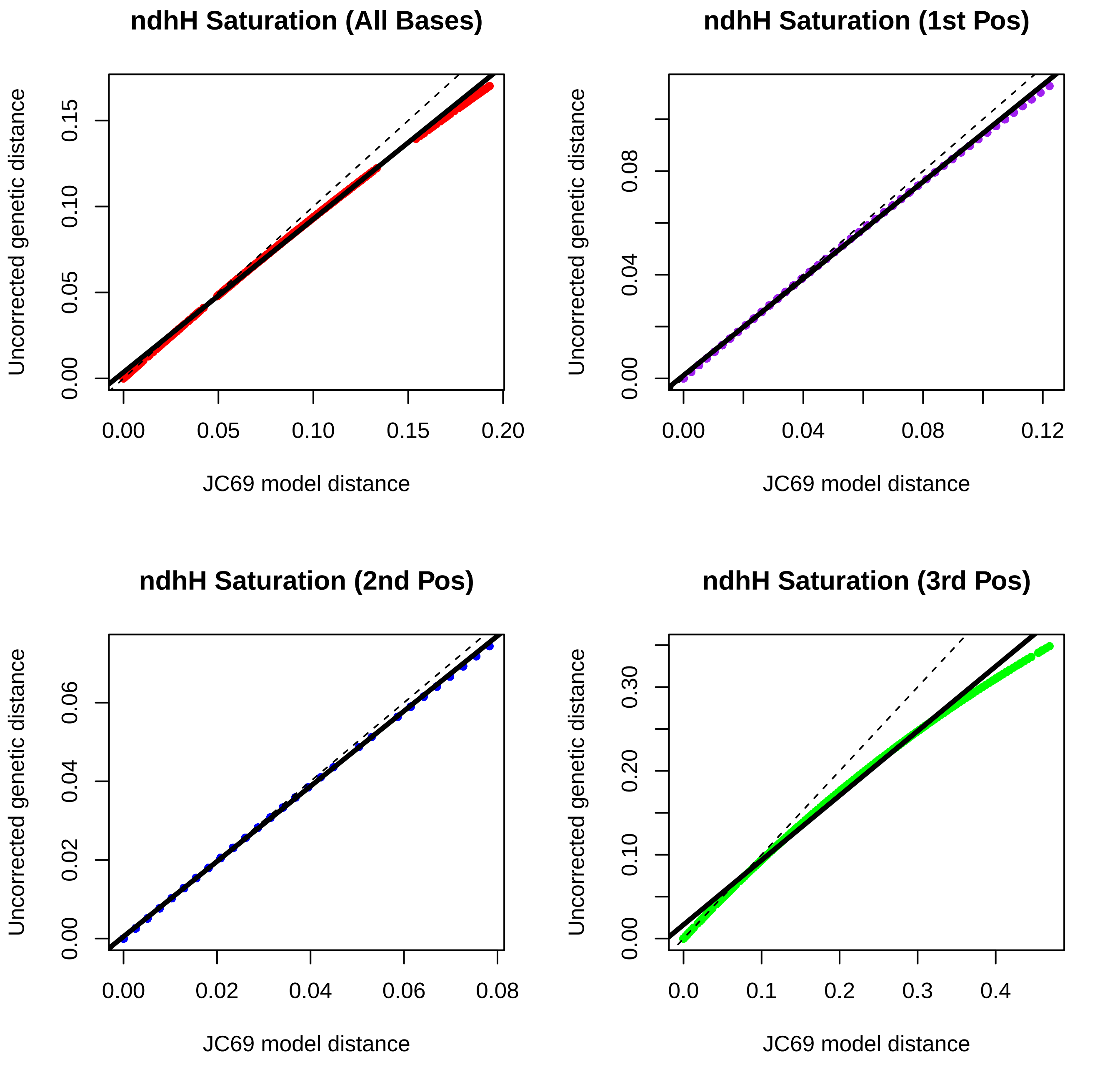
ndhH Saturation (All Bases)
ndhH Saturation (1st Pos)
ndhH Saturation (2nd Pos)
ndhH Saturation (3rd Pos)
Uncorrected genetic distance
JC69 model distance
0.00
0.05
0.10
0.15
0.20
0.04
0.08
0.12
0.02
0.06
0.0
0.1
0.2
0.3
0.4
0.30

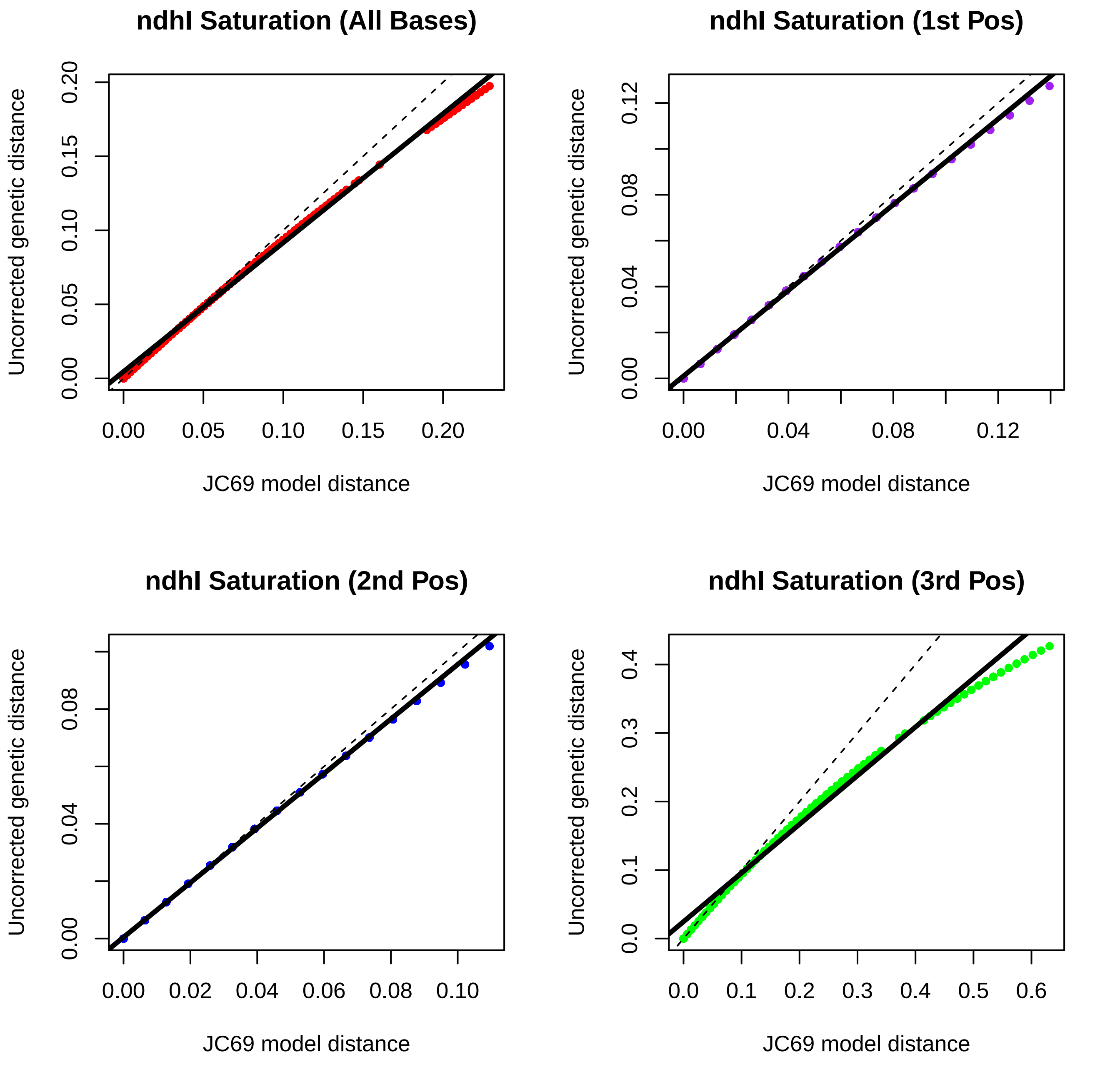
ndhI Saturation (All Bases)
ndhI Saturation (1st Pos)
ndhI Saturation (2nd Pos)
ndhI Saturation (3rd Pos)
Uncorrected genetic distance
JC69 model distance
0.00
0.05
0.10
0.15
0.20
0.04
0.08
0.12
0.02
0.06
0.0
0.1
0.2
0.3
0.4
0.5
0.6

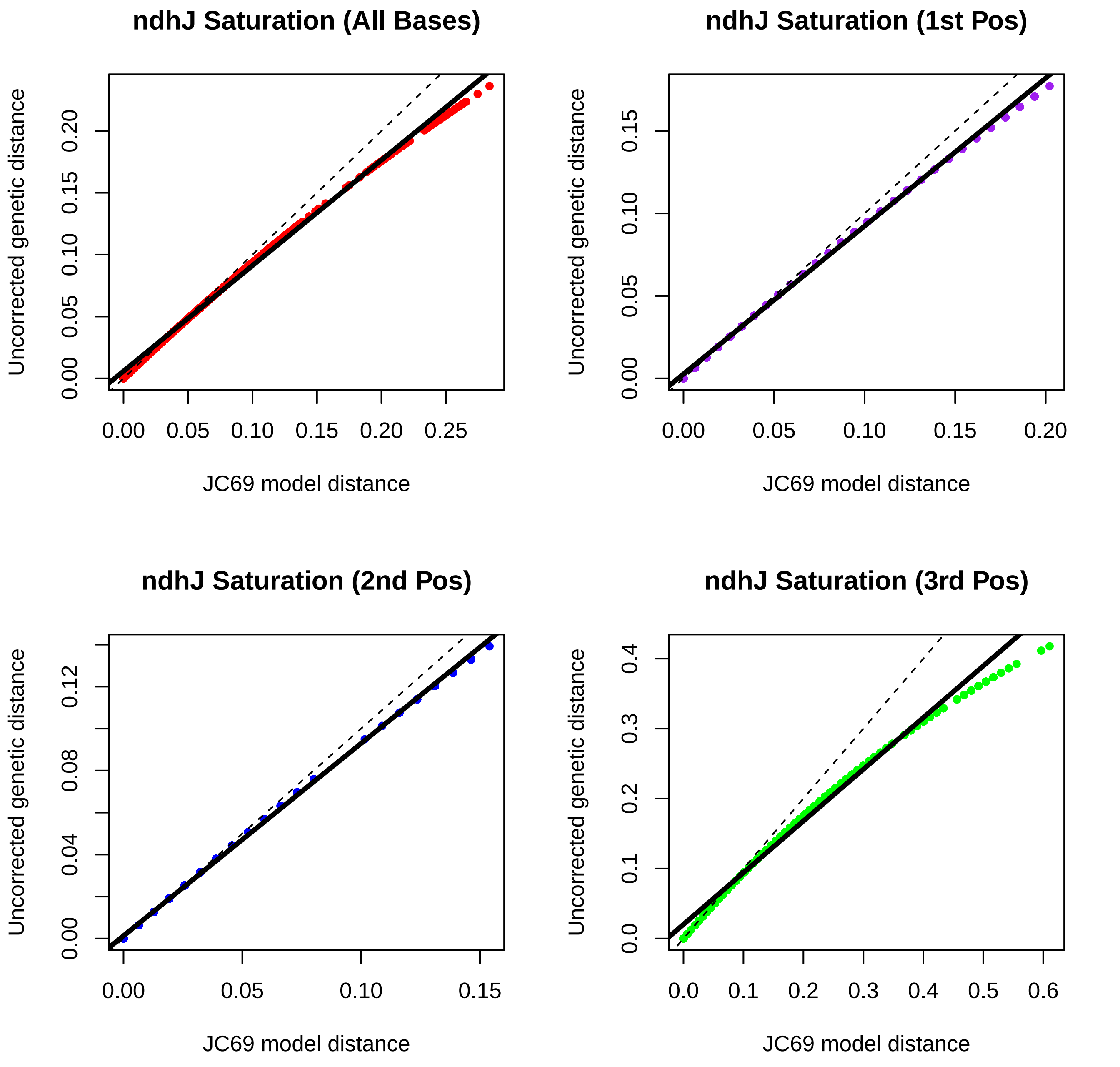

ndhJ Saturation (All Bases)
Uncorrected genetic distance
JC69 model distance
0.00
0.05
0.10
0.15
0.20
0.25
ndhJ Saturation (1st Pos)
Uncorrected genetic distance
JC69 model distance
0.00
0.05
0.10
0.15
0.20
ndhJ Saturation (2nd Pos)
Uncorrected genetic distance
JC69 model distance
0.00
0.04
0.08
0.12
0.05
0.10
0.15
ndhJ Saturation (3rd Pos)
Uncorrected genetic distance
JC69 model distance
0.0
0.1
0.2
0.3
0.4
0.5
0.6

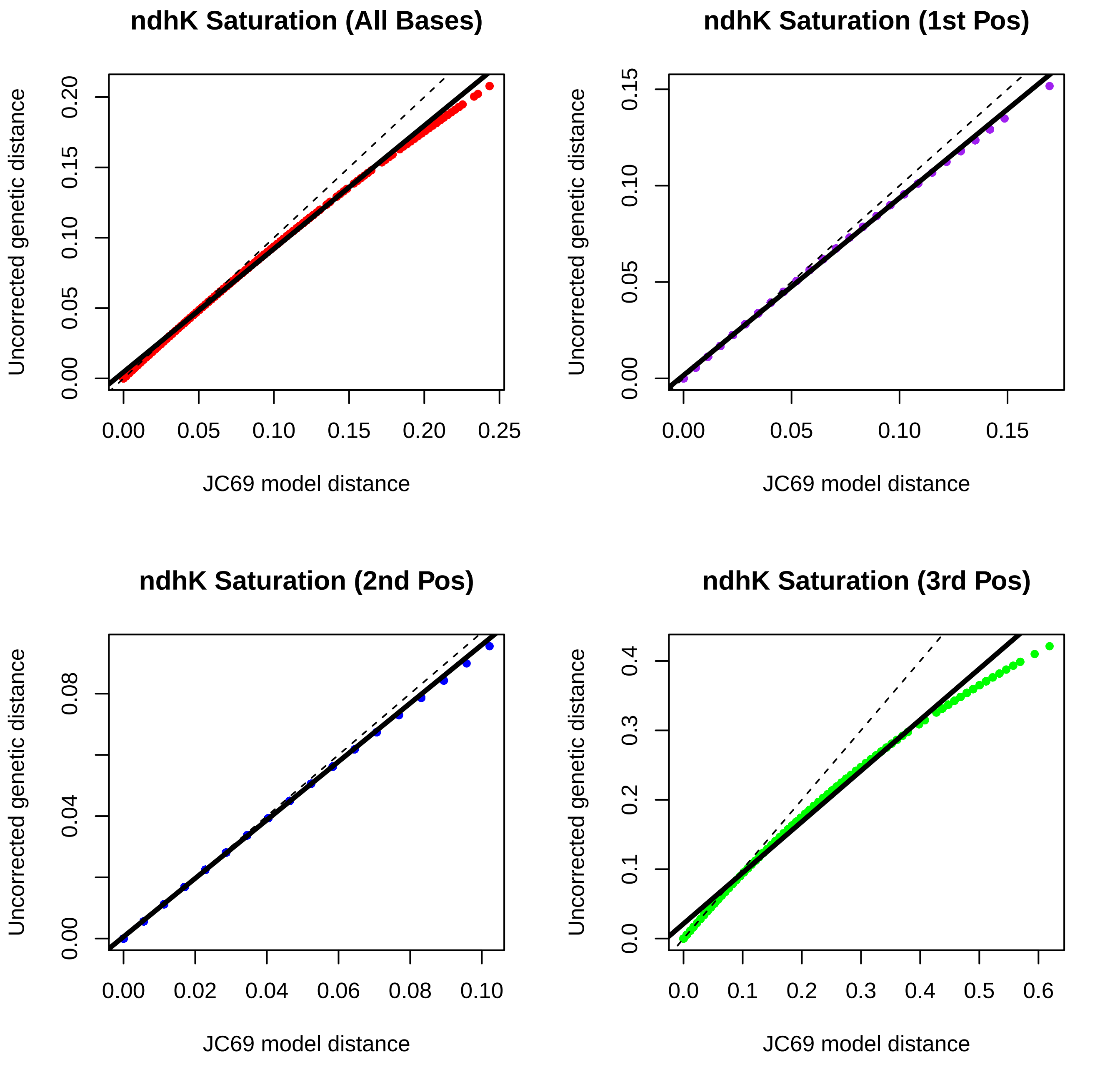
ndhK Saturation (All Bases)
ndhK Saturation (1st Pos)
ndhK Saturation (2nd Pos)
ndhK Saturation (3rd Pos)
Uncorrected genetic distance
JC69 model distance
0.00
0.05
0.10
0.15
0.20
0.25
0.02
0.04
0.06
0.08
0.0
0.1
0.2
0.3
0.4
0.5
0.6

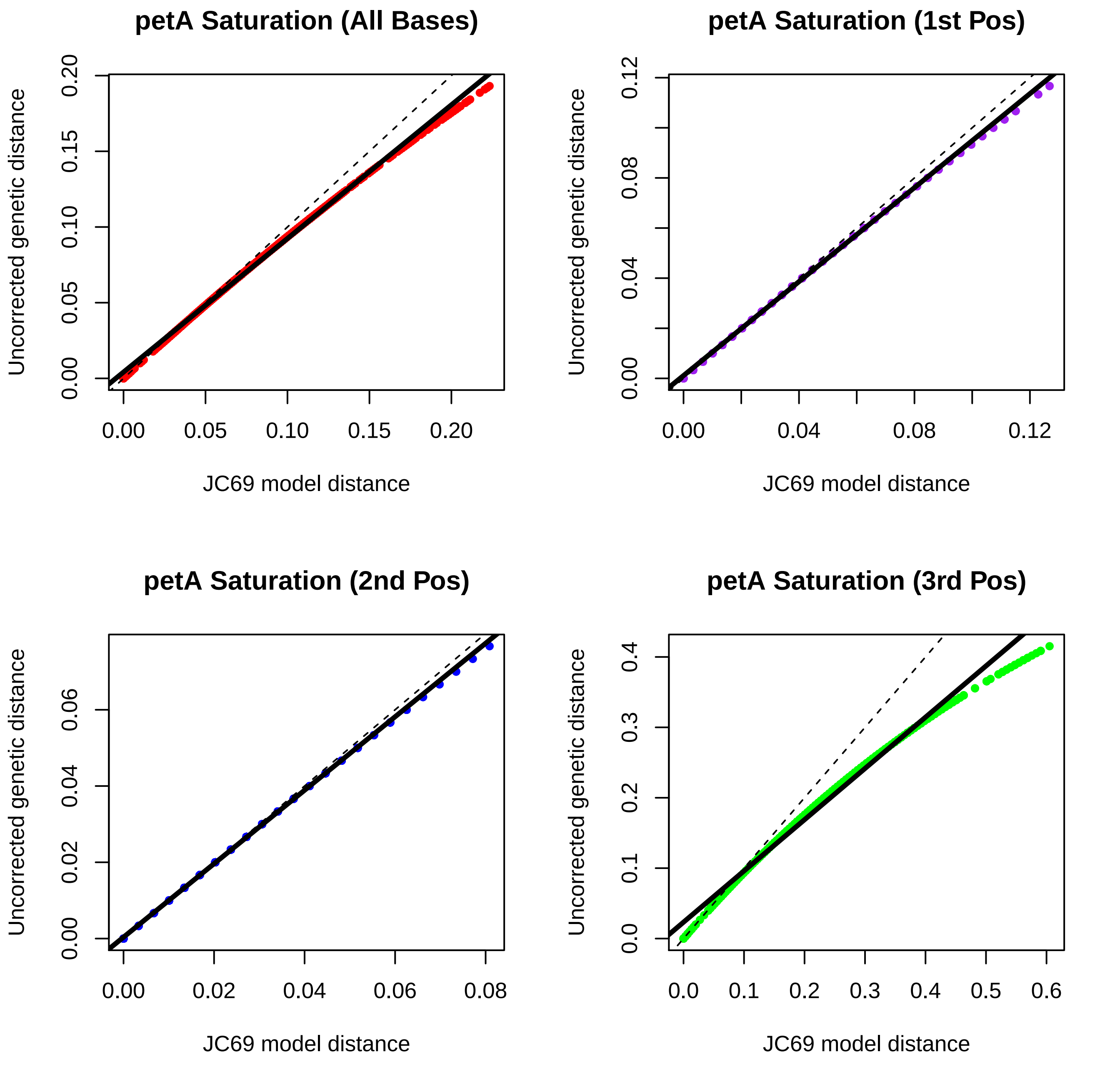
petA Saturation (All Bases)
Uncorrected genetic distance
JC69 model distance
0.00
0.05
0.10
0.15
0.20
petA Saturation (1st Pos)
0.00
0.04
0.08
0.12
petA Saturation (2nd Pos)
0.00
0.02
0.04
0.06
0.08
petA Saturation (3rd Pos)
0.0
0.1
0.2
0.3
0.4
0.5
0.6

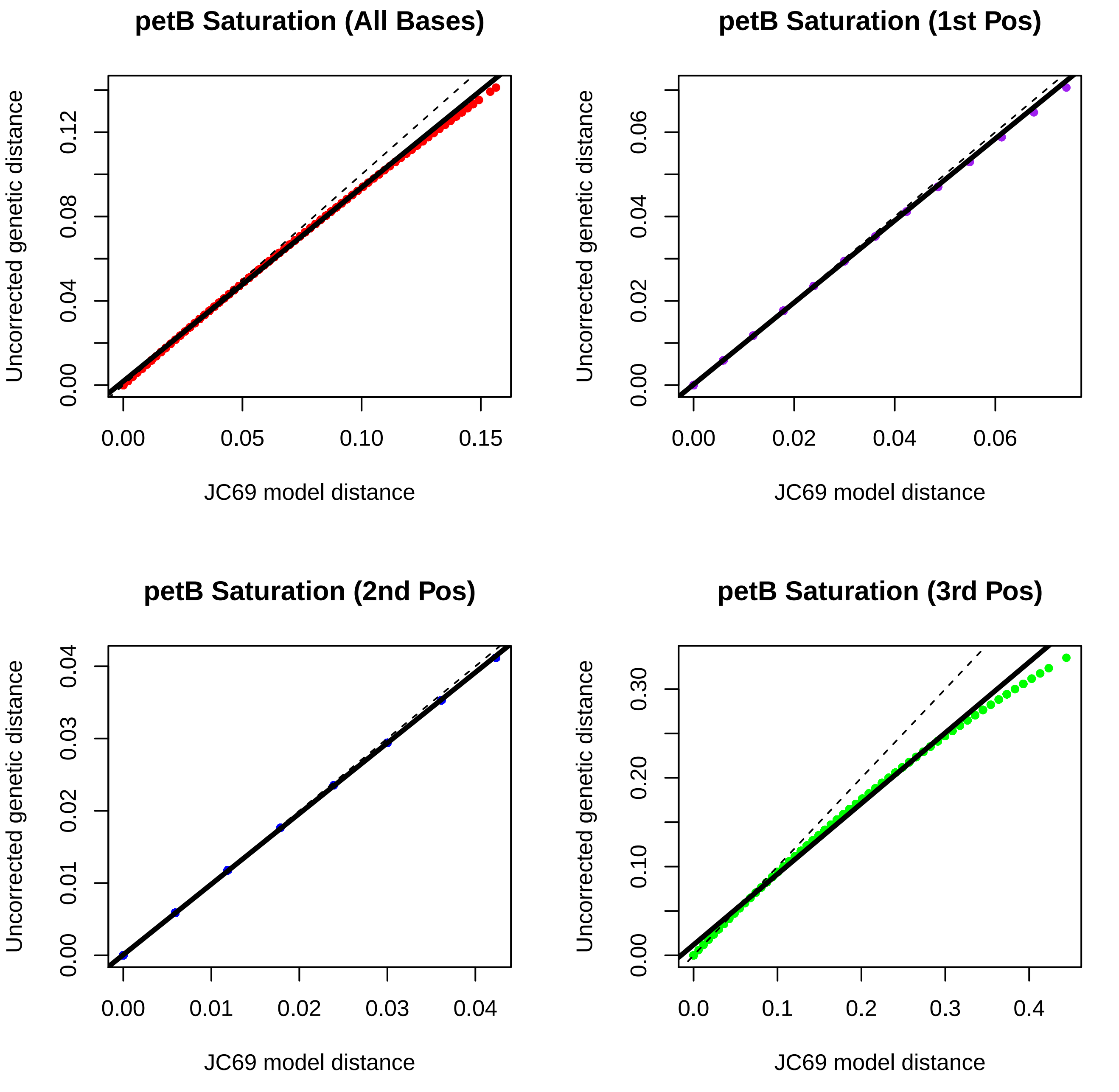
petB Saturation (All Bases)
Uncorrected genetic distance
JC69 model distance
0.00
0.04
0.08
0.12
0.00
0.05
0.10
0.15
petB Saturation (1st Pos)
Uncorrected genetic distance
JC69 model distance
0.00
0.02
0.04
0.06
0.00
0.02
0.04
0.06
petB Saturation (2nd Pos)
Uncorrected genetic distance
JC69 model distance
0.00
0.01
0.02
0.03
0.04
0.00
0.01
0.02
0.03
0.04
petB Saturation (3rd Pos)
Uncorrected genetic distance
JC69 model distance
0.00
0.10
0.20
0.30
0.0
0.1
0.2
0.3
0.4

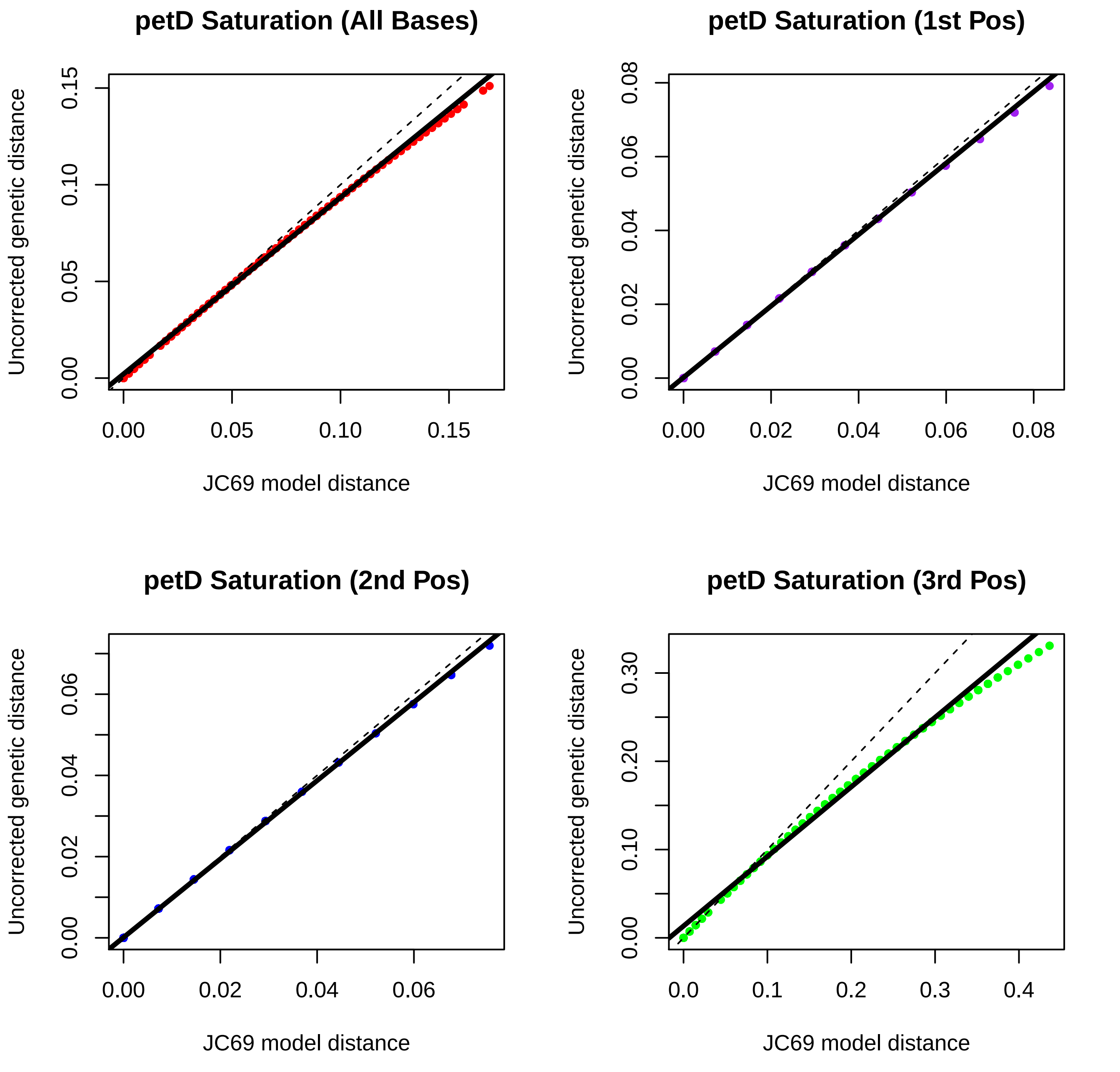

petD Saturation (All Bases)
petD Saturation (1st Pos)
petD Saturation (2nd Pos)
petD Saturation (3rd Pos)
Uncorrected genetic distance
JC69 model distance
0.00
0.05
0.10
0.15
0.02
0.04
0.06
0.08
0.0
0.1
0.2
0.3
0.4
0.20
0.30

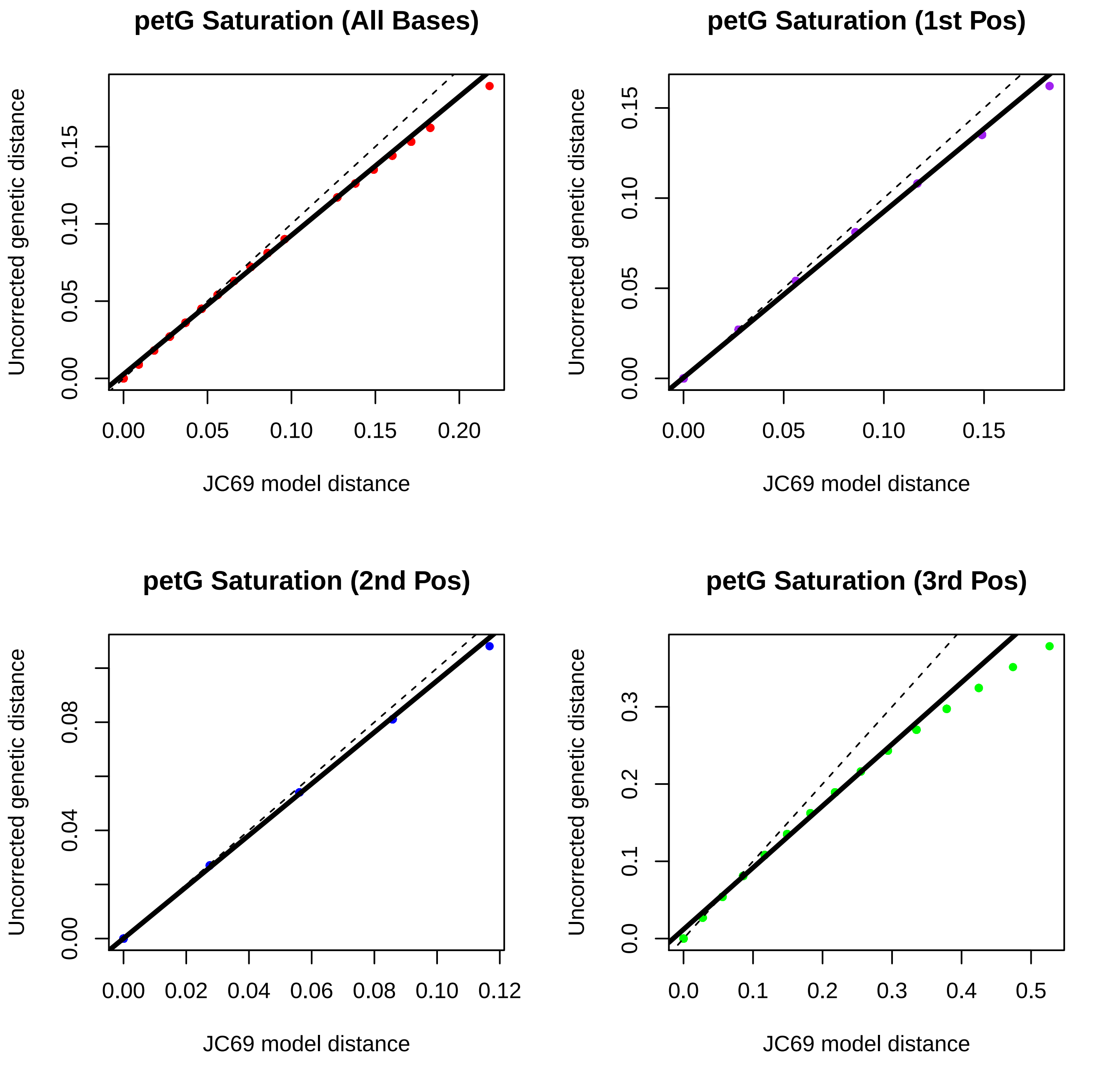
petG Saturation (All Bases)
Uncorrected genetic distance
JC69 model distance
0.00
0.05
0.10
0.15
0.20
petG Saturation (1st Pos)
Uncorrected genetic distance
JC69 model distance
petG Saturation (2nd Pos)
Uncorrected genetic distance
JC69 model distance
0.00
0.02
0.04
0.06
0.08
0.10
0.12
petG Saturation (3rd Pos)
Uncorrected genetic distance
JC69 model distance
0.0
0.1
0.2
0.3
0.4
0.5

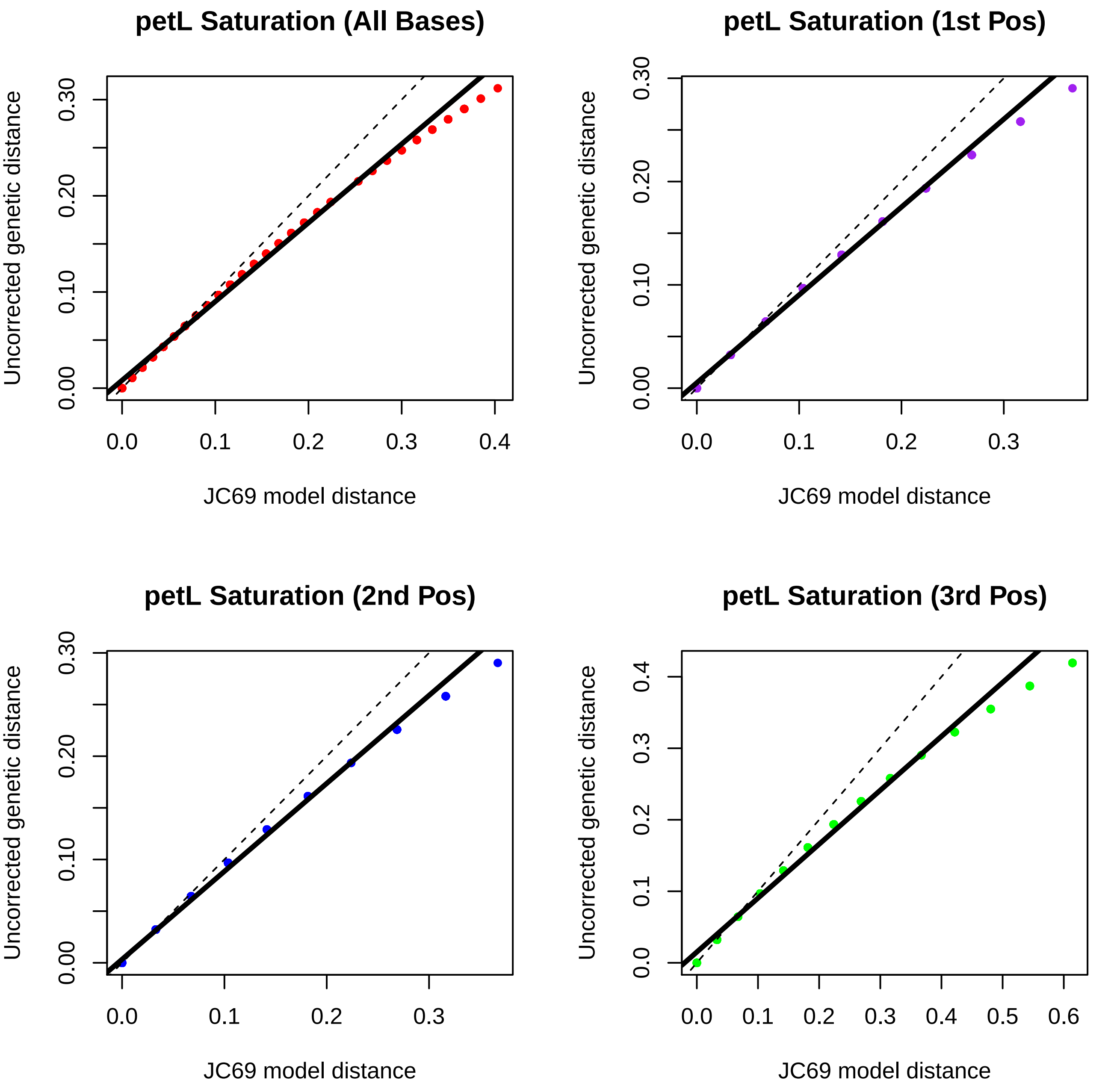

petL Saturation (All Bases)
petL Saturation (1st Pos)
petL Saturation (2nd Pos)
petL Saturation (3rd Pos)
Uncorrected genetic distance
JC69 model distance
0.00
0.10
0.20
0.30
0.0
0.1
0.2
0.3
0.4
0.5
0.6

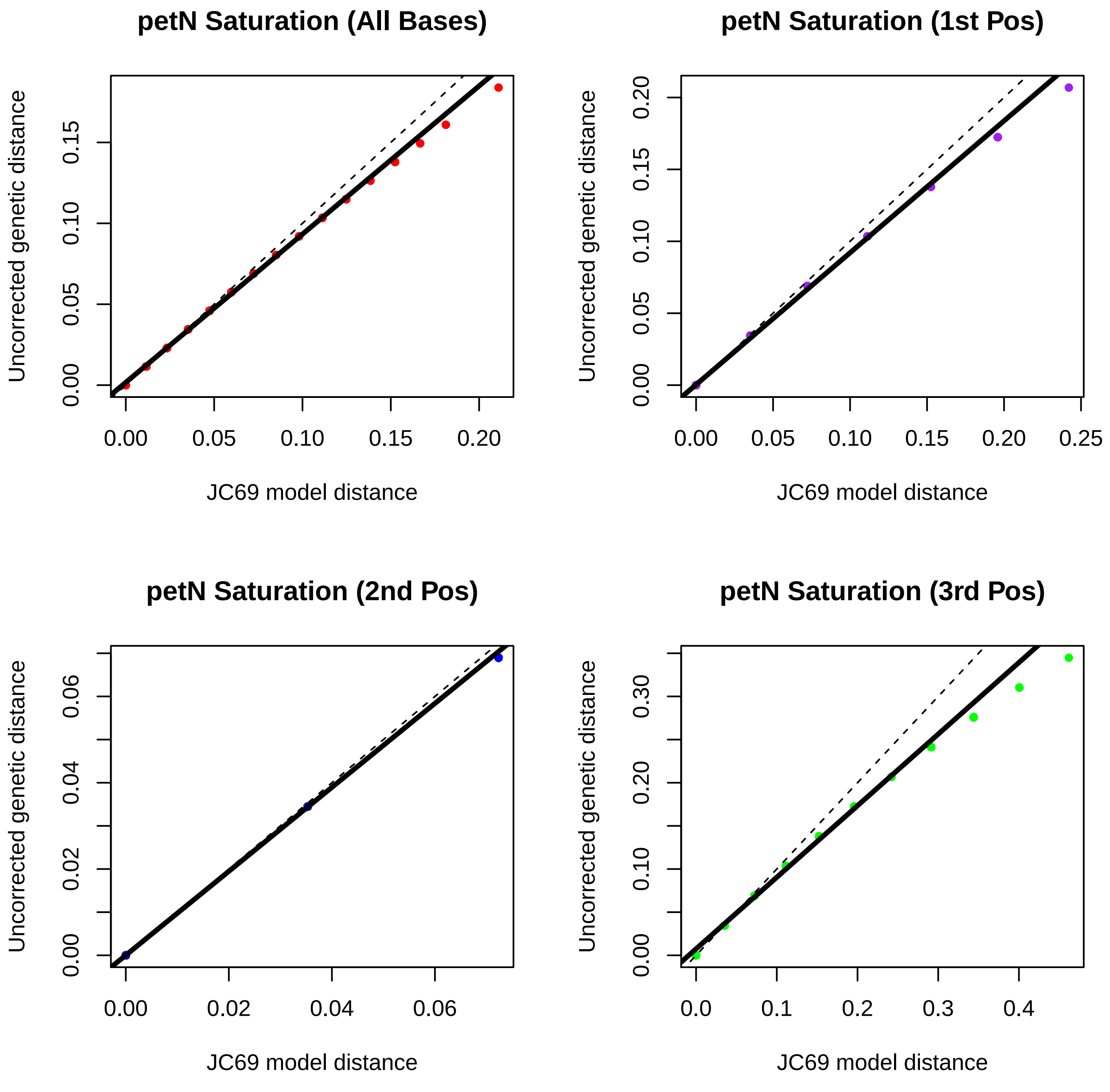
petN Saturation (All Bases)
petN Saturation (1st Pos)
petN Saturation (2nd Pos)
petN Saturation (3rd Pos)
Uncorrected genetic distance
JC69 model distance
0.00
0.05
0.10
0.15
0.20
0.25
0.02
0.04
0.06
0.0
0.1
0.2
0.3
0.4
0.30

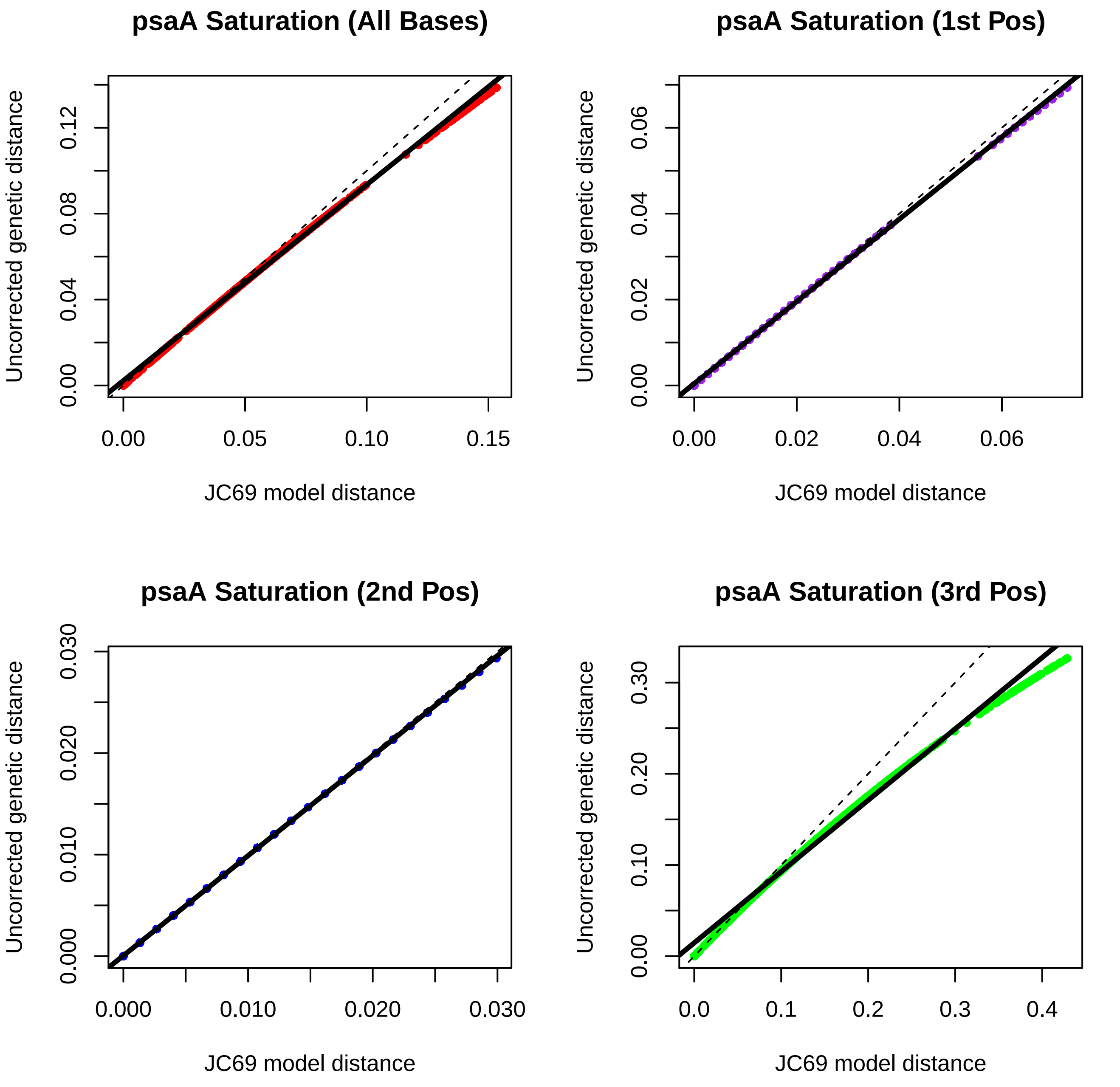
psaA Saturation (All Bases)
Uncorrected genetic distance
0.00
0.04
0.08
0.12
0.00
0.05
0.10
0.15
JC69 model distance
psaA Saturation (1st Pos)
Uncorrected genetic distance
0.00
0.02
0.04
0.06
0.00
0.02
0.04
0.06
JC69 model distance
psaA Saturation (2nd Pos)
Uncorrected genetic distance
0.000
0.010
0.020
0.030
0.000
0.010
0.020
0.030
JC69 model distance
psaA Saturation (3rd Pos)
Uncorrected genetic distance
0.00
0.10
0.20
0.30
0.0
0.1
0.2
0.3
0.4
JC69 model distance

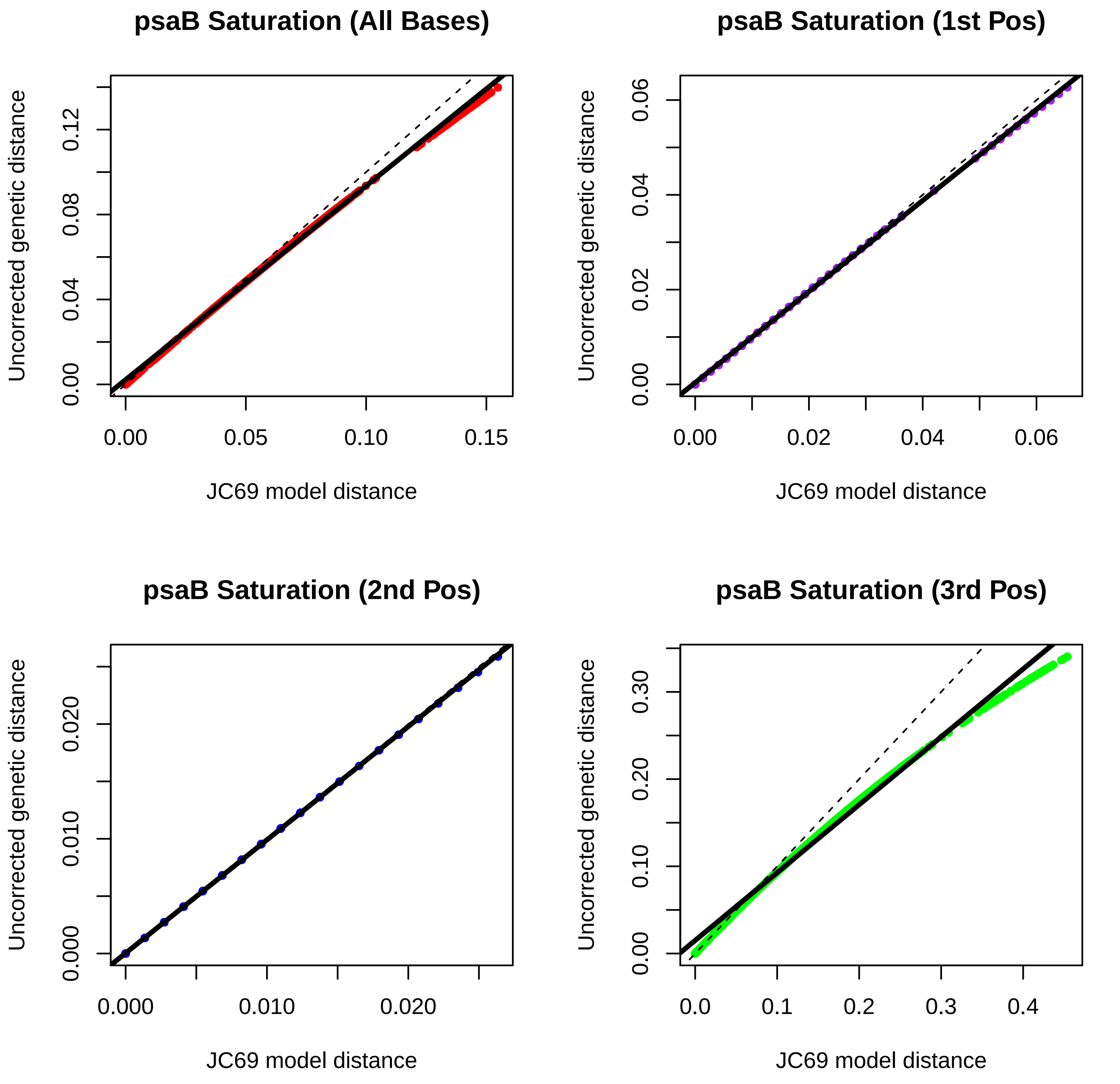
psaB Saturation (All Bases)
Uncorrected genetic distance
JC69 model distance
0.00
0.04
0.08
0.12
0.00
0.05
0.10
0.15
psaB Saturation (1st Pos)
Uncorrected genetic distance
JC69 model distance
0.00
0.02
0.04
0.06
0.00
0.02
0.04
0.06
psaB Saturation (2nd Pos)
Uncorrected genetic distance
JC69 model distance
0.000
0.010
0.020
0.000
0.010
0.020
psaB Saturation (3rd Pos)
Uncorrected genetic distance
JC69 model distance
0.00
0.10
0.20
0.30
0.0
0.1
0.2
0.3
0.4

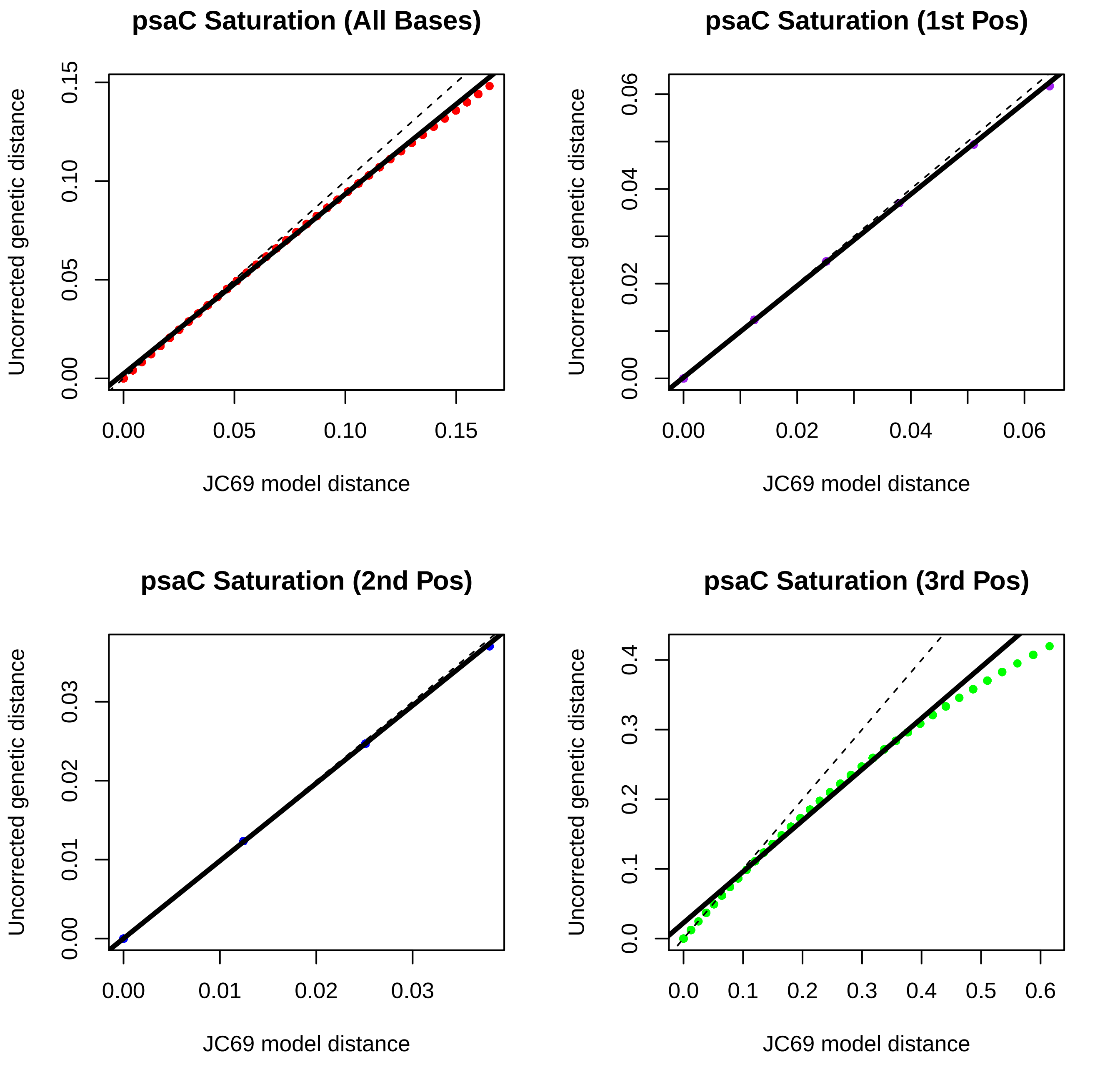

psaC Saturation (All Bases)
Uncorrected genetic distance
0.00
0.05
0.10
0.15
JC69 model distance
0.00
0.05
0.10
0.15
psaC Saturation (1st Pos)
Uncorrected genetic distance
0.00
0.02
0.04
0.06
JC69 model distance
0.00
0.02
0.04
0.06
psaC Saturation (2nd Pos)
Uncorrected genetic distance
0.00
0.01
0.02
0.03
JC69 model distance
0.00
0.01
0.02
0.03
psaC Saturation (3rd Pos)
Uncorrected genetic distance
0.0
0.1
0.2
0.3
0.4
JC69 model distance
0.0
0.1
0.2
0.3
0.4
0.5
0.6

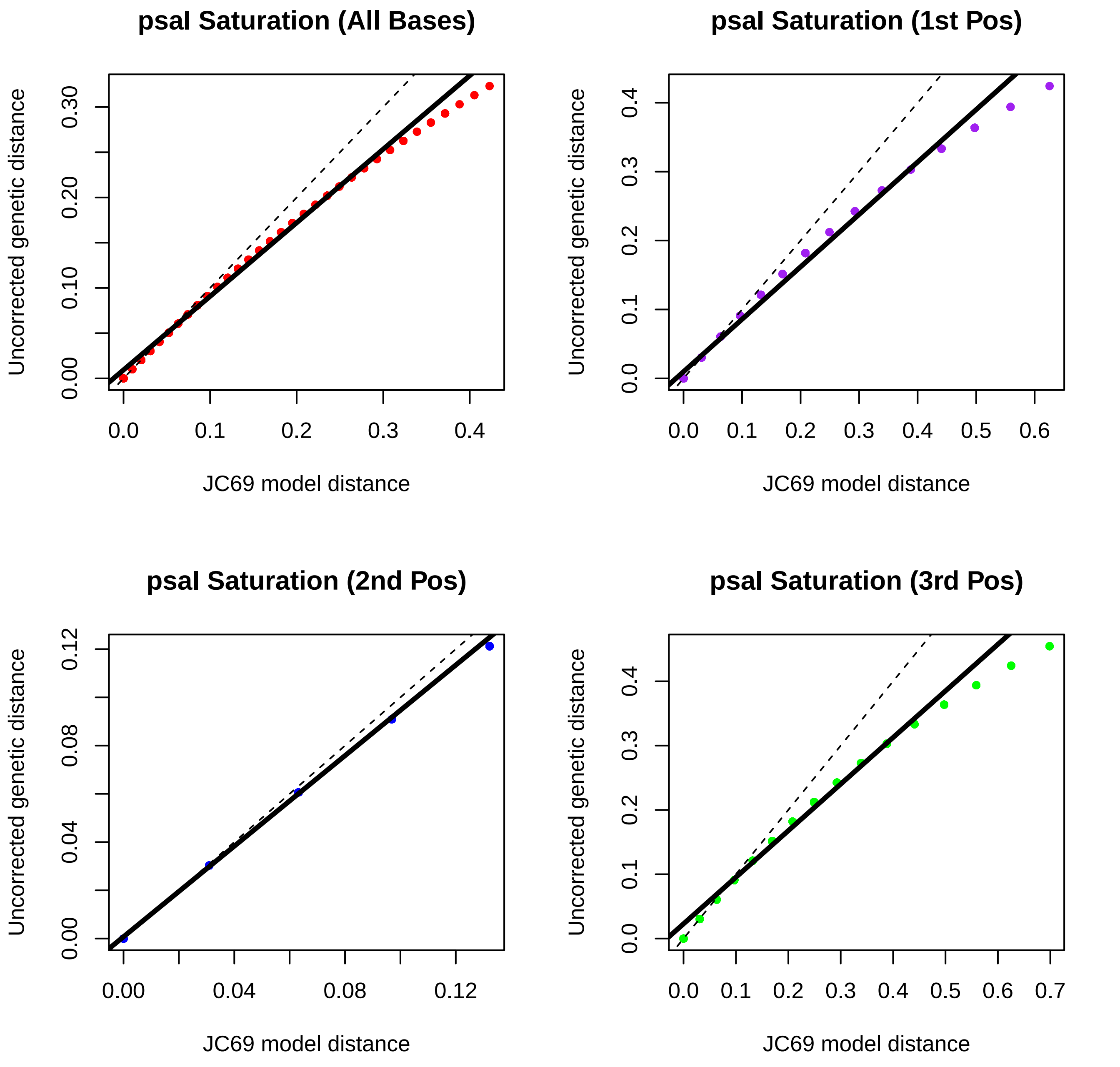
psal Saturation (All Bases)
psal Saturation (1st Pos)
psal Saturation (2nd Pos)
psal Saturation (3rd Pos)
Uncorrected genetic distance
JC69 model distance
0.00
0.10
0.20
0.30
0.0
0.1
0.2
0.3
0.4
0.5
0.6
0.7
0.04
0.08
0.12

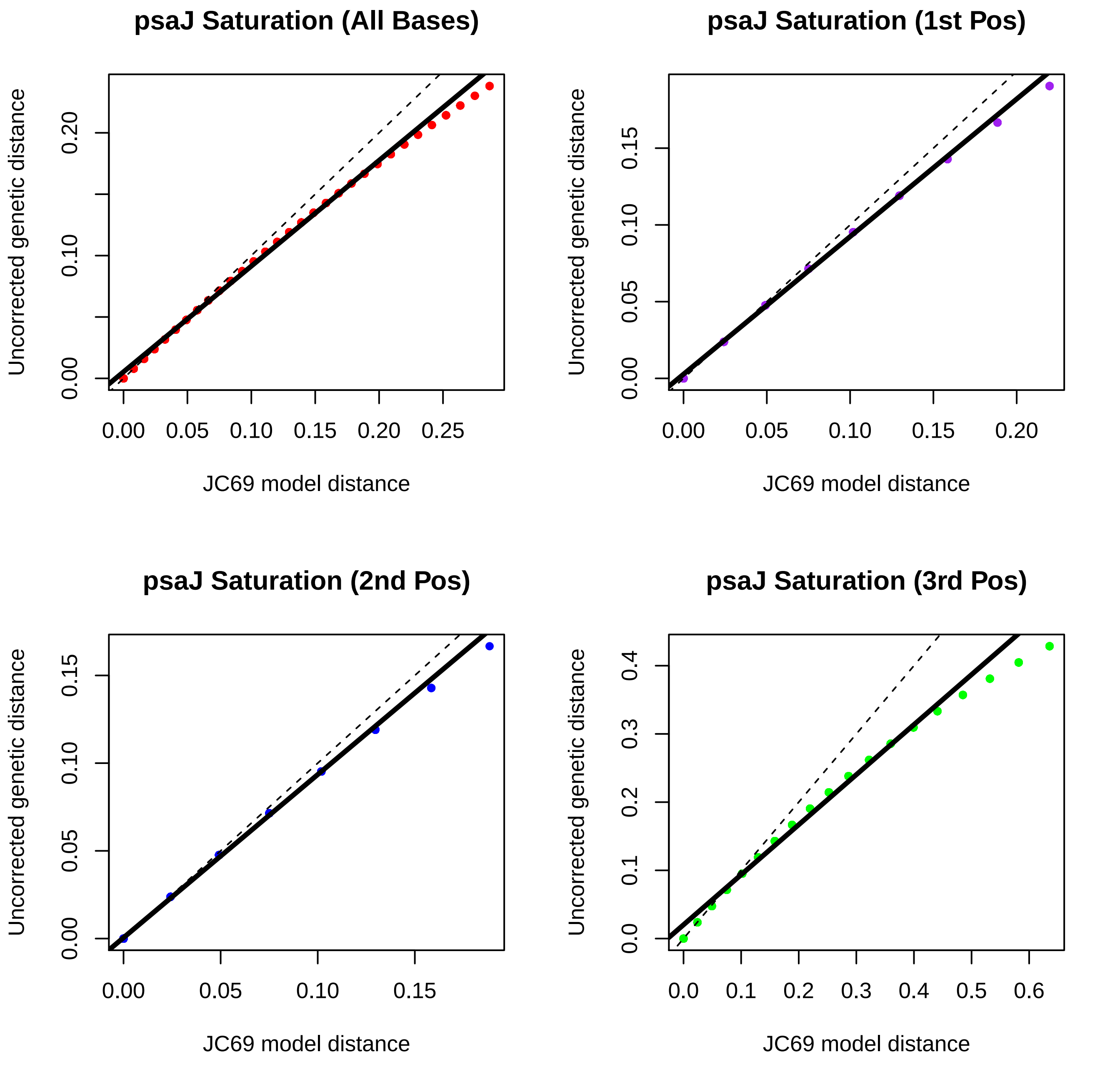

psaJ Saturation (All Bases)
psaJ Saturation (1st Pos)
psaJ Saturation (2nd Pos)
psaJ Saturation (3rd Pos)
Uncorrected genetic distance
JC69 model distance

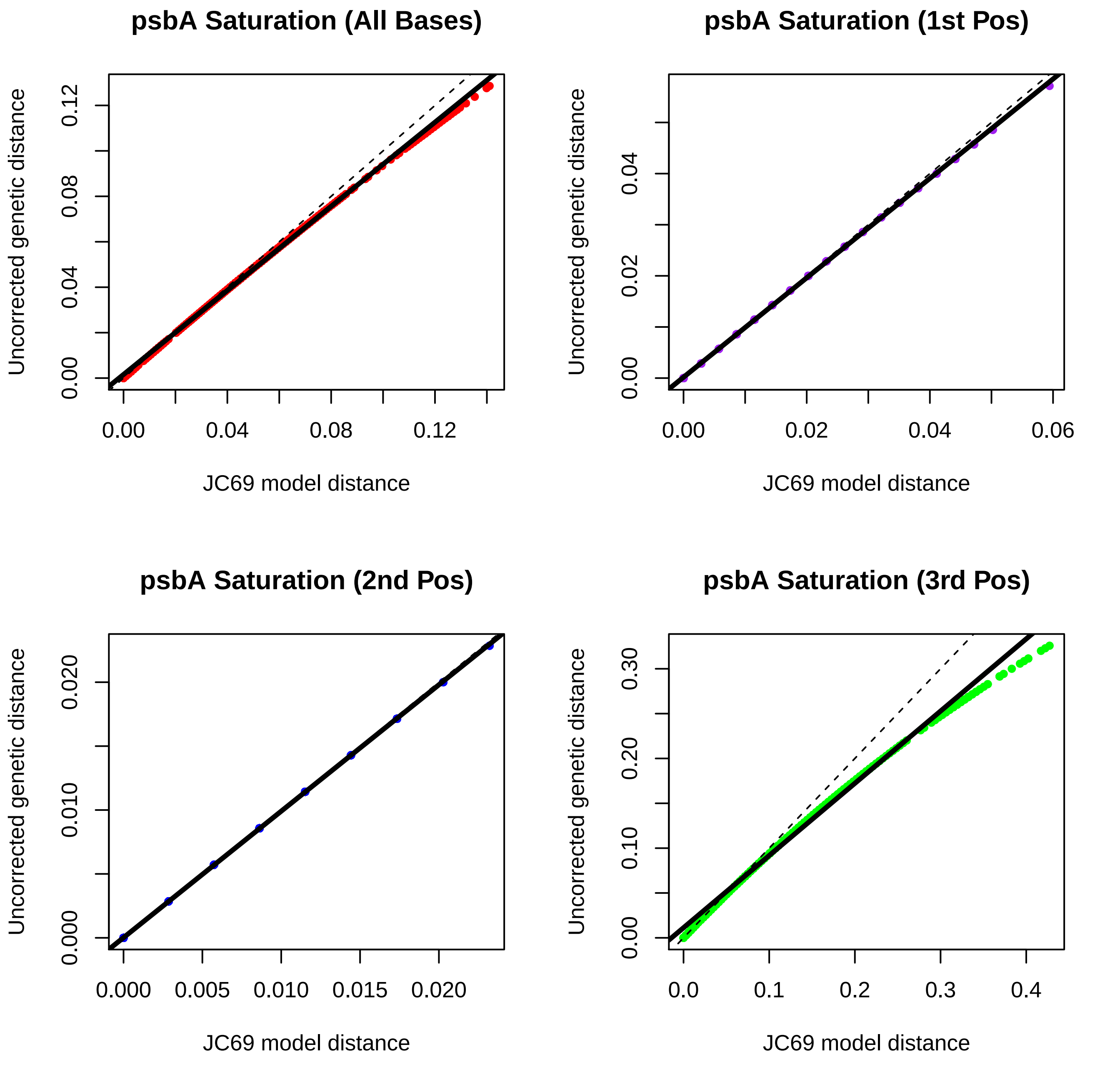
psbA Saturation (All Bases)
Uncorrected genetic distance
JC69 model distance
0.00
0.04
0.08
0.12
psbA Saturation (1st Pos)
0.00
0.02
0.04
0.06
psbA Saturation (2nd Pos)
0.000
0.005
0.010
0.015
0.020
psbA Saturation (3rd Pos)
0.0
0.1
0.2
0.3
0.4
0.00
0.10
0.20
0.30

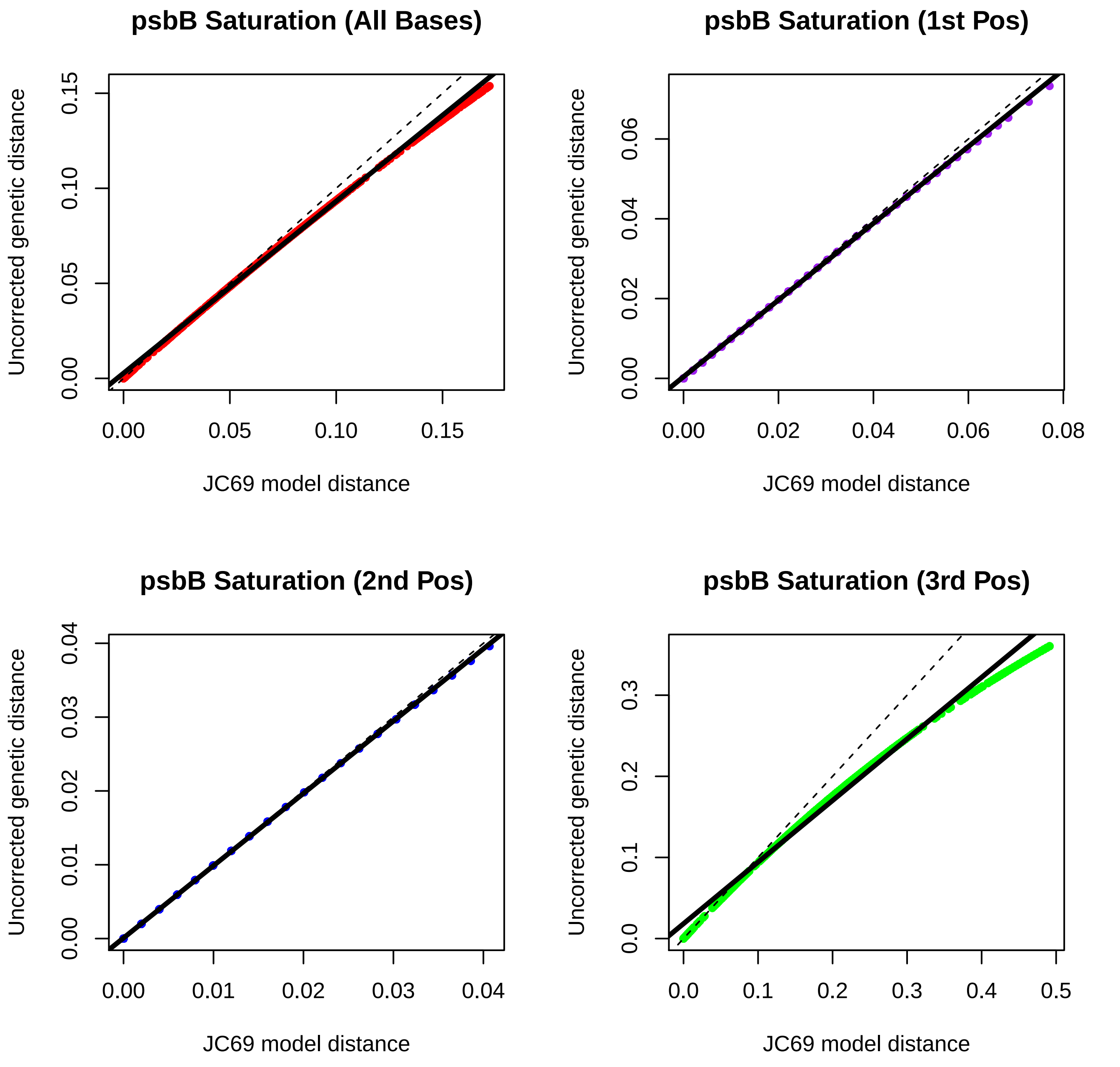
psbB Saturation (All Bases)
psbB Saturation (1st Pos)
psbB Saturation (2nd Pos)
psbB Saturation (3rd Pos)
Uncorrected genetic distance
JC69 model distance
0.00
0.05
0.10
0.15
0.02
0.04
0.06
0.08
0.01
0.03
0.0
0.1
0.2
0.3
0.4
0.5

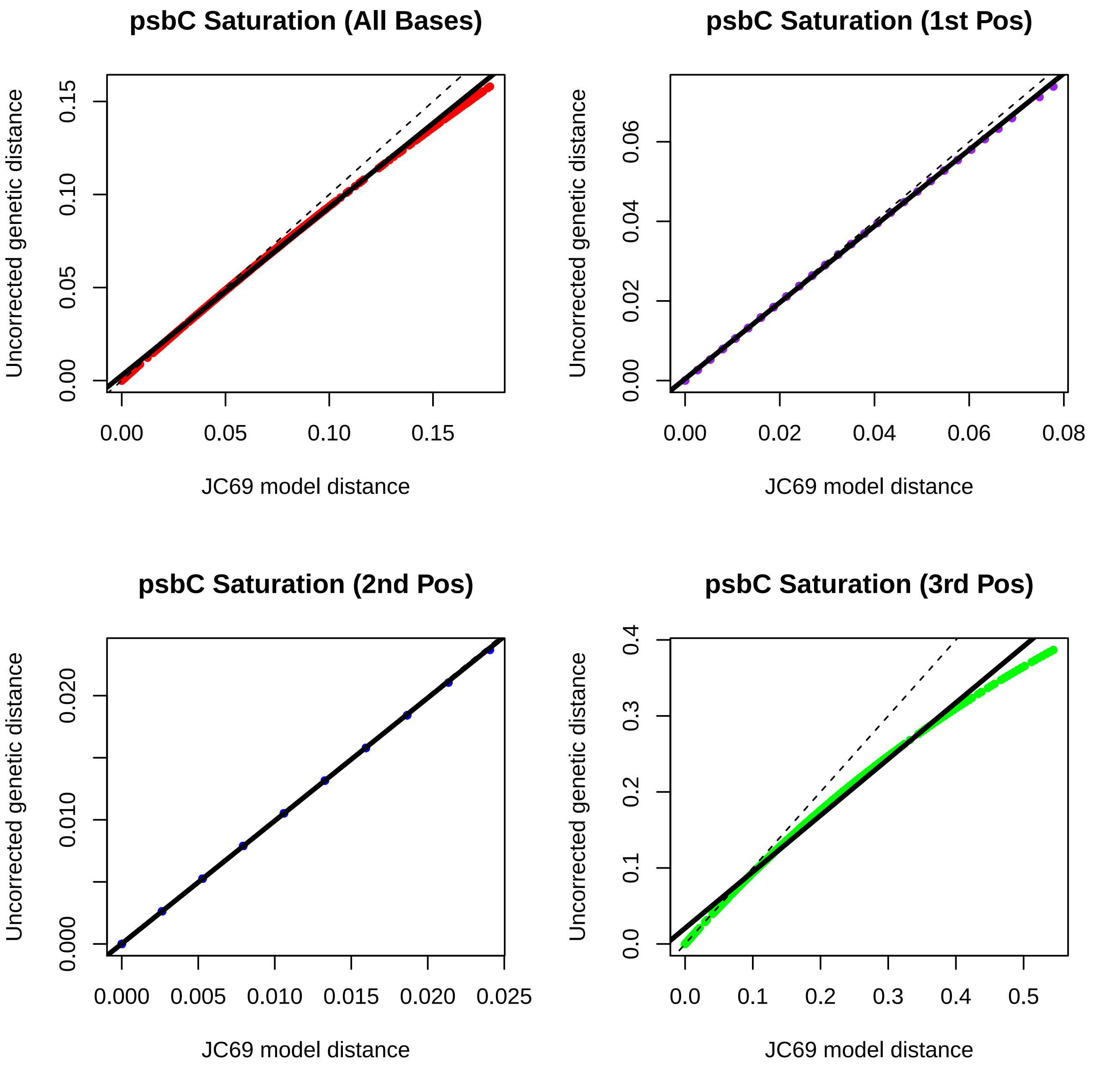
psbC Saturation (All Bases)
Uncorrected genetic distance
0.00
0.05
0.10
0.15
JC69 model distance
psbC Saturation (1st Pos)
Uncorrected genetic distance
0.00
0.02
0.04
0.06
0.08
JC69 model distance
psbC Saturation (2nd Pos)
Uncorrected genetic distance
0.000
0.005
0.010
0.015
0.020
0.025
JC69 model distance
psbC Saturation (3rd Pos)
Uncorrected genetic distance
0.0
0.1
0.2
0.3
0.4
0.5
JC69 model distance

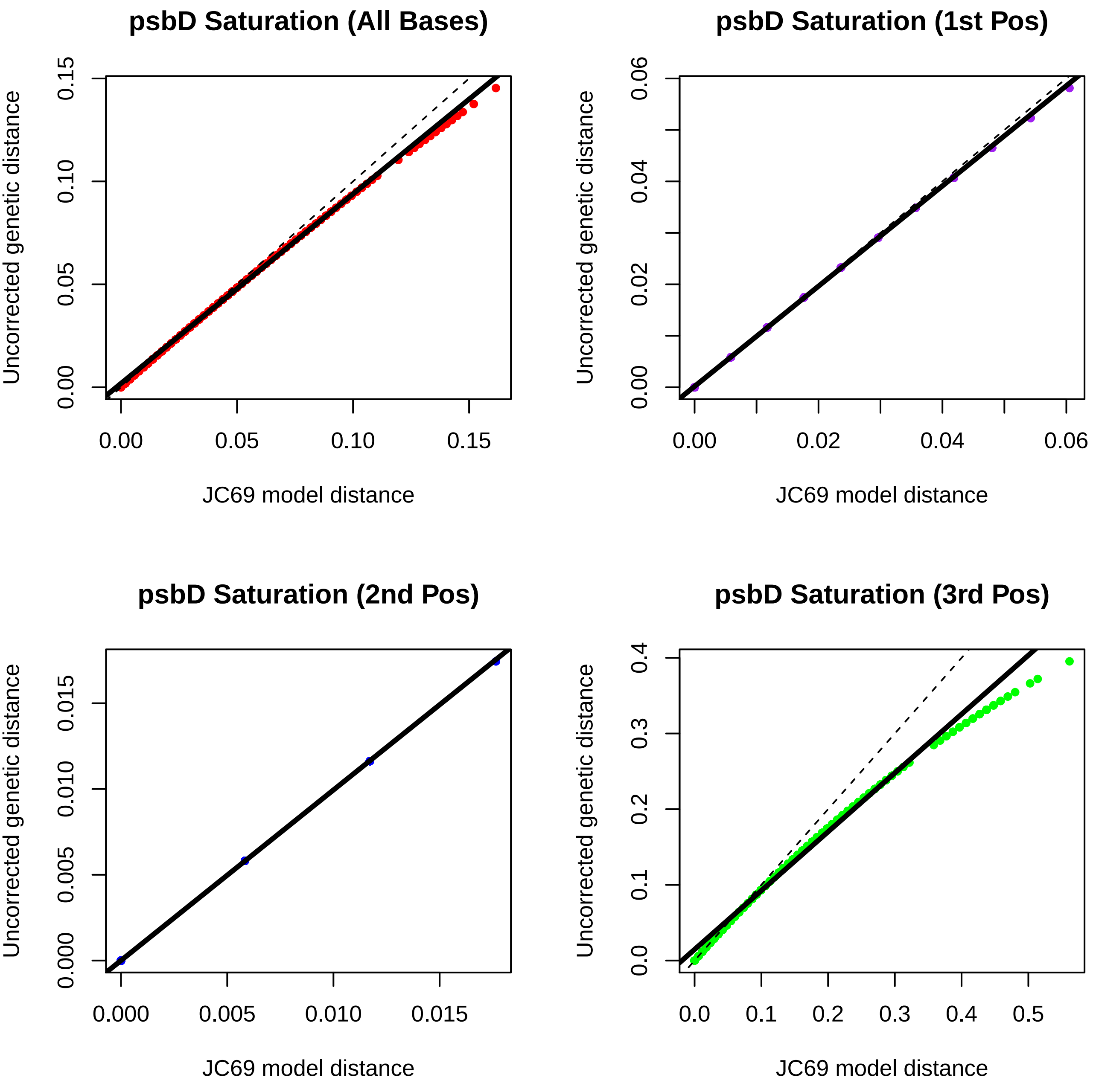
psbD Saturation (All Bases)
psbD Saturation (1st Pos)
psbD Saturation (2nd Pos)
psbD Saturation (3rd Pos)
Uncorrected genetic distance
JC69 model distance
0.00
0.05
0.10
0.15
0.02
0.04
0.06
0.000
0.005
0.010
0.015
0.0
0.1
0.2
0.3
0.4
0.5

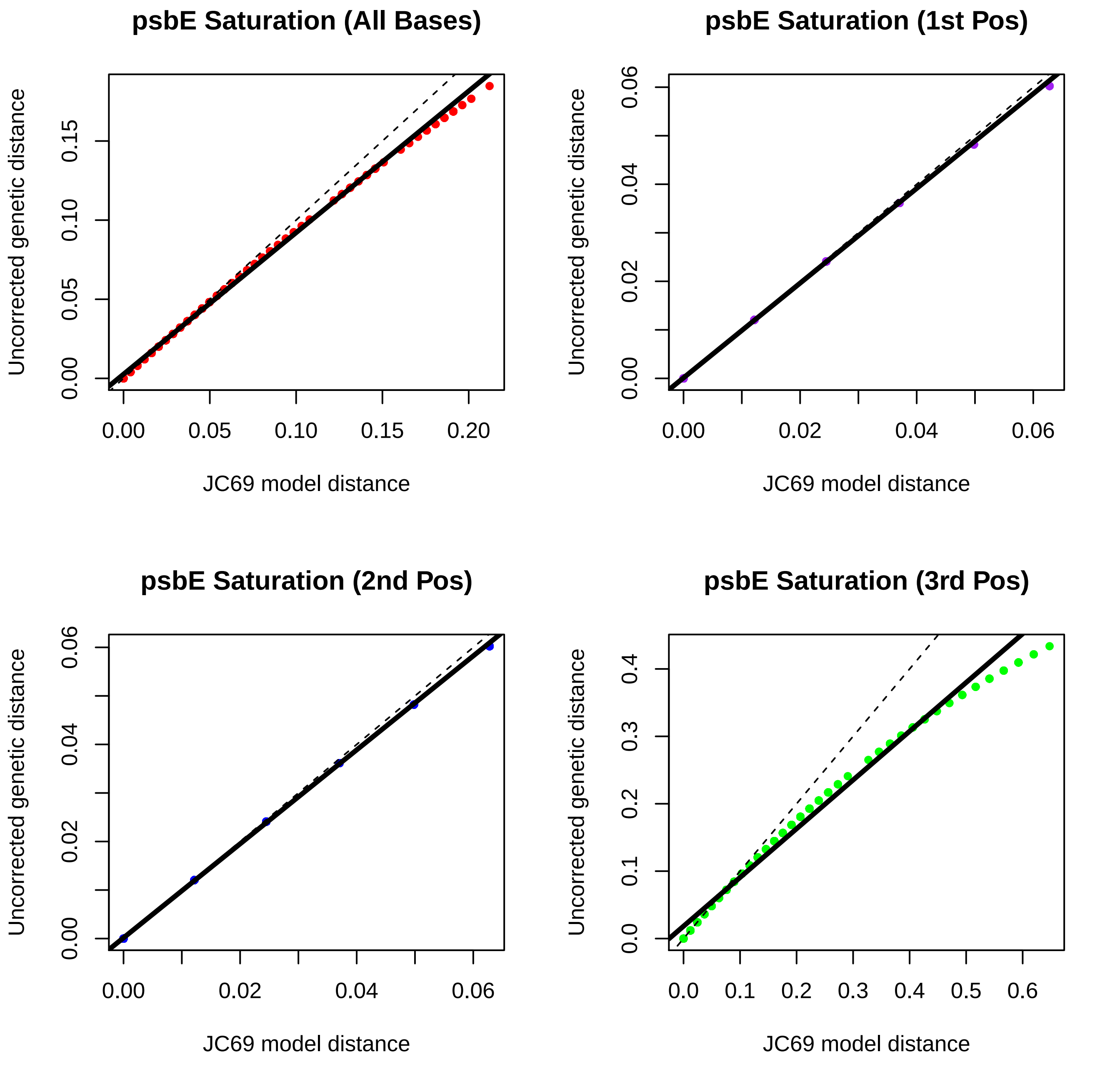
psbE Saturation (All Bases)
Uncorrected genetic distance
0.00
0.05
0.10
0.15
JC69 model distance
0.00
0.05
0.10
0.15
0.20
psbE Saturation (1st Pos)
Uncorrected genetic distance
0.00
0.02
0.04
0.06
JC69 model distance
0.00
0.02
0.04
0.06
psbE Saturation (2nd Pos)
Uncorrected genetic distance
0.00
0.02
0.04
0.06
JC69 model distance
0.00
0.02
0.04
0.06
psbE Saturation (3rd Pos)
Uncorrected genetic distance
0.0
0.1
0.2
0.3
0.4
JC69 model distance
0.0
0.1
0.2
0.3
0.4
0.5
0.6

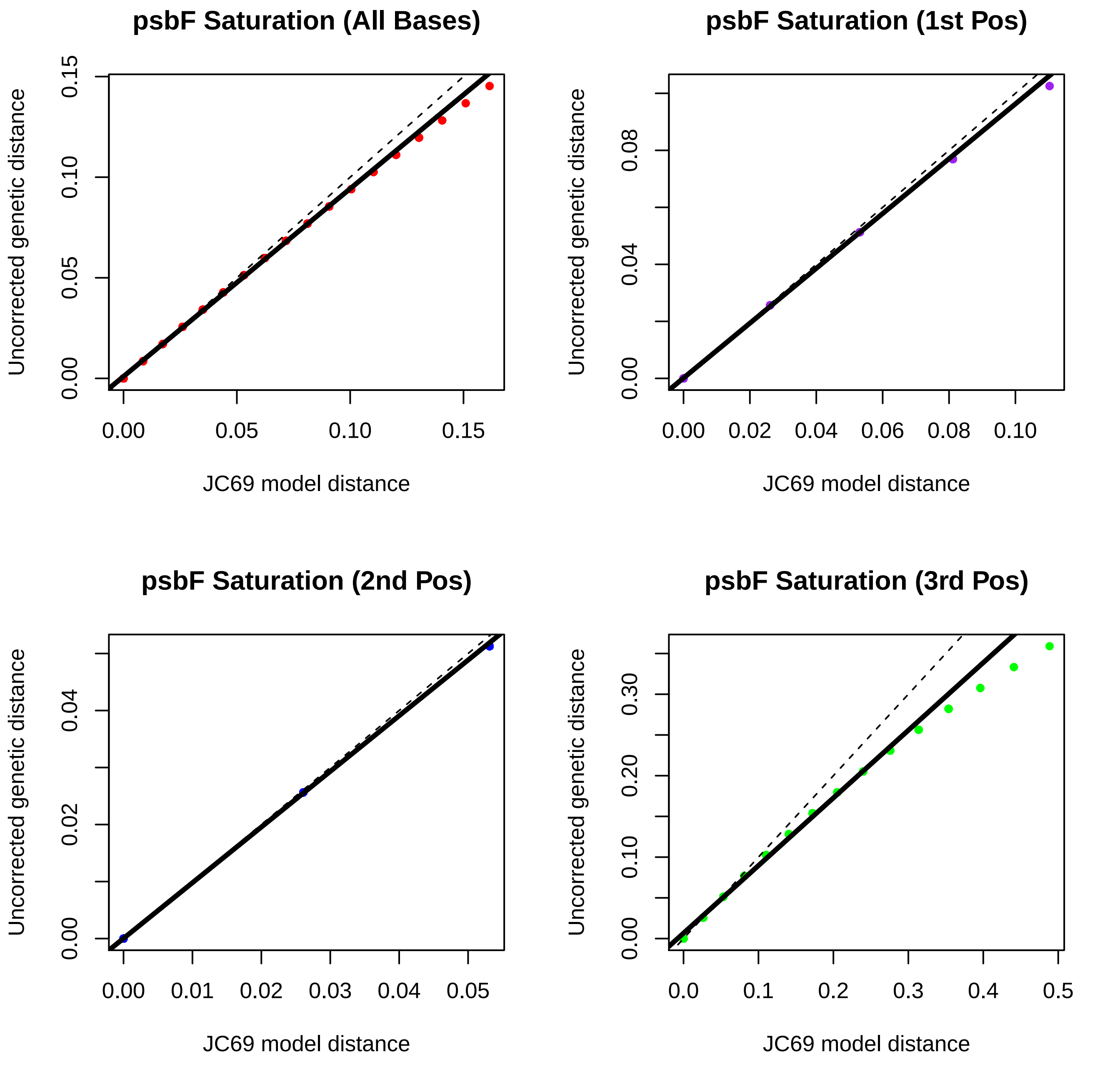
psbF Saturation (All Bases)
psbF Saturation (1st Pos)
psbF Saturation (2nd Pos)
psbF Saturation (3rd Pos)
Uncorrected genetic distance
JC69 model distance
0.00
0.05
0.10
0.15
0.02
0.04
0.06
0.08
0.01
0.03
0.0
0.1
0.2
0.3
0.4
0.5
0.20
0.30

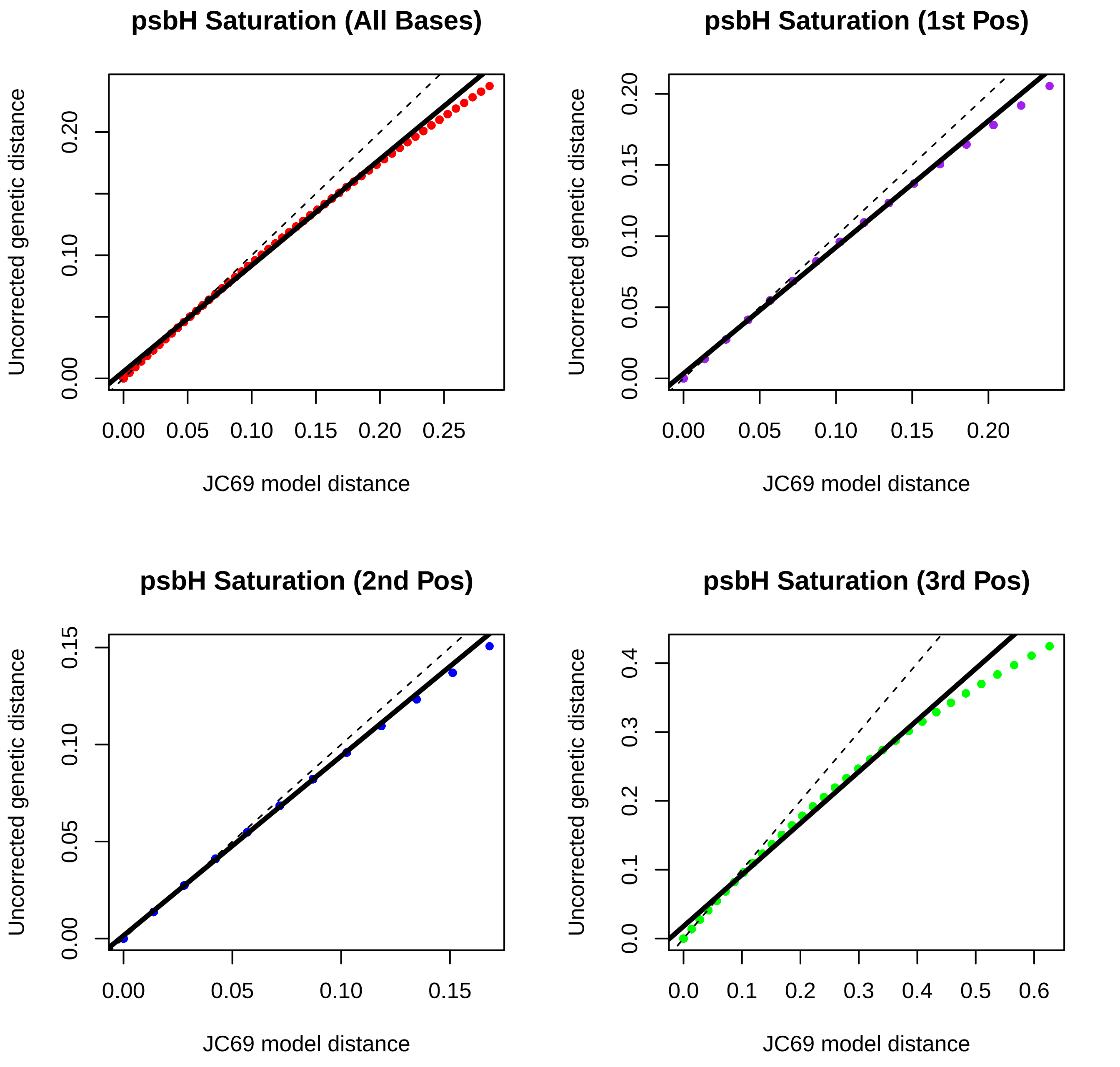
psbH Saturation (All Bases)
psbH Saturation (1st Pos)
psbH Saturation (2nd Pos)
psbH Saturation (3rd Pos)
Uncorrected genetic distance
JC69 model distance

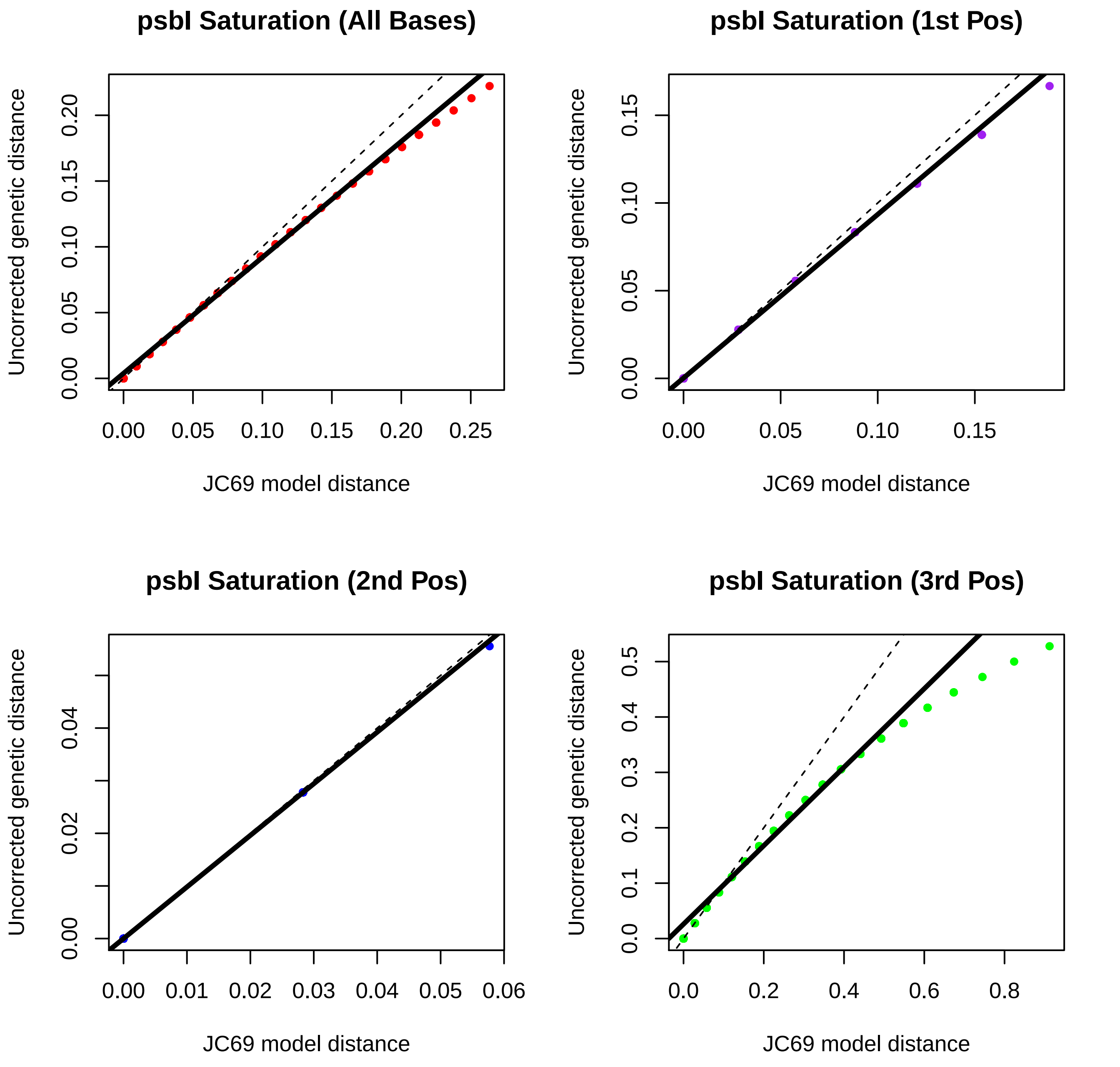
psbI Saturation (All Bases)
psbI Saturation (1st Pos)
psbI Saturation (2nd Pos)
psbI Saturation (3rd Pos)
Uncorrected genetic distance
JC69 model distance
0.00
0.05
0.10
0.15
0.20
0.25
0.01
0.02
0.03
0.04
0.06
0.0
0.1
0.2
0.3
0.4
0.5
0.6
0.8

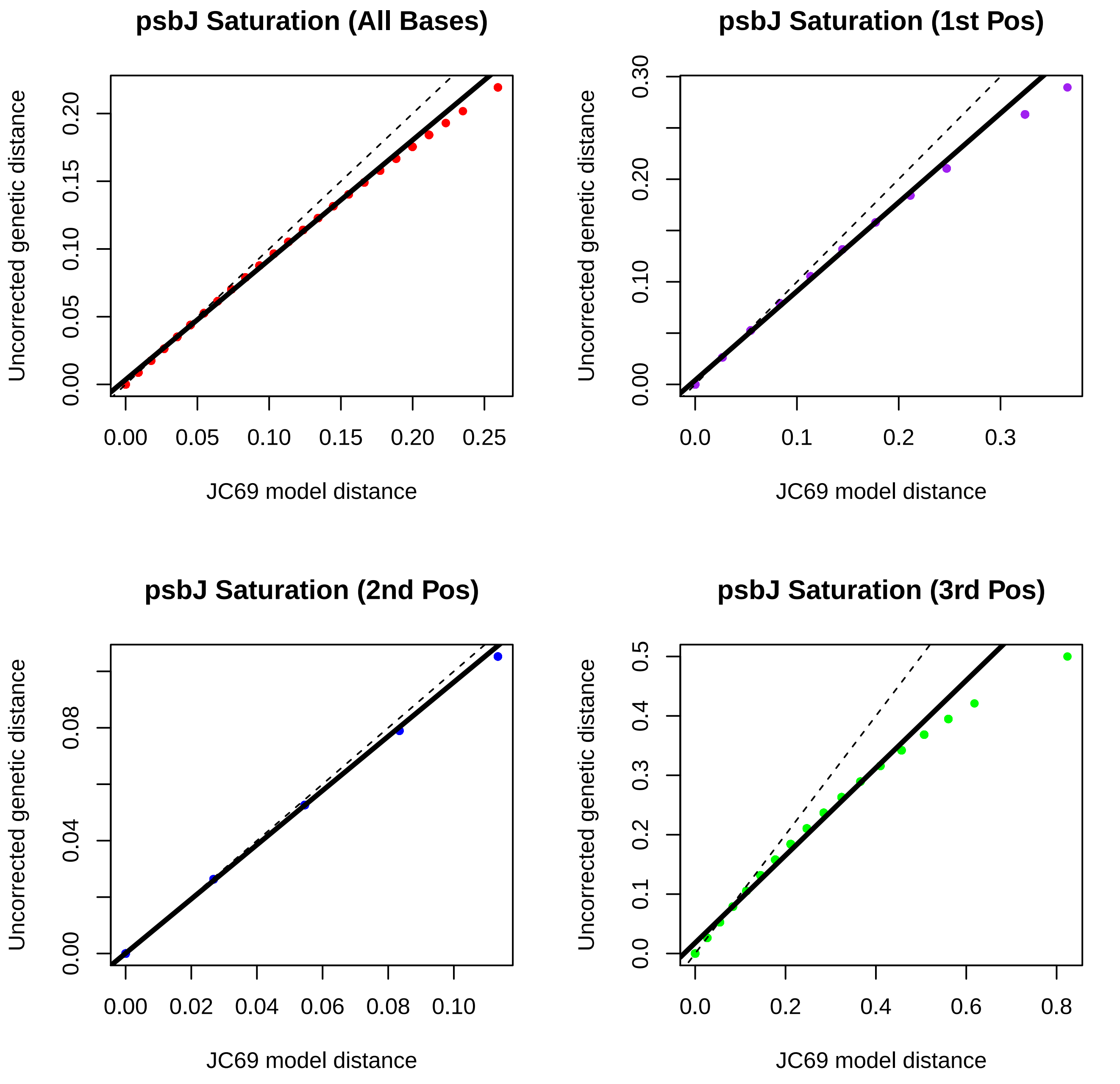

psbJ Saturation (All Bases)
Uncorrected genetic distance
JC69 model distance
0.00
0.05
0.10
0.15
0.20
0.25
psbJ Saturation (1st Pos)
0.0
0.1
0.2
0.3
0.00
0.10
0.20
0.30
psbJ Saturation (2nd Pos)
0.00
0.02
0.04
0.06
0.08
0.10
psbJ Saturation (3rd Pos)
0.0
0.2
0.4
0.6
0.8
0.0
0.1
0.2
0.3
0.4
0.5

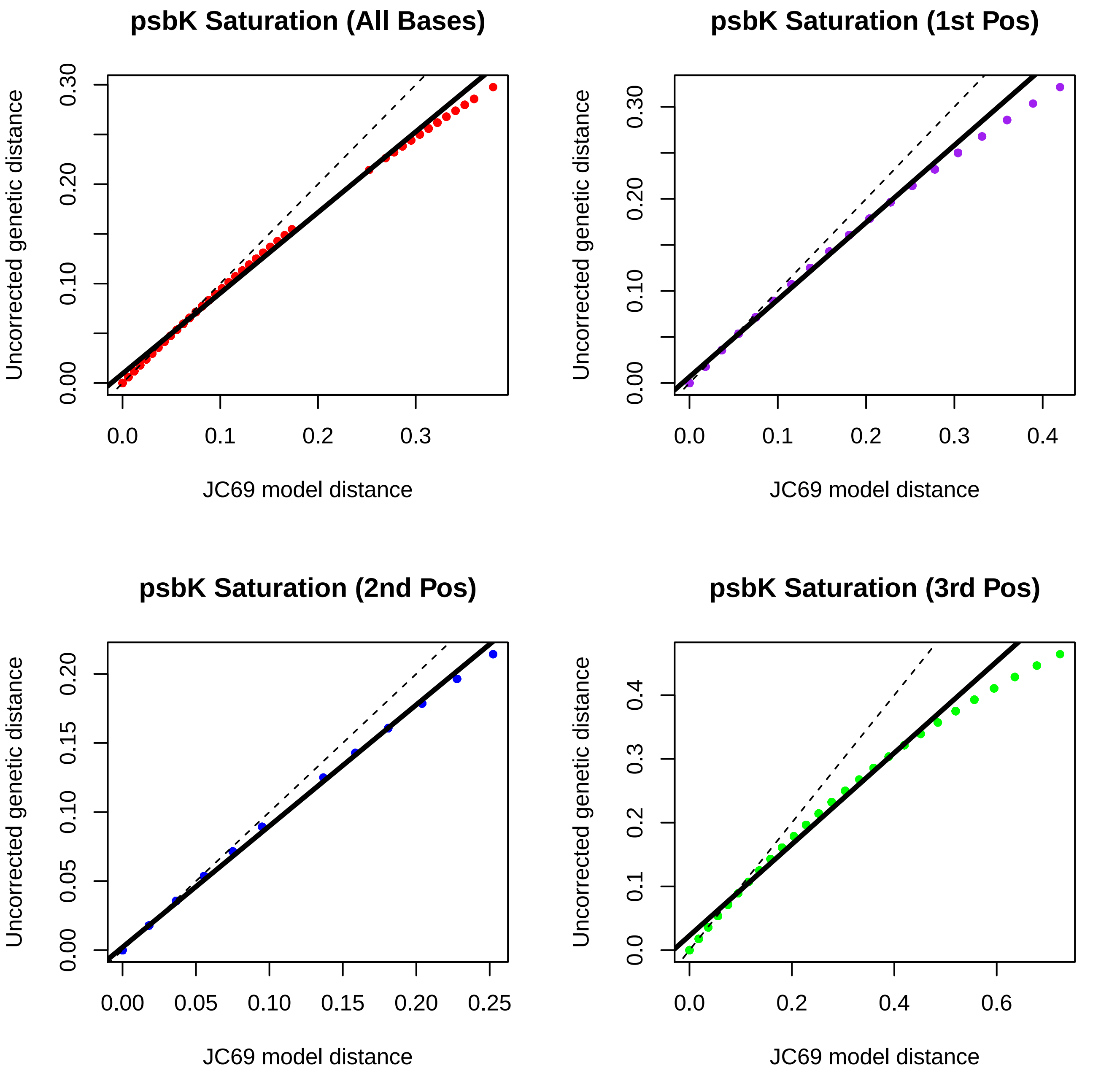
psbK Saturation (All Bases)
psbK Saturation (1st Pos)
psbK Saturation (2nd Pos)
psbK Saturation (3rd Pos)
Uncorrected genetic distance
JC69 model distance
0.00
0.10
0.20
0.30
0.0
0.1
0.2
0.3
0.4
0.05
0.15
0.25
0.6

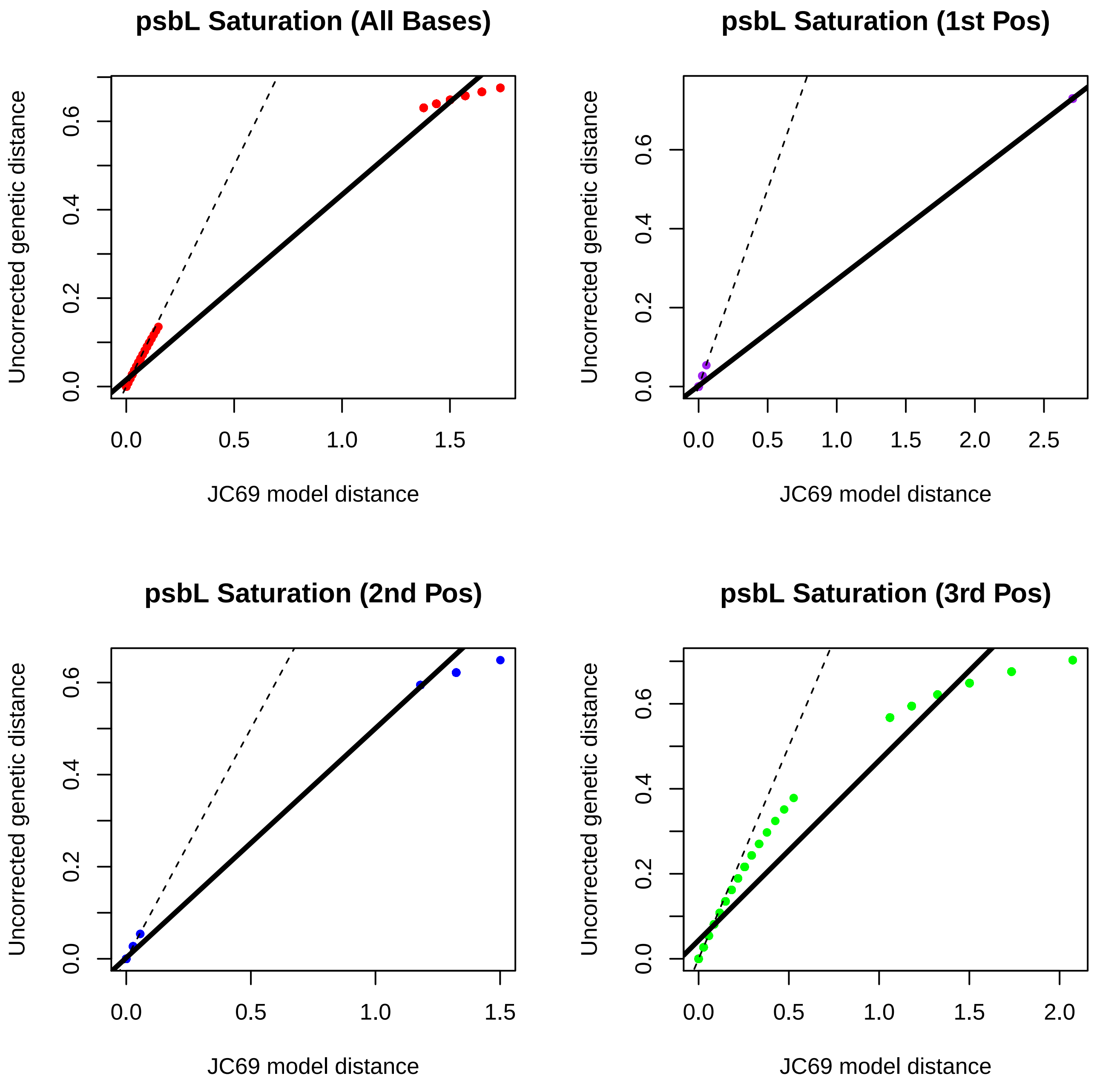
psbL Saturation (All Bases)
Uncorrected genetic distance
JC69 model distance
0.0
0.2
0.4
0.6
0.0
0.5
1.0
1.5
psbL Saturation (1st Pos)
Uncorrected genetic distance
JC69 model distance
0.0
0.2
0.4
0.6
0.0
0.5
1.0
1.5
2.0
2.5
psbL Saturation (2nd Pos)
Uncorrected genetic distance
JC69 model distance
0.0
0.2
0.4
0.6
0.0
0.5
1.0
1.5
psbL Saturation (3rd Pos)
Uncorrected genetic distance
JC69 model distance
0.0
0.2
0.4
0.6
0.0
0.5
1.0
1.5
2.0

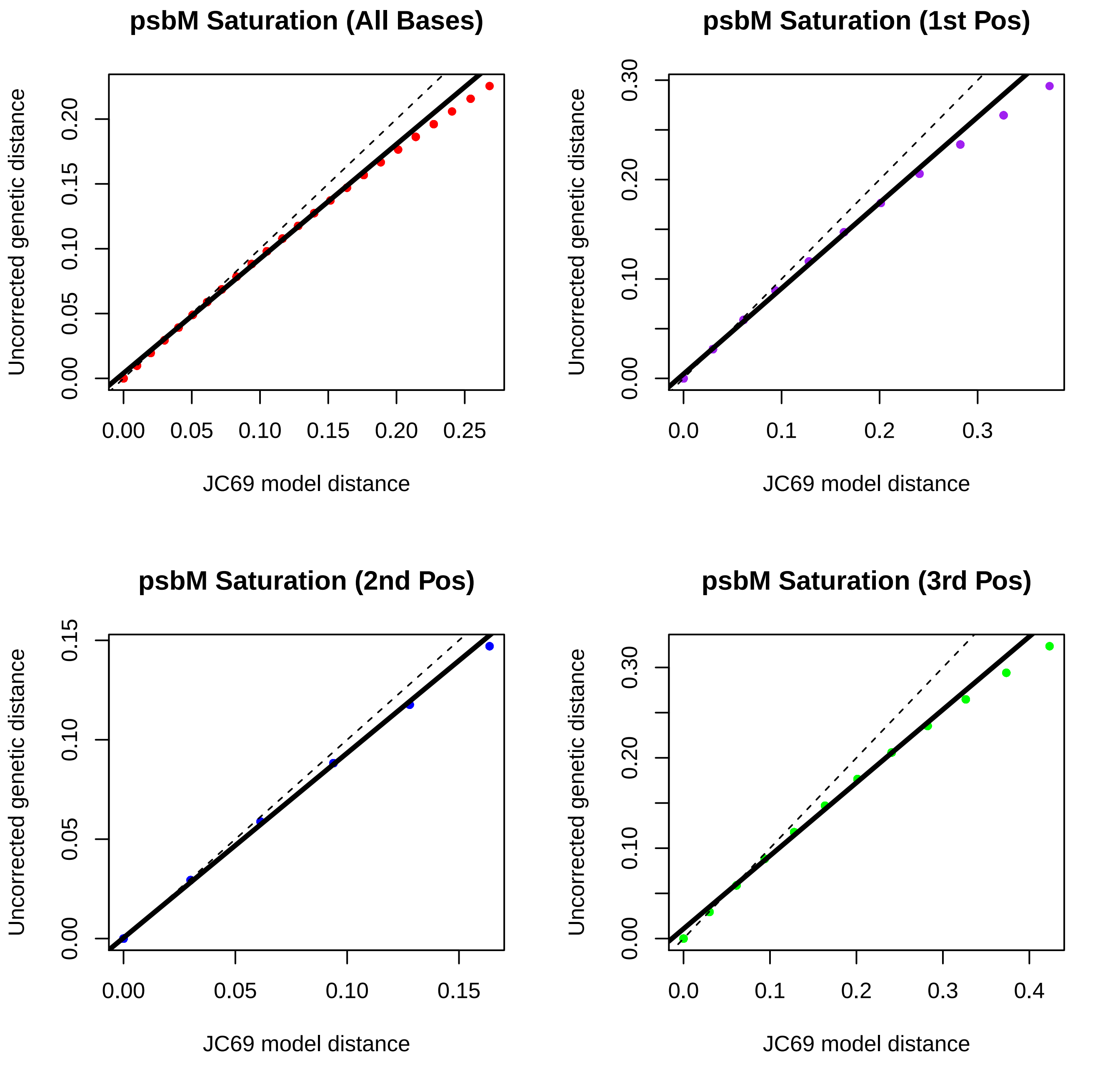
psbM Saturation (All Bases)
Uncorrected genetic distance
0.00
0.05
0.10
0.15
0.20
0.25
JC69 model distance
psbM Saturation (1st Pos)
0.0
0.1
0.2
0.3
0.00
0.10
0.20
0.30
psbM Saturation (2nd Pos)
0.15
psbM Saturation (3rd Pos)
0.4

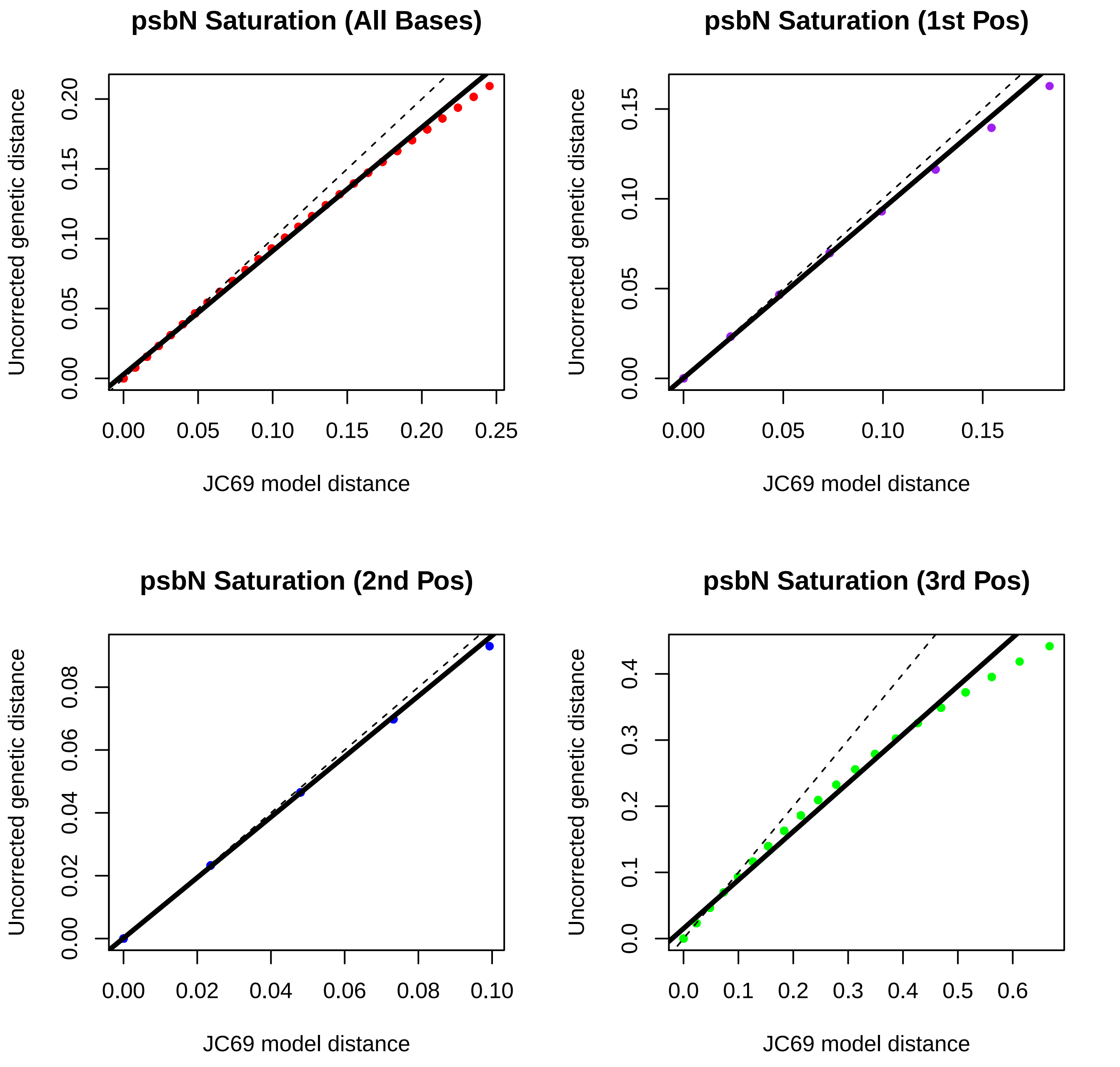
psbN Saturation (All Bases)
Uncorrected genetic distance
JC69 model distance
0.00
0.05
0.10
0.15
0.20
0.25
psbN Saturation (1st Pos)
Uncorrected genetic distance
JC69 model distance
0.00
0.05
0.10
0.15
psbN Saturation (2nd Pos)
Uncorrected genetic distance
JC69 model distance
0.00
0.02
0.04
0.06
0.08
0.10
psbN Saturation (3rd Pos)
Uncorrected genetic distance
JC69 model distance
0.0
0.1
0.2
0.3
0.4
0.5
0.6

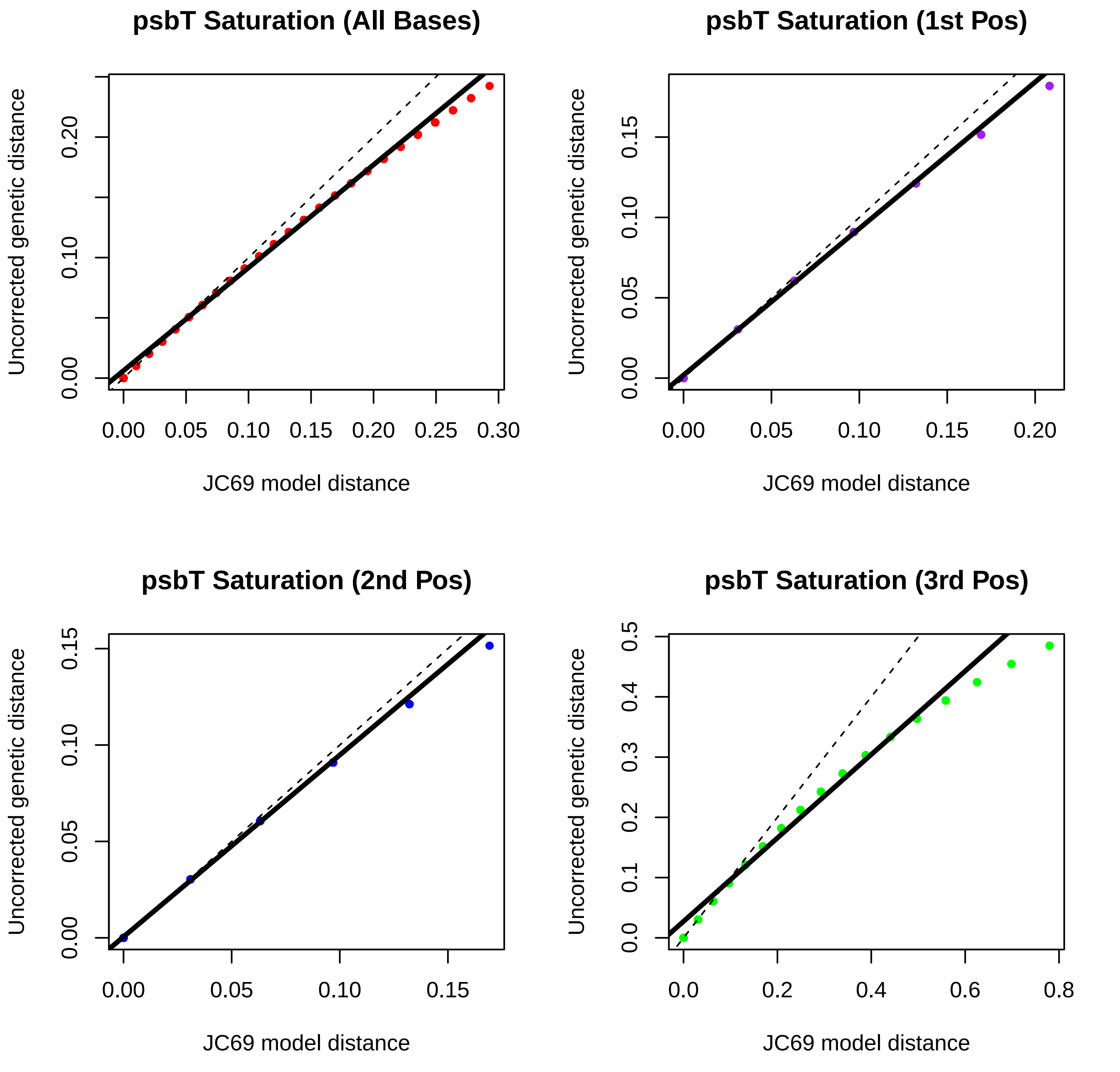
psbT Saturation (All Bases)
Uncorrected genetic distance
JC69 model distance
0.00
0.10
0.20
0.00
0.05
0.10
0.15
0.20
0.25
0.30
psbT Saturation (1st Pos)
Uncorrected genetic distance
JC69 model distance
0.00
0.05
0.10
0.15
0.00
0.05
0.10
0.15
0.20
psbT Saturation (2nd Pos)
Uncorrected genetic distance
JC69 model distance
0.00
0.05
0.10
0.15
0.00
0.05
0.10
0.15
psbT Saturation (3rd Pos)
Uncorrected genetic distance
JC69 model distance
0.0
0.1
0.2
0.3
0.4
0.5
0.0
0.2
0.4
0.6
0.8

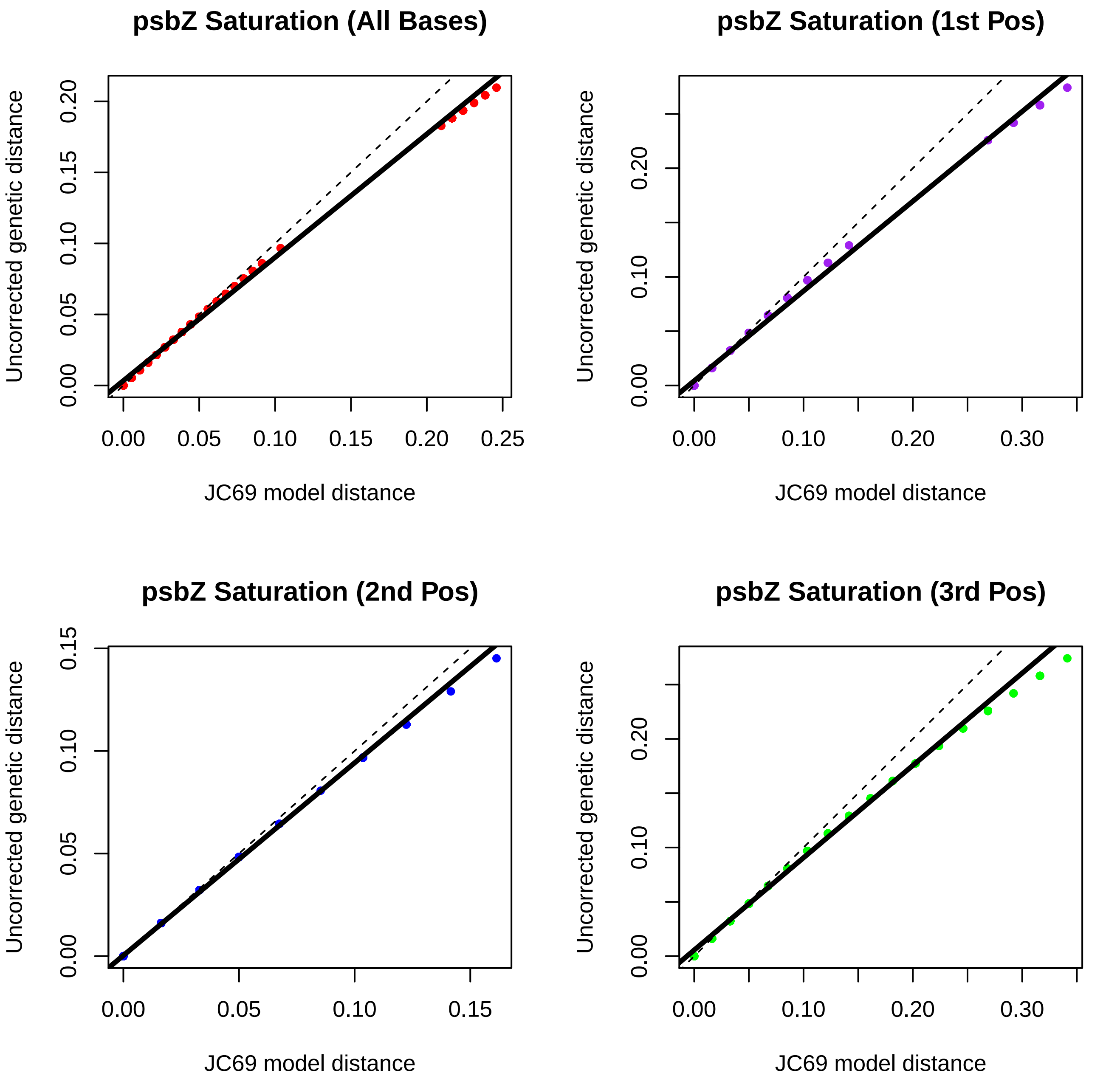
psbZ Saturation (All Bases)
Uncorrected genetic distance
0.00
0.05
0.10
0.15
0.20
0.00
0.05
0.10
0.15
0.20
0.25
JC69 model distance
psbZ Saturation (1st Pos)
Uncorrected genetic distance
0.00
0.10
0.20
0.00
0.10
0.20
0.30
JC69 model distance
psbZ Saturation (2nd Pos)
Uncorrected genetic distance
0.00
0.05
0.10
0.15
0.00
0.05
0.10
0.15
JC69 model distance
psbZ Saturation (3rd Pos)
Uncorrected genetic distance
0.00
0.10
0.20
0.00
0.10
0.20
0.30
JC69 model distance

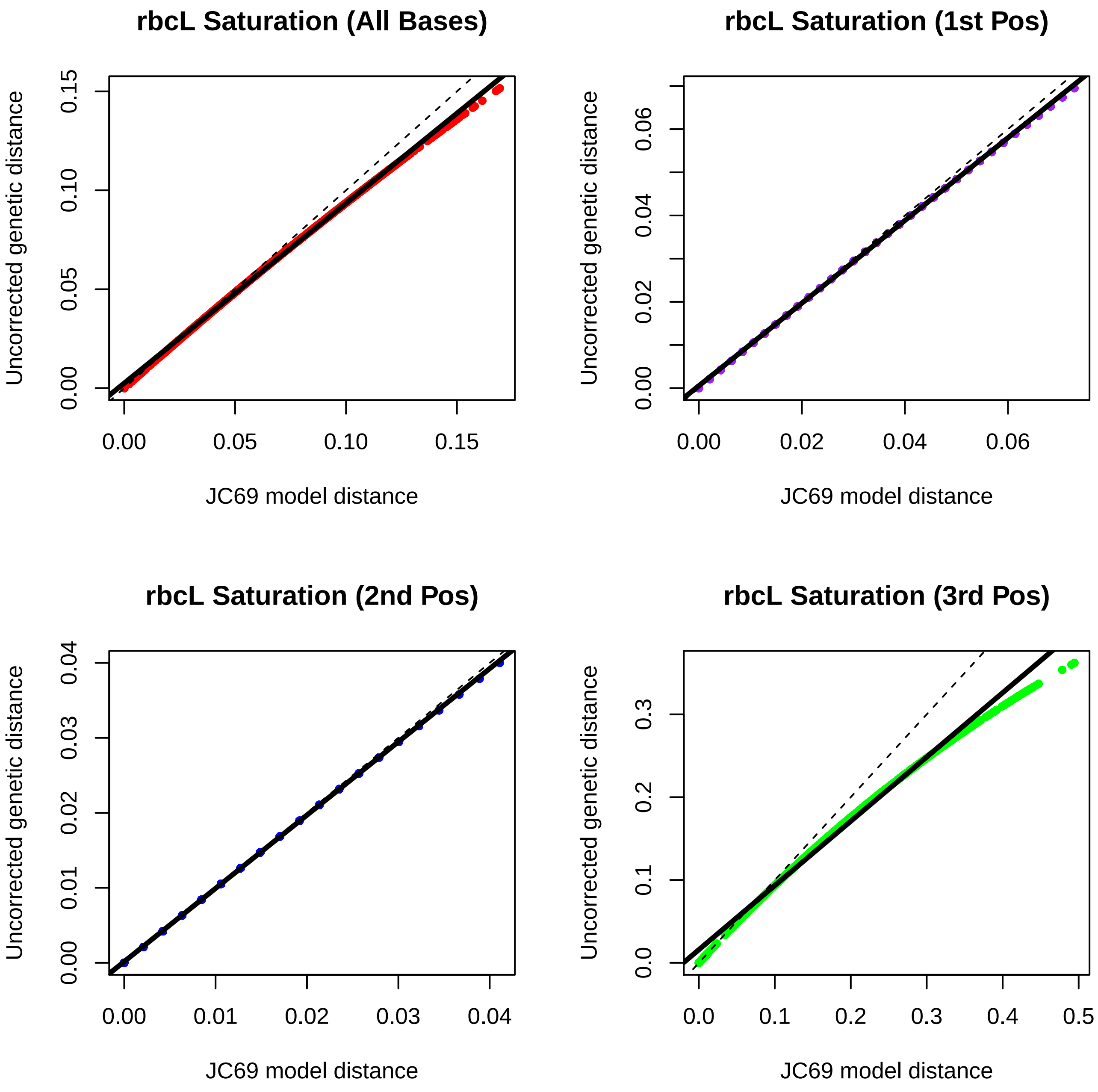
rbcL Saturation (All Bases)
Uncorrected genetic distance
0.00
0.05
0.10
0.15
JC69 model distance
0.00
0.05
0.10
0.15
rbcL Saturation (1st Pos)
Uncorrected genetic distance
0.00
0.02
0.04
0.06
JC69 model distance
0.00
0.02
0.04
0.06
rbcL Saturation (2nd Pos)
Uncorrected genetic distance
0.00
0.01
0.02
0.03
0.04
JC69 model distance
0.00
0.01
0.02
0.03
0.04
rbcL Saturation (3rd Pos)
Uncorrected genetic distance
0.0
0.1
0.2
0.3
JC69 model distance
0.0
0.1
0.2
0.3
0.4
0.5

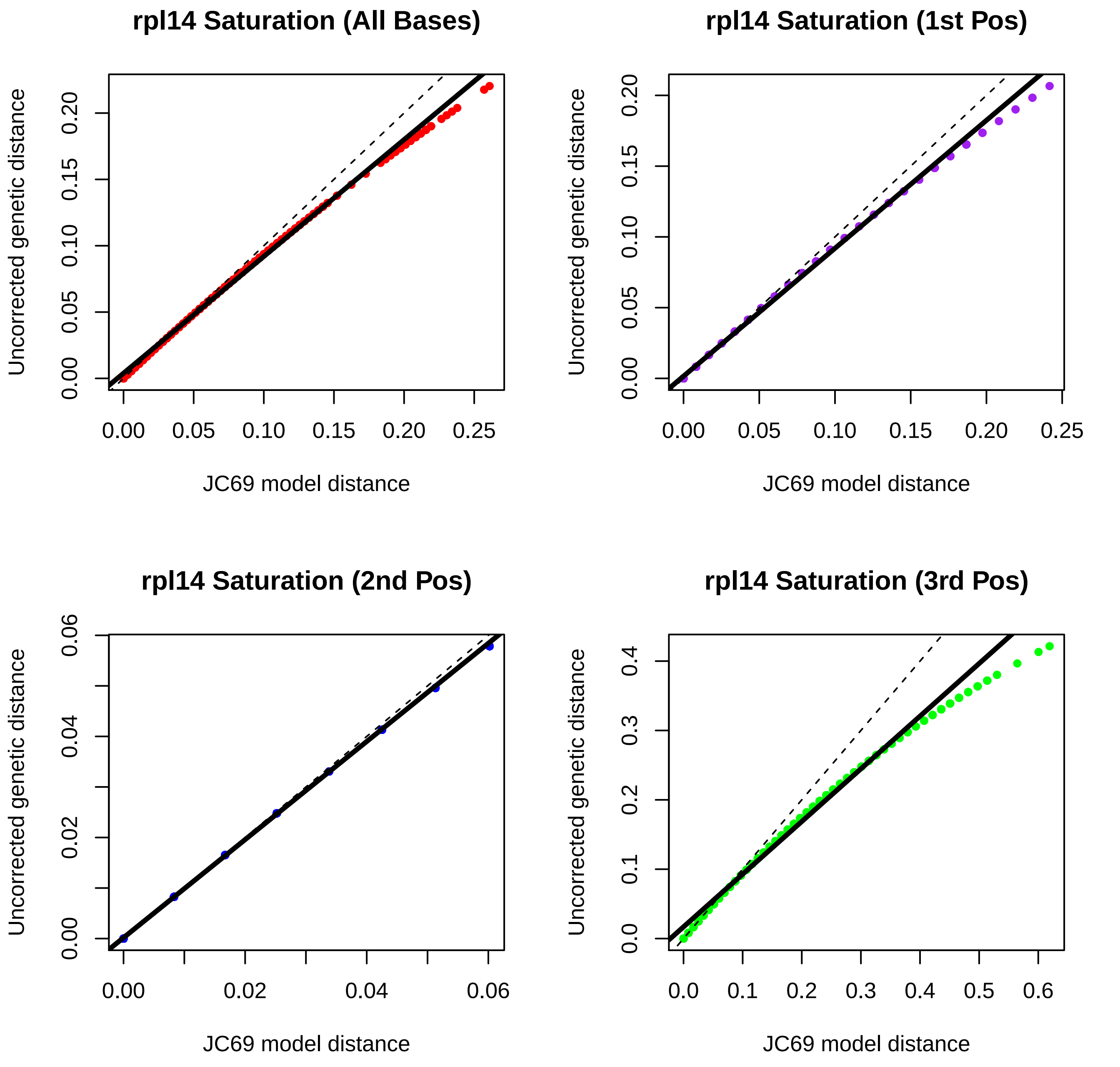
rpl14 Saturation (All Bases)
rpl14 Saturation (1st Pos)
rpl14 Saturation (2nd Pos)
rpl14 Saturation (3rd Pos)
Uncorrected genetic distance
JC69 model distance
0.00
0.05
0.10
0.15
0.20
0.25
0.00
0.02
0.04
0.06
0.0
0.1
0.2
0.3
0.4
0.5
0.6

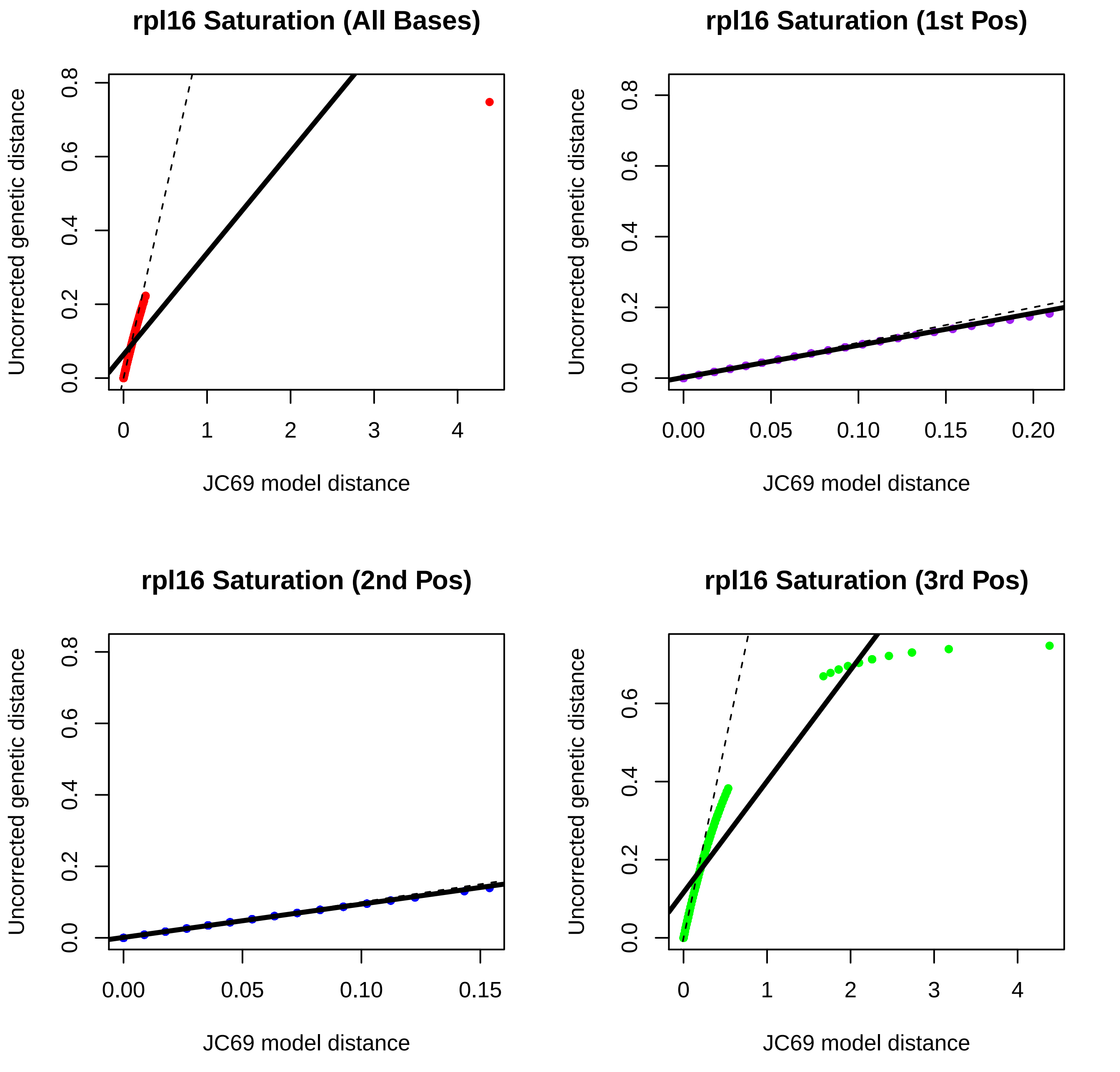
rpl16 Saturation (All Bases)
rpl16 Saturation (1st Pos)
rpl16 Saturation (2nd Pos)
rpl16 Saturation (3rd Pos)
Uncorrected genetic distance
JC69 model distance
0.0
0.2
0.4
0.6
0.8
0
1
2
3
4
0.00
0.05
0.10
0.15
0.20

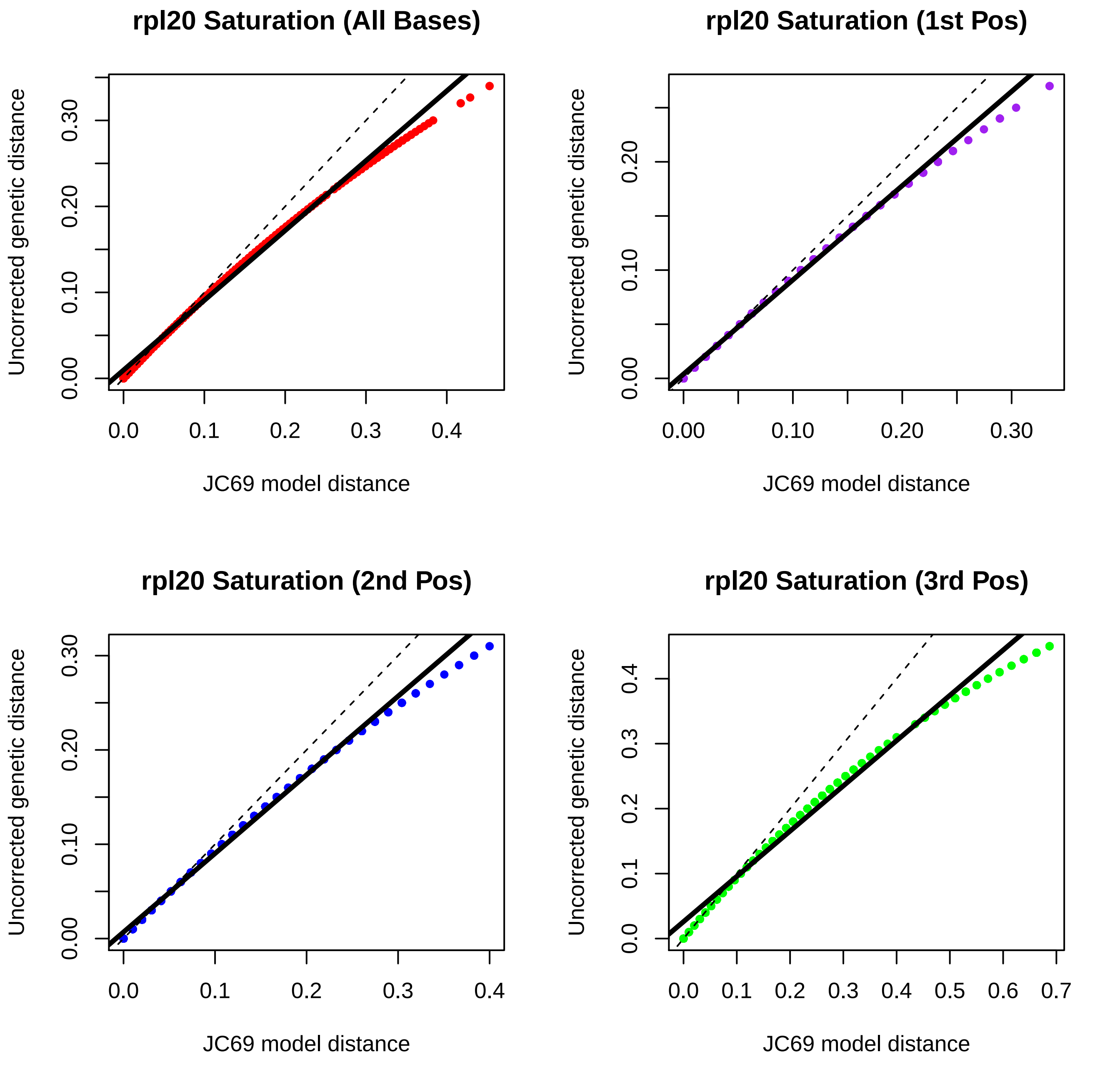
rpl20 Saturation (All Bases)
rpl20 Saturation (1st Pos)
rpl20 Saturation (2nd Pos)
rpl20 Saturation (3rd Pos)
Uncorrected genetic distance
JC69 model distance

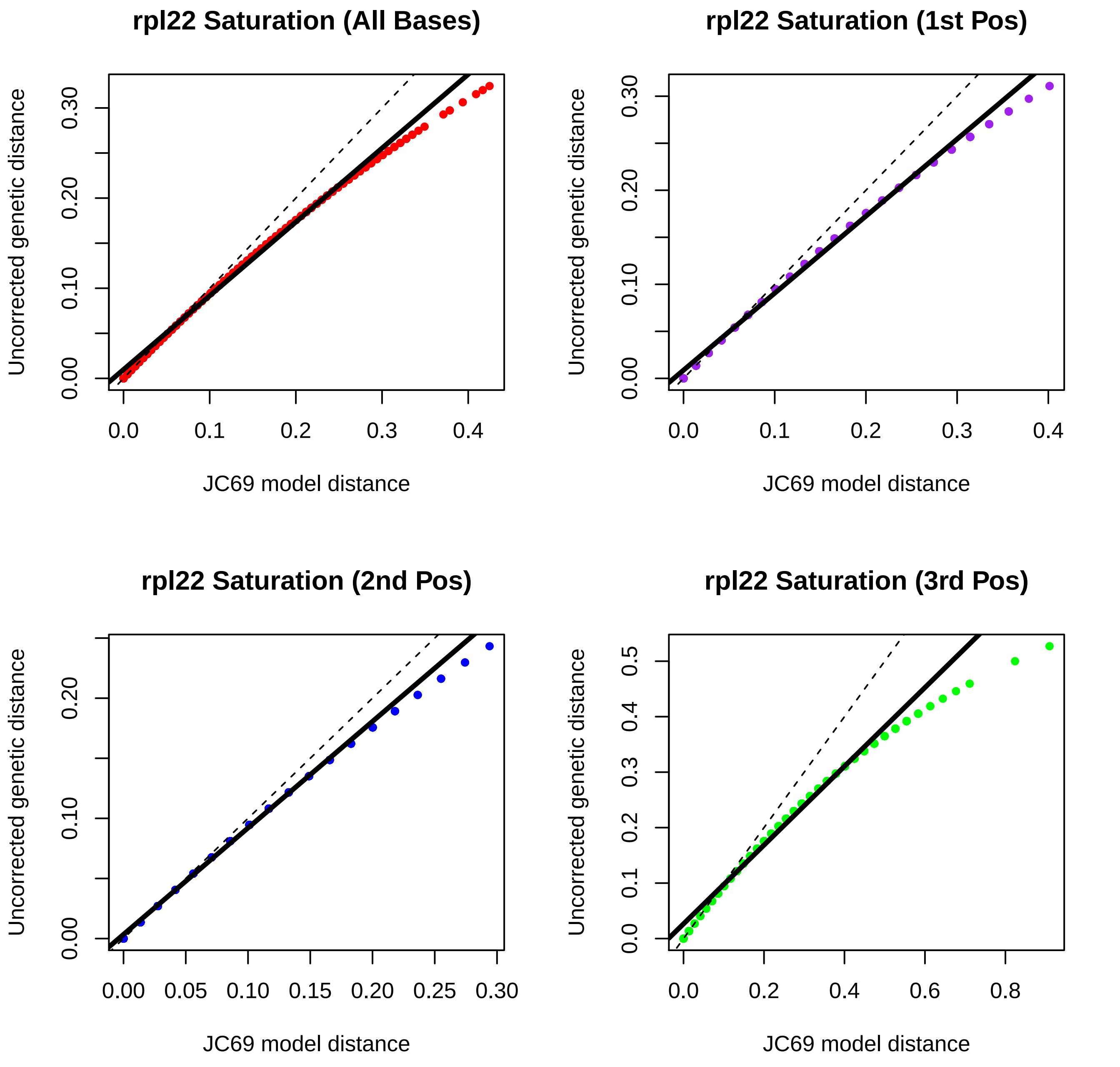
rpl22 Saturation (All Bases)
rpl22 Saturation (1st Pos)
rpl22 Saturation (2nd Pos)
rpl22 Saturation (3rd Pos)
Uncorrected genetic distance
JC69 model distance
0.00
0.10
0.20
0.30
0.0
0.1
0.2
0.3
0.4
0.05
0.15
0.25
0.5
0.6
0.8

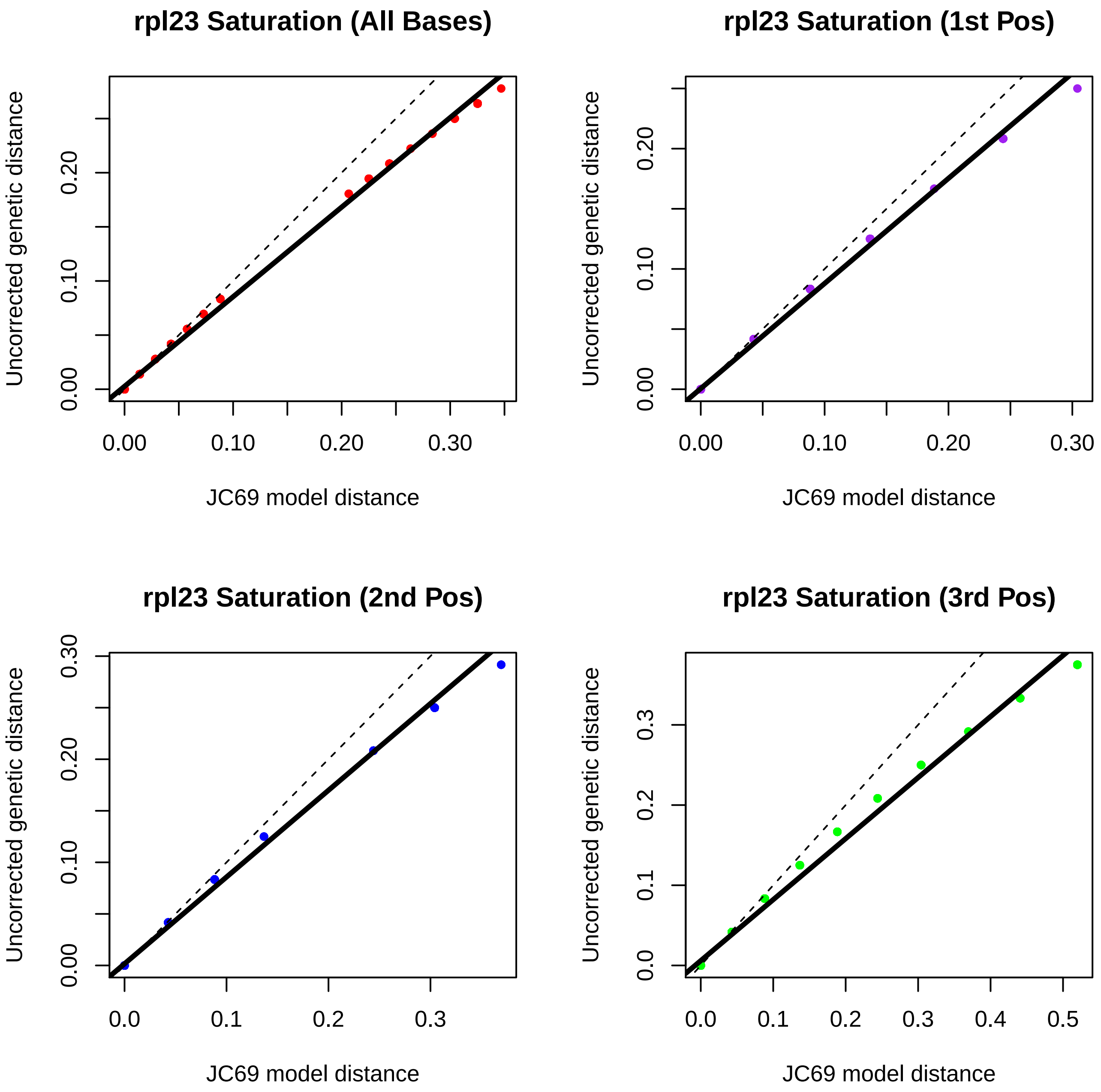
rpl23 Saturation (All Bases)
Uncorrected genetic distance
0.00
0.10
0.20
JC69 model distance
0.00
0.10
0.20
0.30
rpl23 Saturation (1st Pos)
Uncorrected genetic distance
0.00
0.10
0.20
JC69 model distance
0.00
0.10
0.20
0.30
rpl23 Saturation (2nd Pos)
Uncorrected genetic distance
0.00
0.10
0.20
0.30
JC69 model distance
0.0
0.1
0.2
0.3
rpl23 Saturation (3rd Pos)
Uncorrected genetic distance
0.0
0.1
0.2
0.3
JC69 model distance
0.0
0.1
0.2
0.3
0.4
0.5

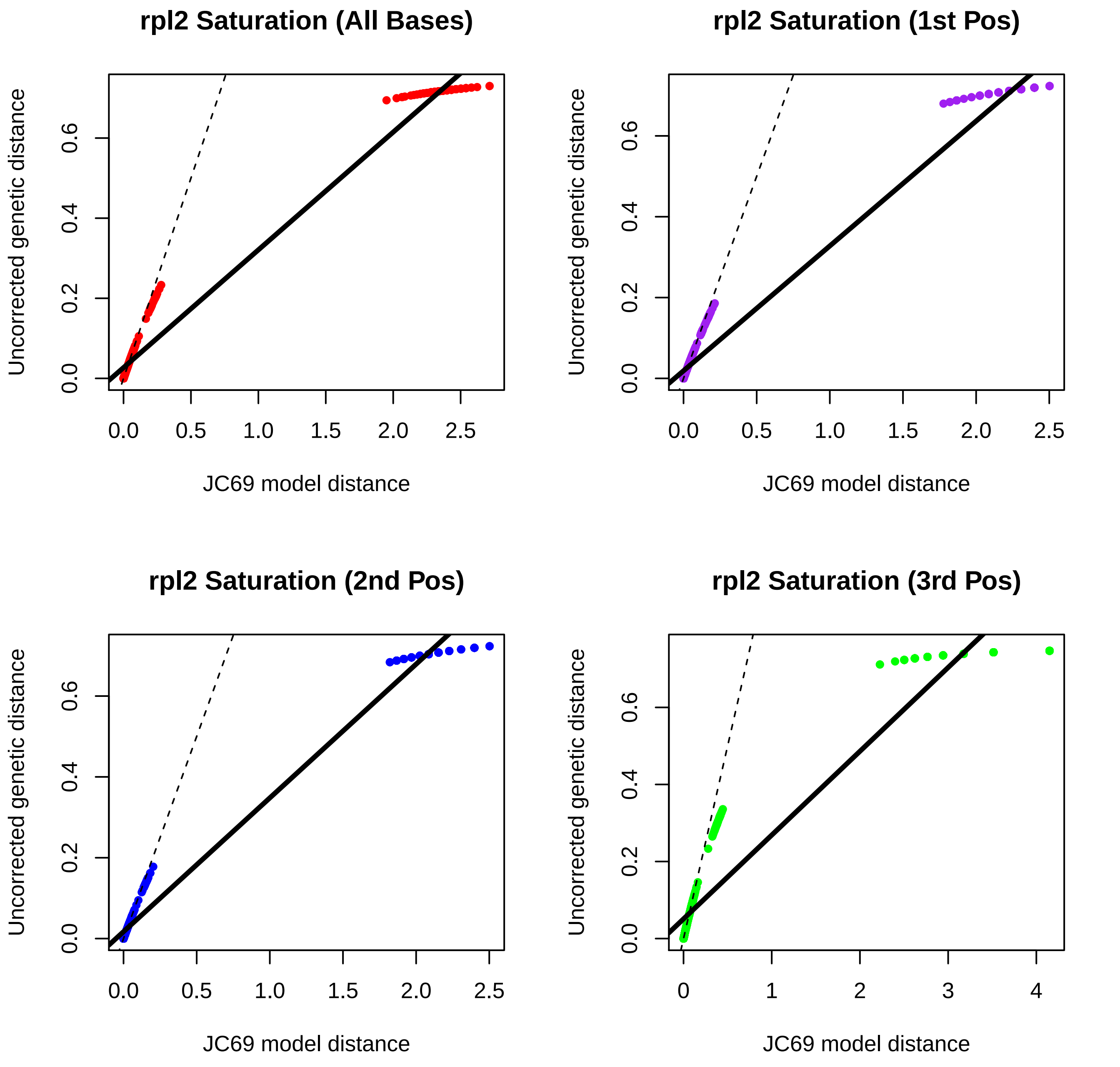

rpl2 Saturation (All Bases)
rpl2 Saturation (1st Pos)
rpl2 Saturation (2nd Pos)
rpl2 Saturation (3rd Pos)
Uncorrected genetic distance
JC69 model distance
0.0
0.2
0.4
0.6
0.5
1.0
1.5
2.0
2.5
0
1
2
3
4

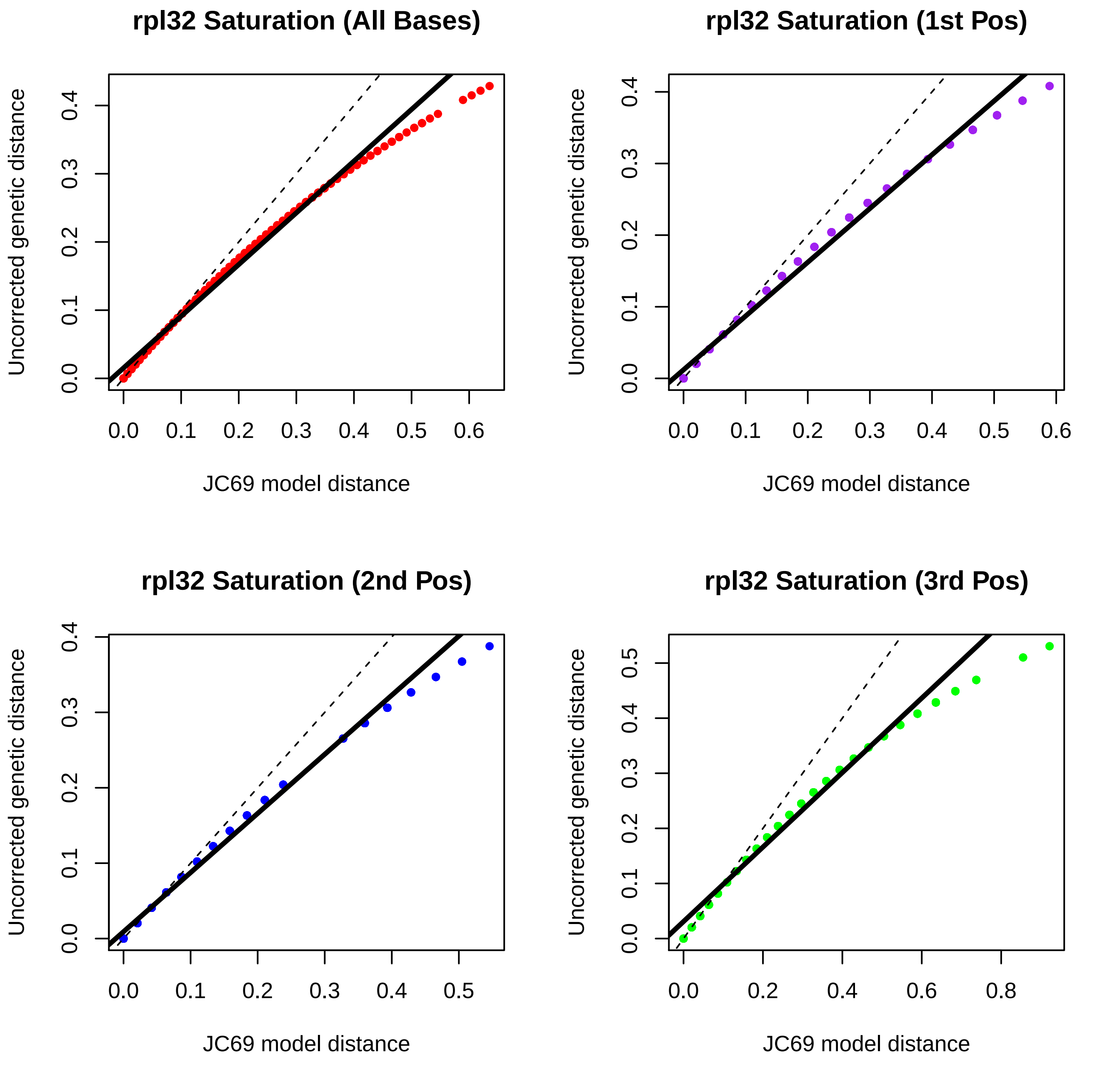

rpl32 Saturation (All Bases)
rpl32 Saturation (1st Pos)
rpl32 Saturation (2nd Pos)
rpl32 Saturation (3rd Pos)
Uncorrected genetic distance
JC69 model distance
0.0
0.1
0.2
0.3
0.4
0.5
0.6
0.8

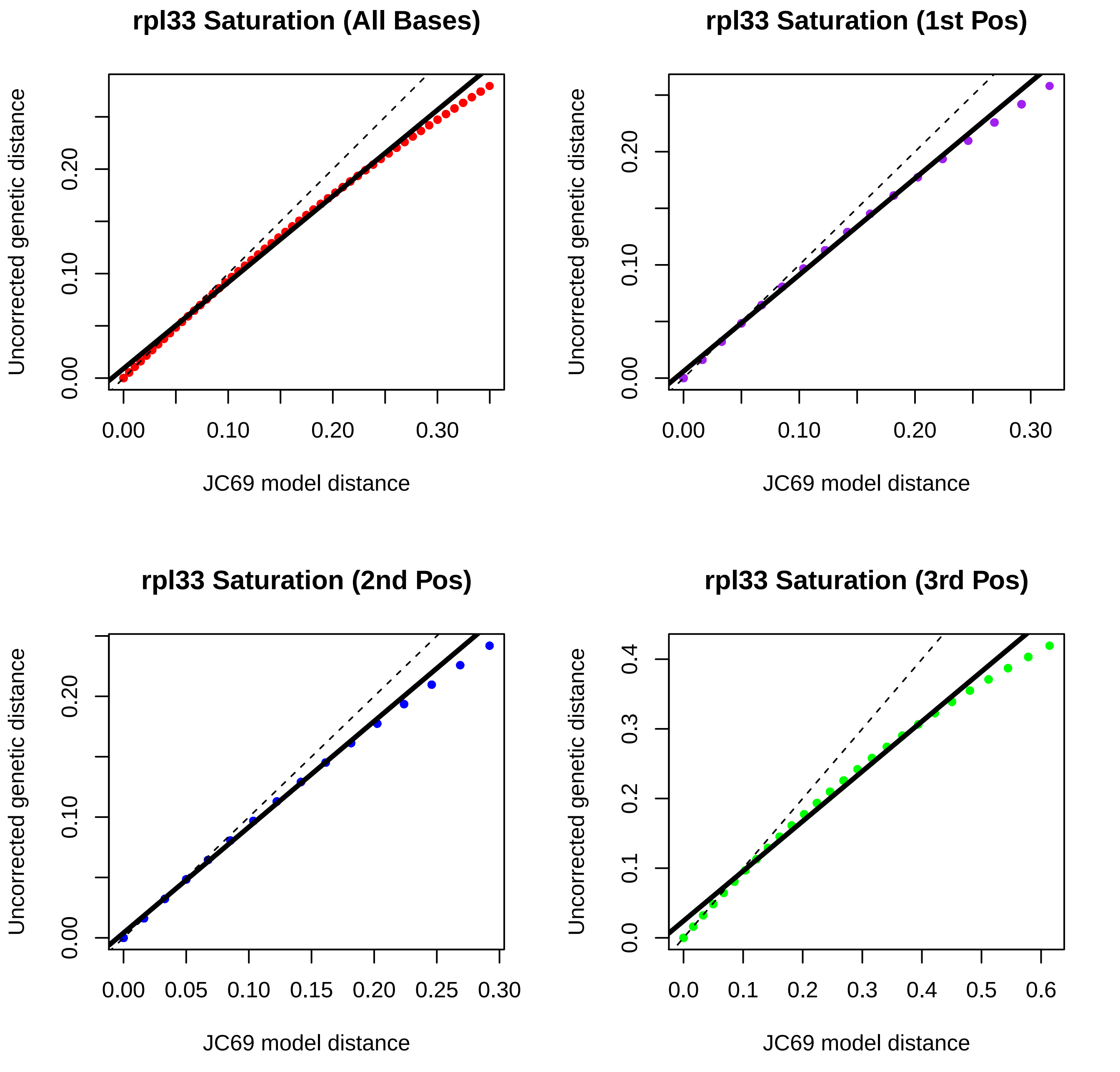
rpl33 Saturation (All Bases)
rpl33 Saturation (1st Pos)
rpl33 Saturation (2nd Pos)
rpl33 Saturation (3rd Pos)
Uncorrected genetic distance
JC69 model distance
0.00
0.10
0.20
0.30
0.05
0.15
0.25
0.0
0.1
0.2
0.3
0.4
0.5
0.6

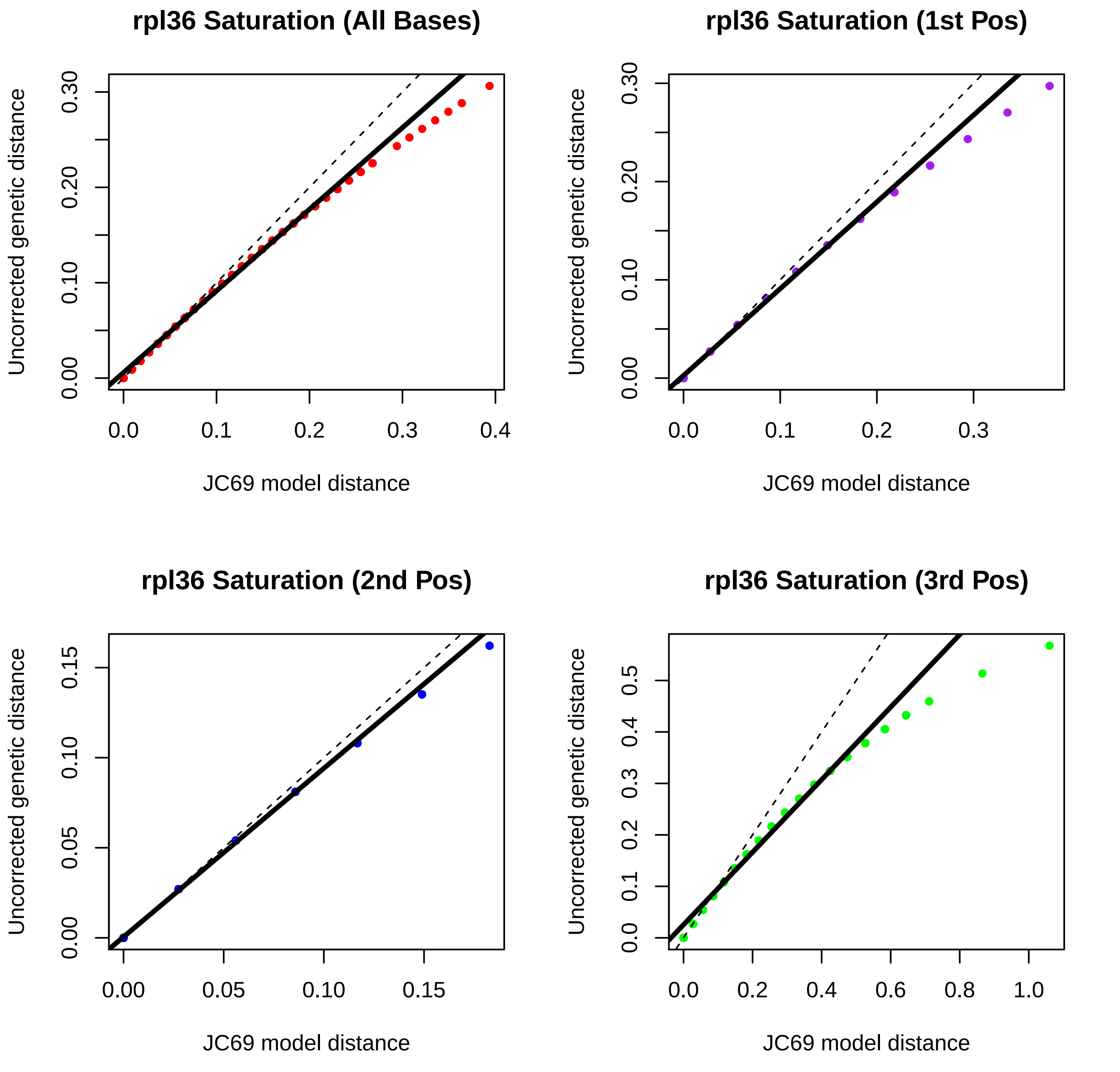

rpl36 Saturation (All Bases)
rpl36 Saturation (1st Pos)
rpl36 Saturation (2nd Pos)
rpl36 Saturation (3rd Pos)
Uncorrected genetic distance
JC69 model distance
0.00
0.10
0.20
0.30
0.0
0.1
0.2
0.3
0.4
0.05
0.15
0.5
0.6
0.8
1.0

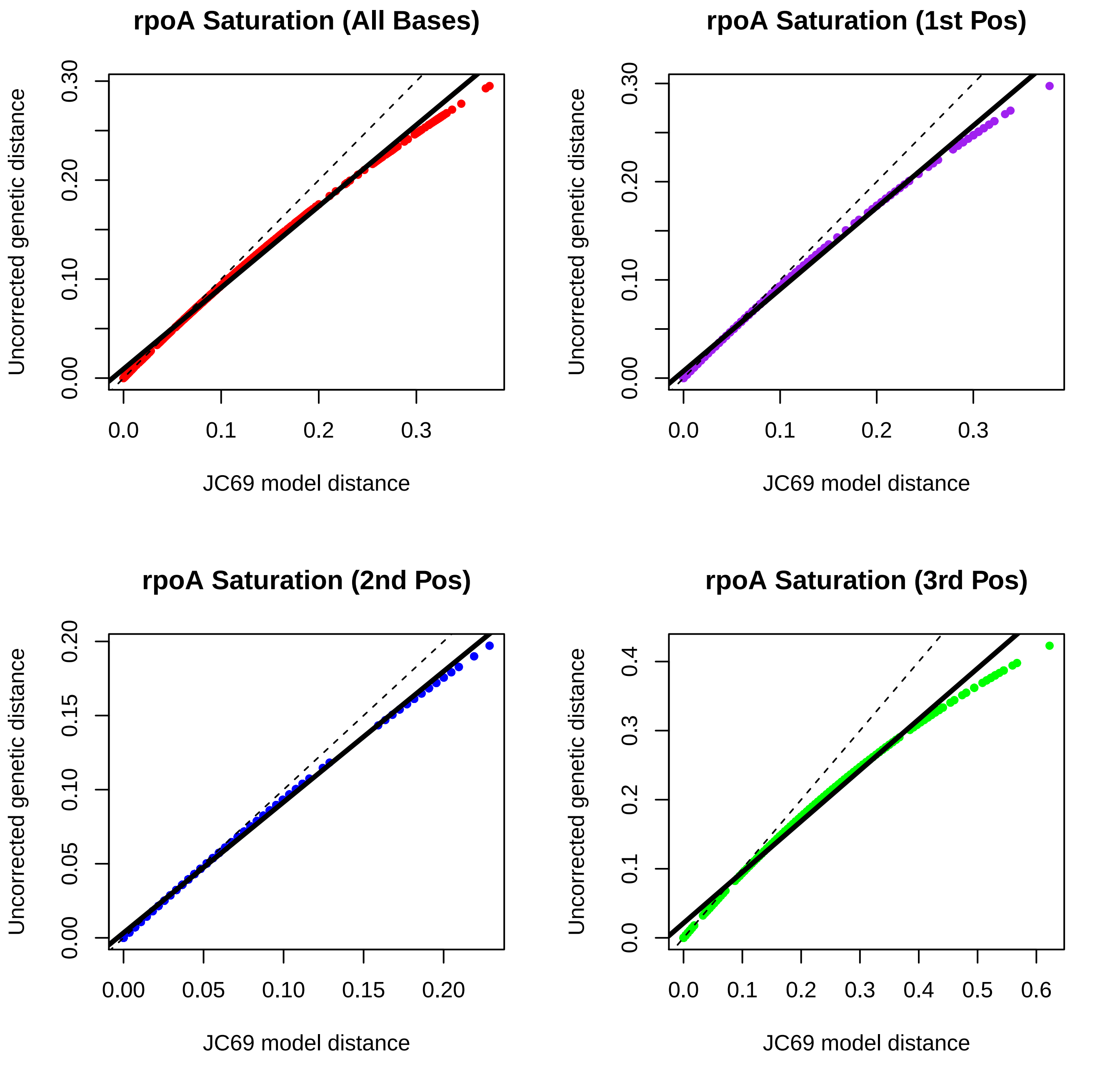

rpoA Saturation (All Bases)
rpoA Saturation (1st Pos)
rpoA Saturation (2nd Pos)
rpoA Saturation (3rd Pos)
Uncorrected genetic distance
JC69 model distance
0.00
0.10
0.20
0.30
0.0
0.1
0.2
0.3
0.05
0.15
0.4
0.5
0.6

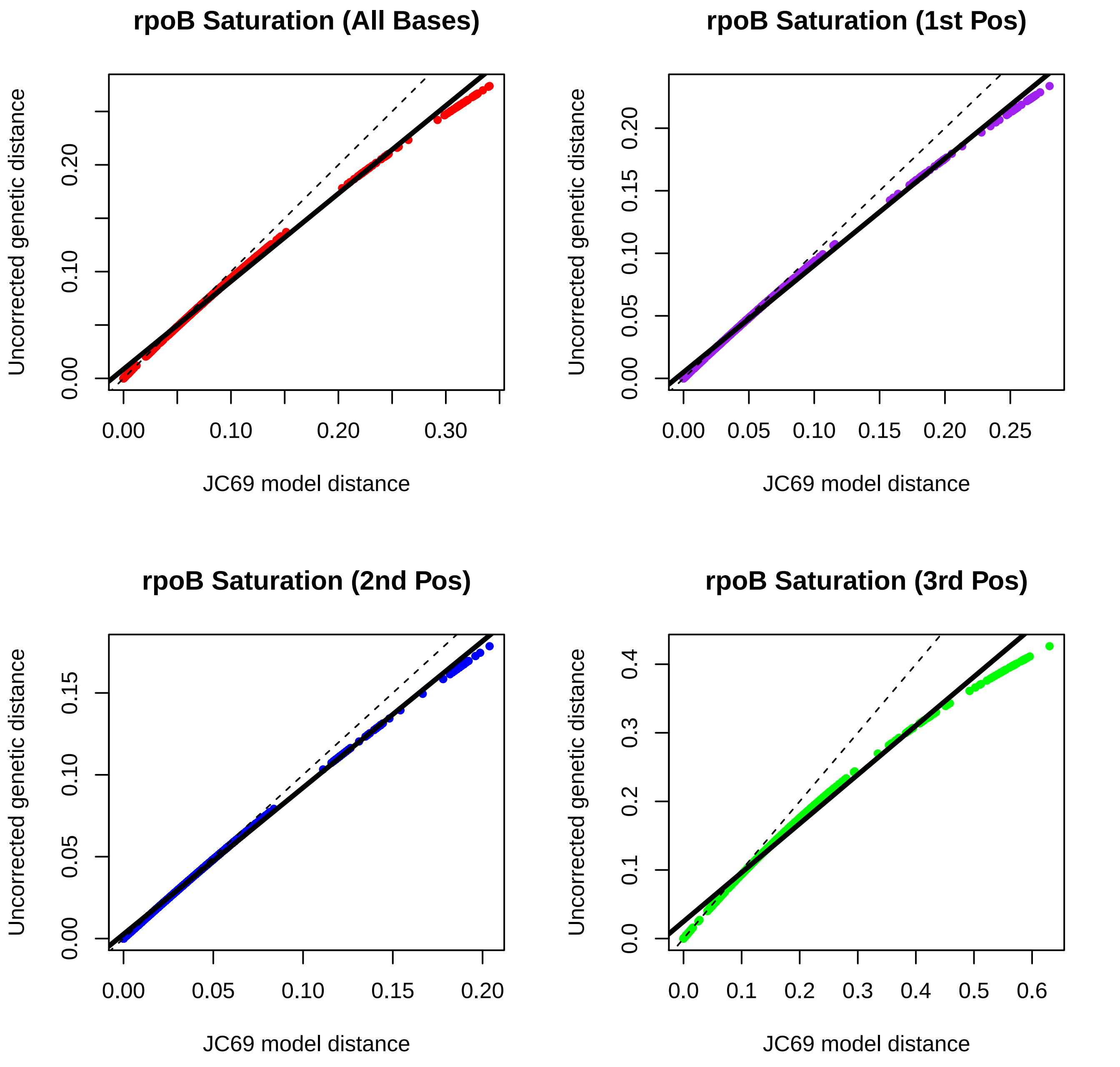
rpoB Saturation (All Bases)
rpoB Saturation (1st Pos)
rpoB Saturation (2nd Pos)
rpoB Saturation (3rd Pos)
Uncorrected genetic distance
JC69 model distance
0.00
0.10
0.20
0.30
0.05
0.15
0.25
0.0
0.1
0.2
0.3
0.4
0.5
0.6

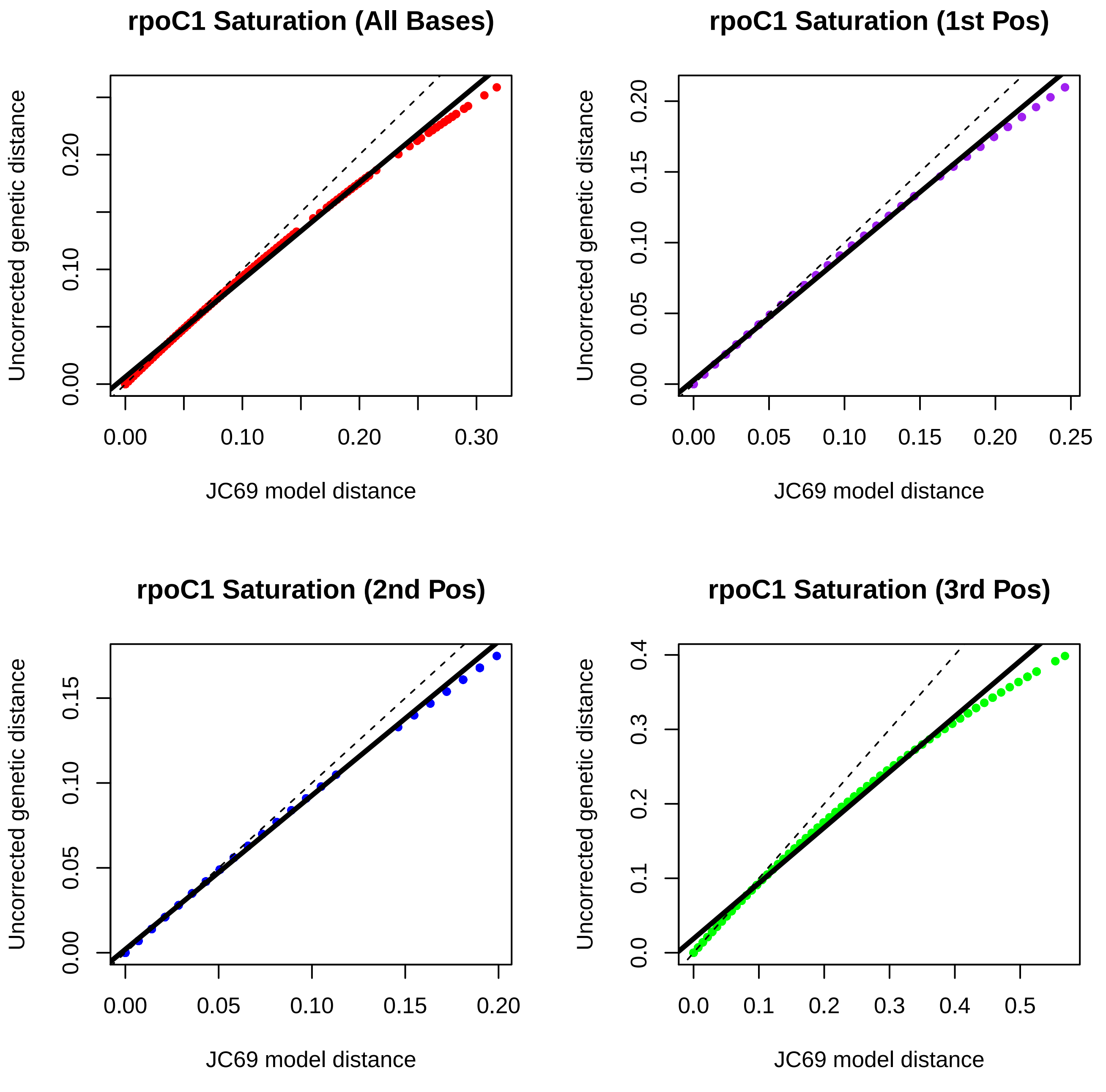
rpoC1 Saturation (All Bases)
Uncorrected genetic distance
JC69 model distance
0.00
0.10
0.20
0.00
0.10
0.20
0.30
rpoC1 Saturation (1st Pos)
Uncorrected genetic distance
JC69 model distance
0.00
0.05
0.10
0.15
0.20
0.00
0.05
0.10
0.15
0.20
0.25
rpoC1 Saturation (2nd Pos)
Uncorrected genetic distance
JC69 model distance
0.00
0.05
0.10
0.15
0.00
0.05
0.10
0.15
0.20
rpoC1 Saturation (3rd Pos)
Uncorrected genetic distance
JC69 model distance
0.0
0.1
0.2
0.3
0.4
0.0
0.1
0.2
0.3
0.4
0.5

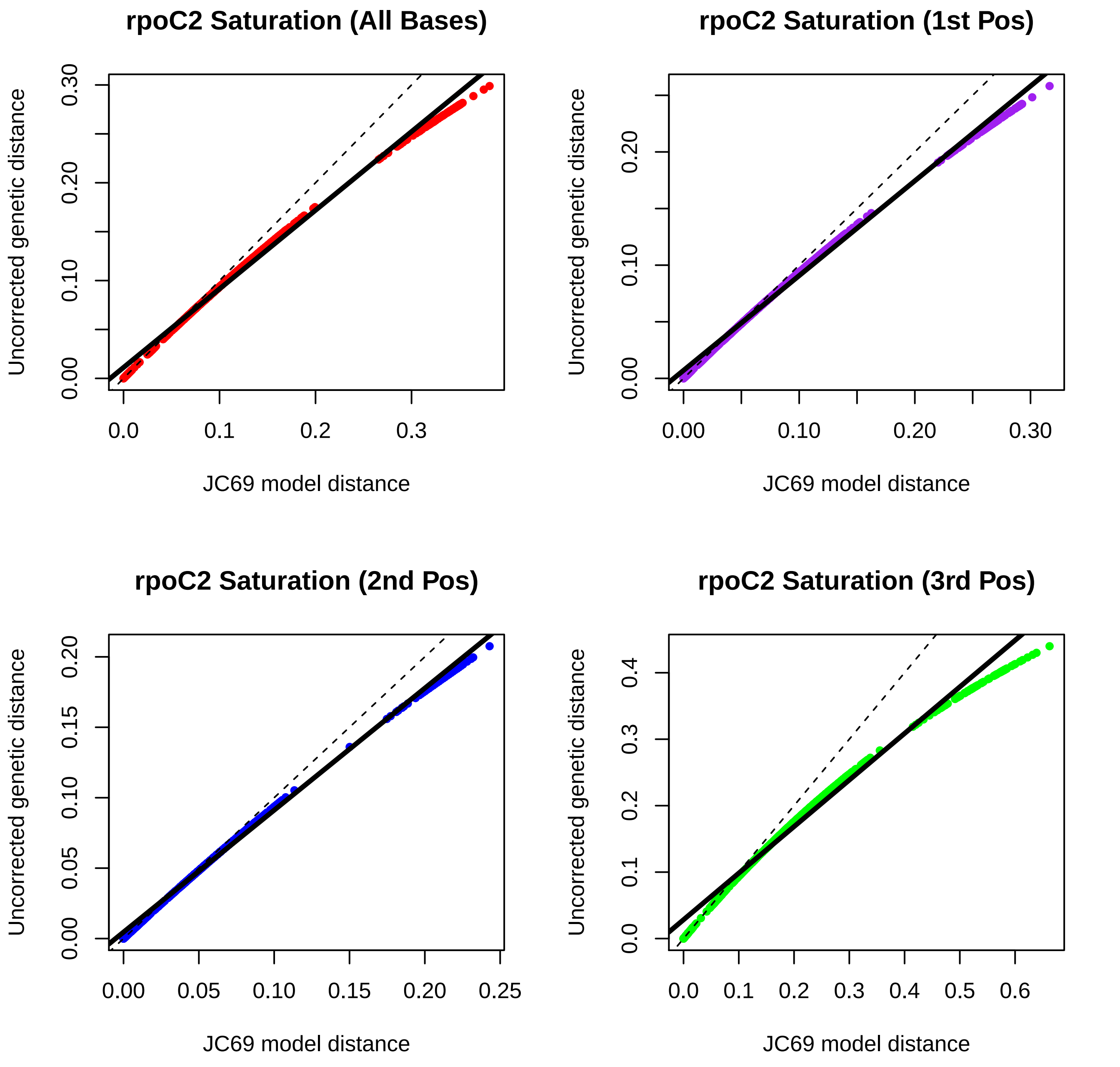
rpoC2 Saturation (All Bases)
Uncorrected genetic distance
0.00
0.10
0.20
0.30
0.0
0.1
0.2
0.3
JC69 model distance
rpoC2 Saturation (1st Pos)
Uncorrected genetic distance
0.00
0.10
0.20
0.00
0.10
0.20
0.30
JC69 model distance
rpoC2 Saturation (2nd Pos)
Uncorrected genetic distance
0.00
0.05
0.10
0.15
0.20
0.00
0.05
0.10
0.15
0.20
0.25
JC69 model distance
rpoC2 Saturation (3rd Pos)
Uncorrected genetic distance
0.0
0.1
0.2
0.3
0.4
0.0
0.1
0.2
0.3
0.4
0.5
0.6
JC69 model distance

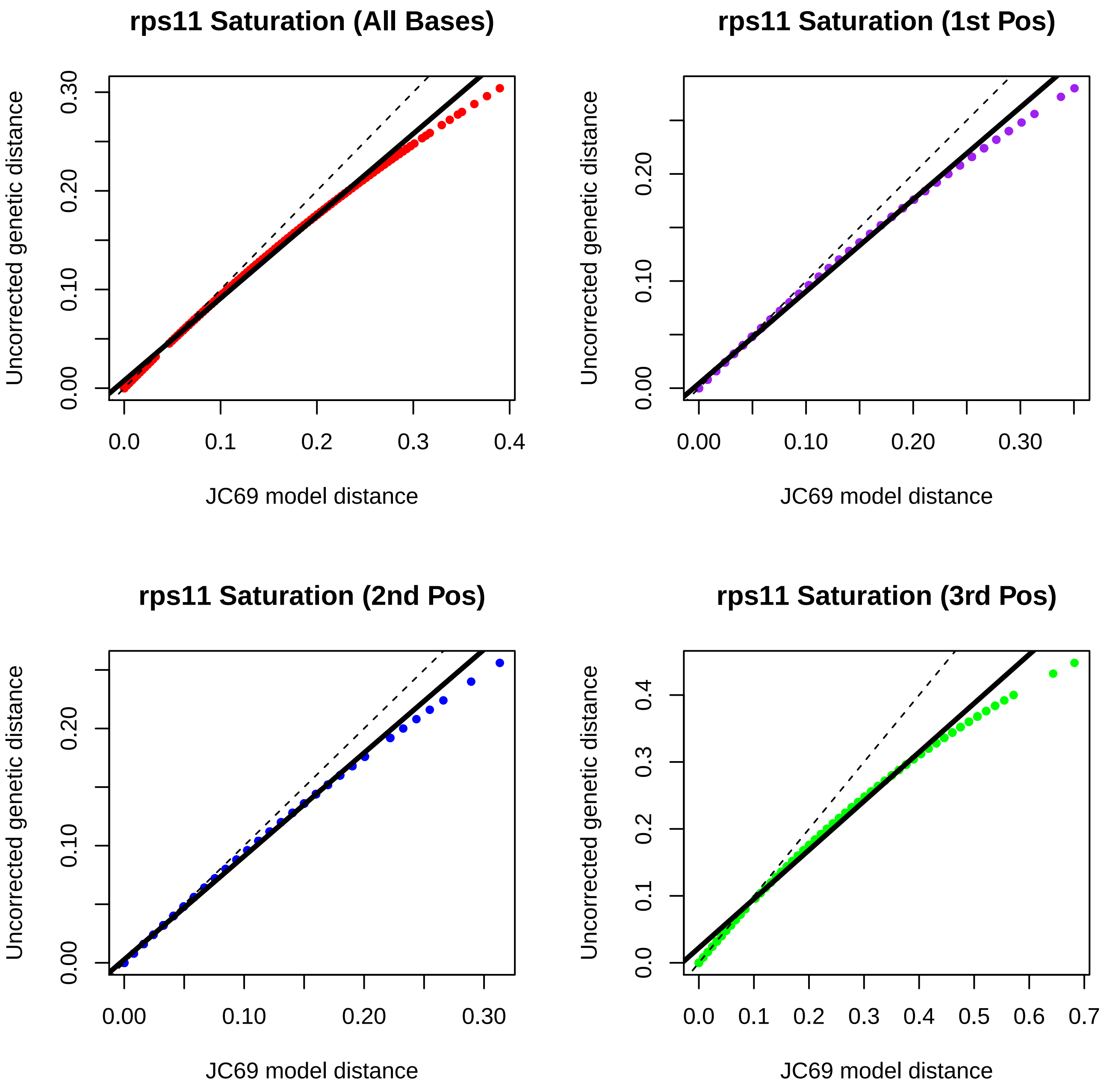
rps11 Saturation (All Bases)
rps11 Saturation (1st Pos)
rps11 Saturation (2nd Pos)
rps11 Saturation (3rd Pos)
Uncorrected genetic distance
JC69 model distance
0.00
0.10
0.20
0.30
0.0
0.1
0.2
0.3
0.4
0.5
0.6
0.7

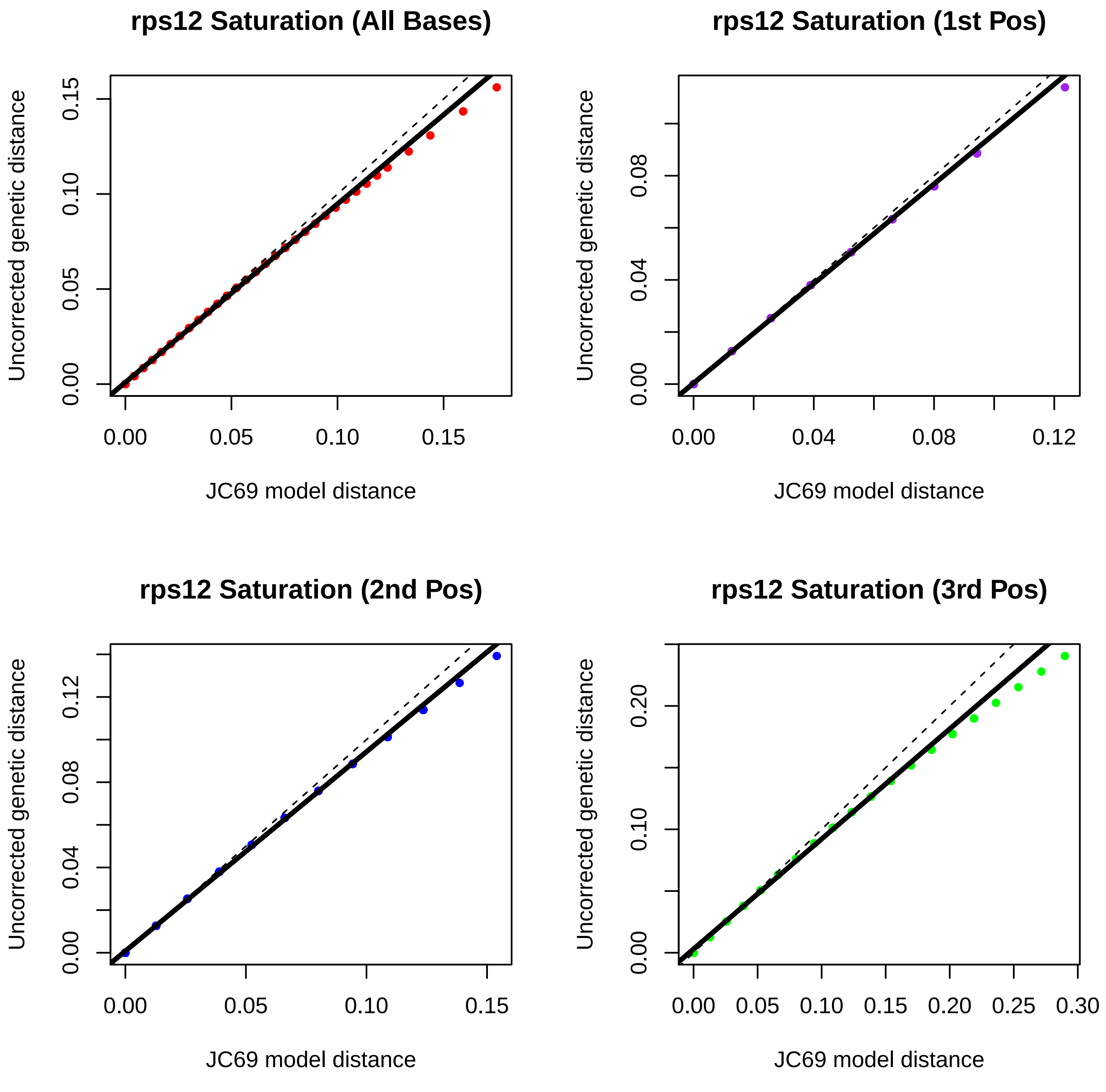

rps12 Saturation (All Bases)
rps12 Saturation (1st Pos)
rps12 Saturation (2nd Pos)
rps12 Saturation (3rd Pos)
Uncorrected genetic distance
JC69 model distance
0.00
0.05
0.10
0.15
0.04
0.08
0.12
0.20
0.25
0.30

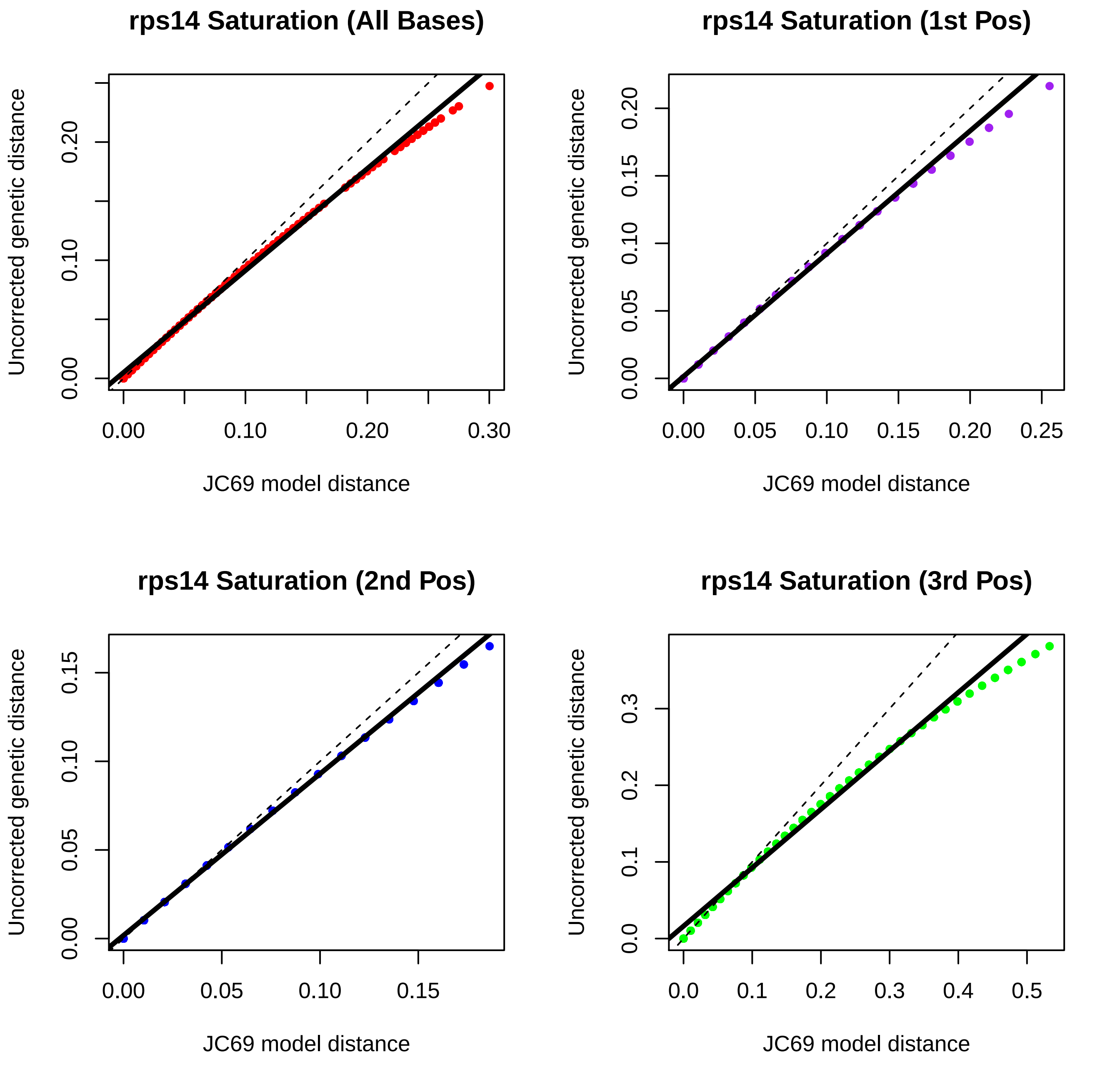

rps14 Saturation (All Bases)
rps14 Saturation (1st Pos)
rps14 Saturation (2nd Pos)
rps14 Saturation (3rd Pos)
Uncorrected genetic distance
JC69 model distance

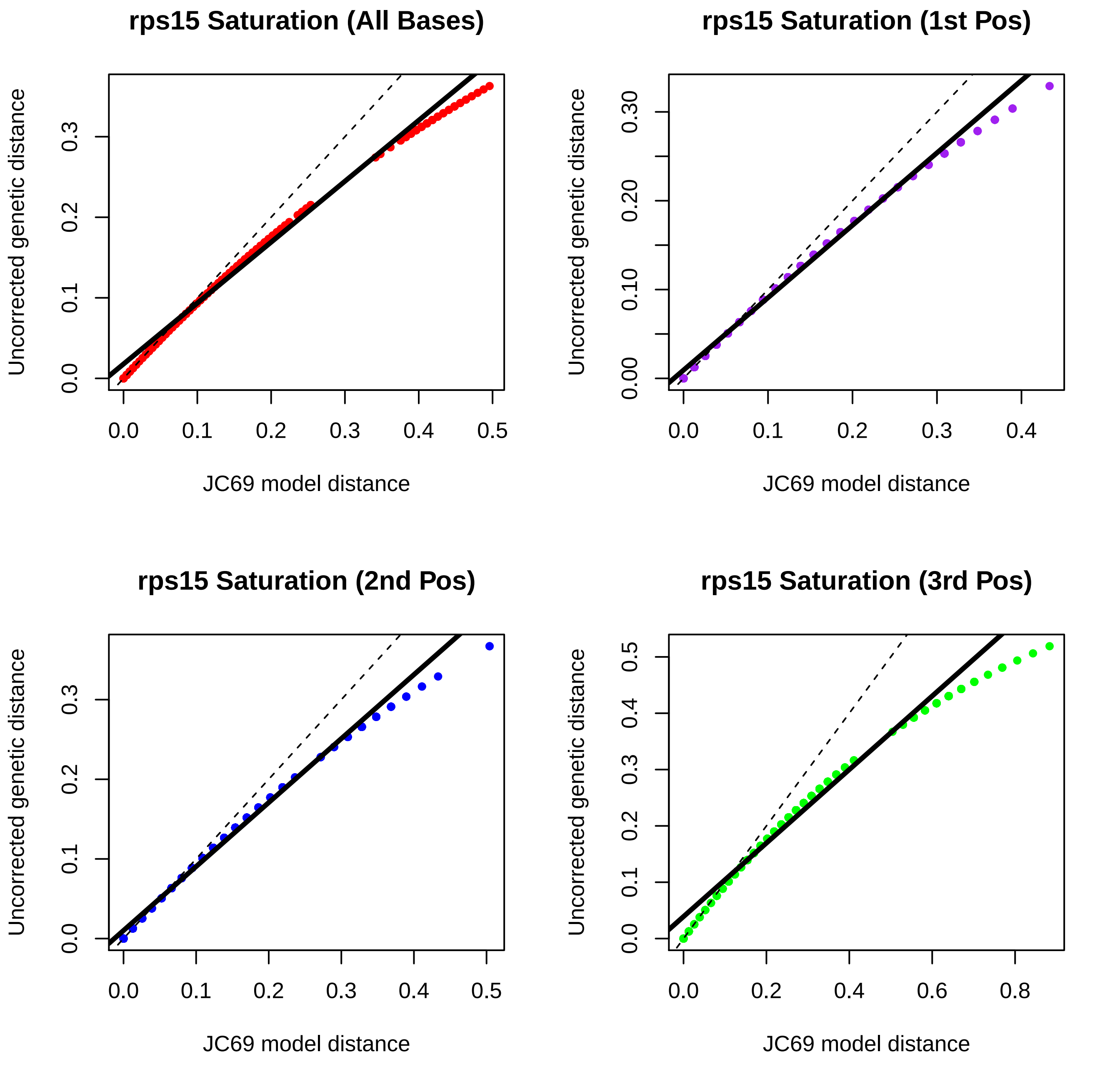
rps15 Saturation (All Bases)
rps15 Saturation (1st Pos)
rps15 Saturation (2nd Pos)
rps15 Saturation (3rd Pos)
Uncorrected genetic distance
JC69 model distance
0.0
0.1
0.2
0.3
0.4
0.5
0.00
0.10
0.20
0.30
0.6
0.8

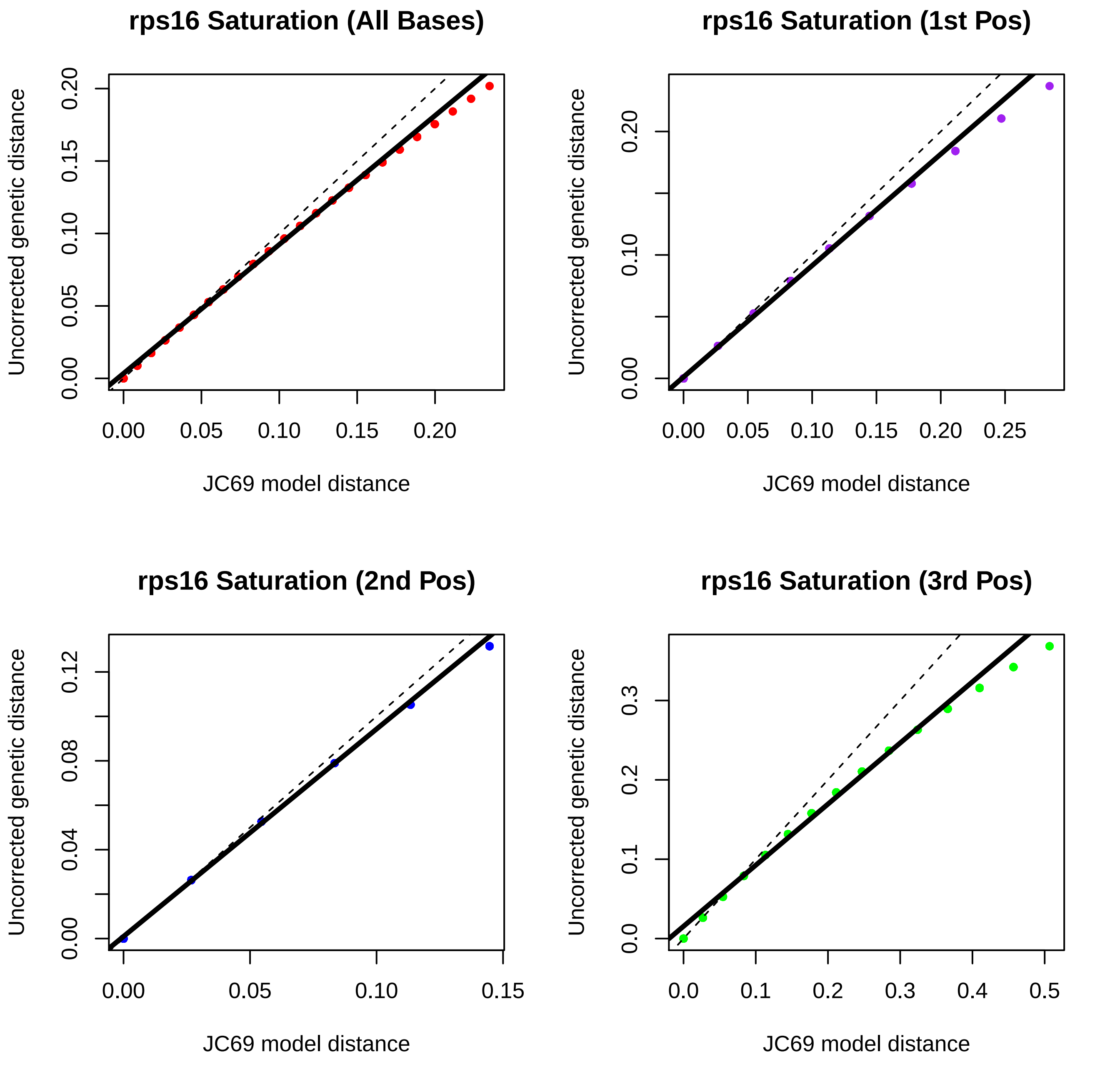
rps16 Saturation (All Bases)
rps16 Saturation (1st Pos)
rps16 Saturation (2nd Pos)
rps16 Saturation (3rd Pos)
Uncorrected genetic distance
JC69 model distance
0.00
0.05
0.10
0.15
0.20
0.25
0.04
0.08
0.12
0.0
0.1
0.2
0.3
0.4
0.5

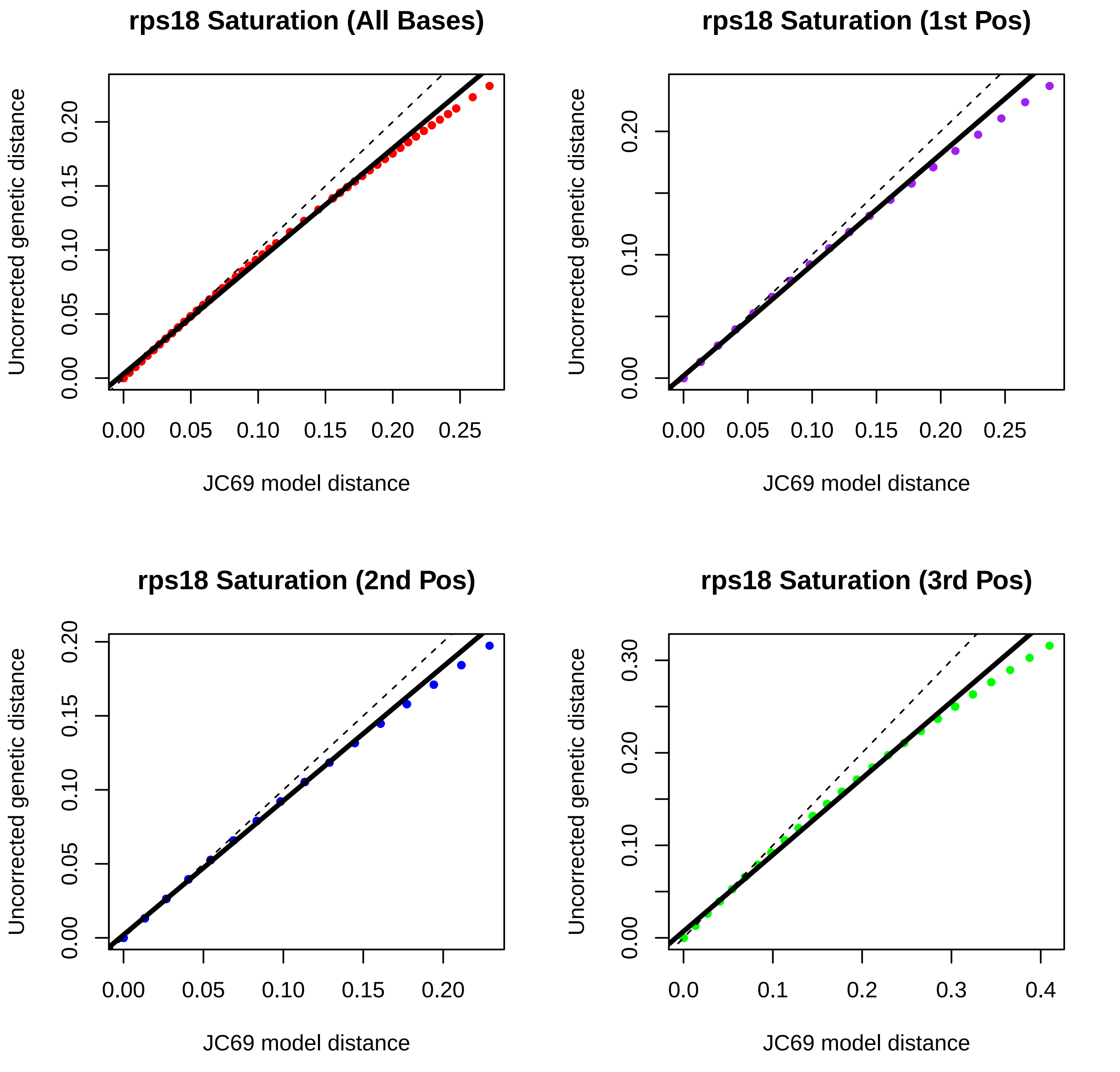
rps18 Saturation (All Bases)
rps18 Saturation (1st Pos)
rps18 Saturation (2nd Pos)
rps18 Saturation (3rd Pos)
Uncorrected genetic distance
JC69 model distance
0.00
0.05
0.10
0.15
0.20
0.25
0.30
0.0
0.1
0.2
0.3
0.4

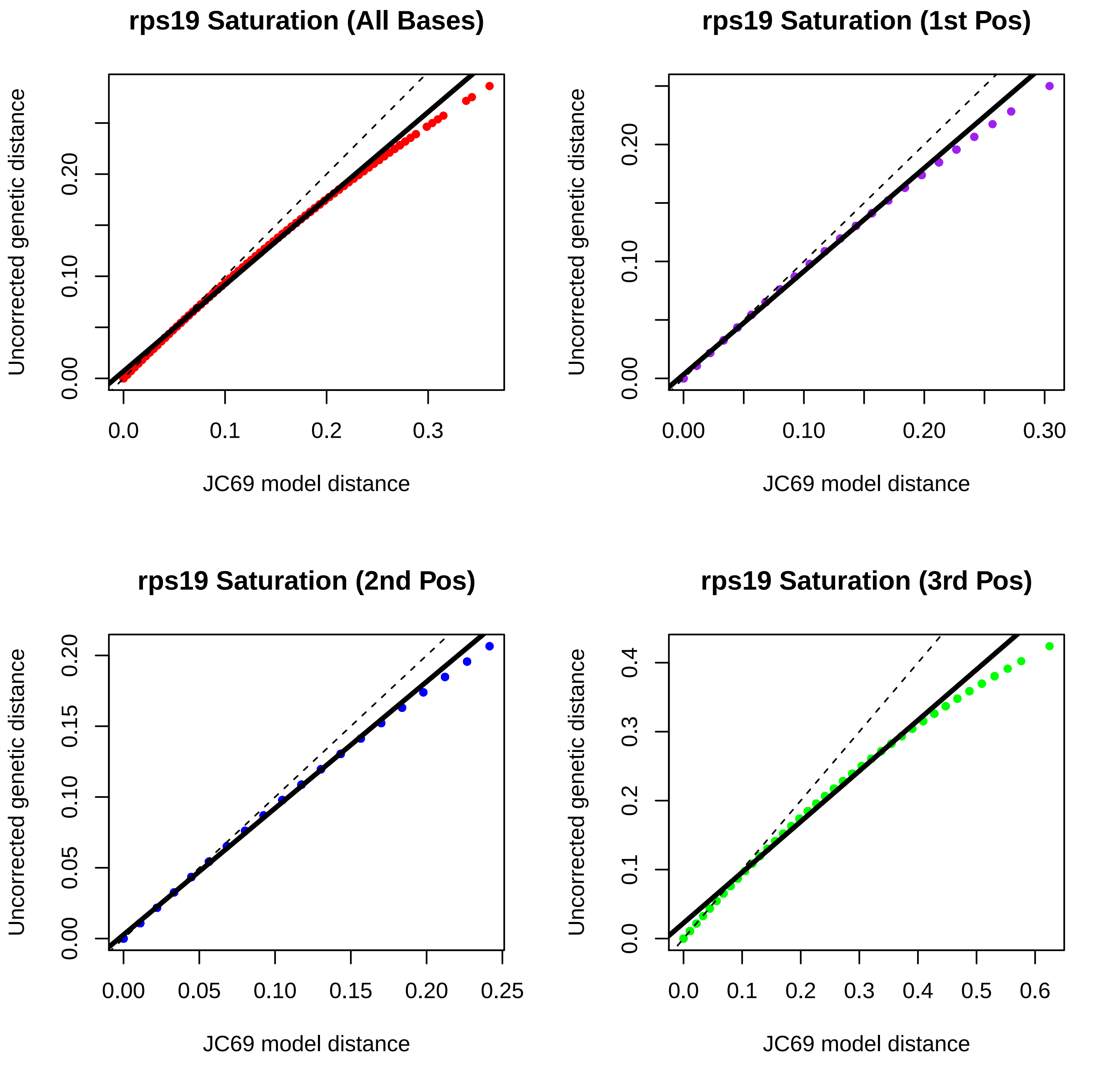
rps19 Saturation (All Bases)
Uncorrected genetic distance
0.00
0.10
0.20
0.0
0.1
0.2
0.3
JC69 model distance
rps19 Saturation (1st Pos)
Uncorrected genetic distance
0.00
0.10
0.20
0.00
0.10
0.20
0.30
JC69 model distance
rps19 Saturation (2nd Pos)
Uncorrected genetic distance
0.00
0.05
0.10
0.15
0.20
0.00
0.05
0.10
0.15
0.20
0.25
JC69 model distance
rps19 Saturation (3rd Pos)
Uncorrected genetic distance
0.0
0.1
0.2
0.3
0.4
0.0
0.1
0.2
0.3
0.4
0.5
0.6
JC69 model distance

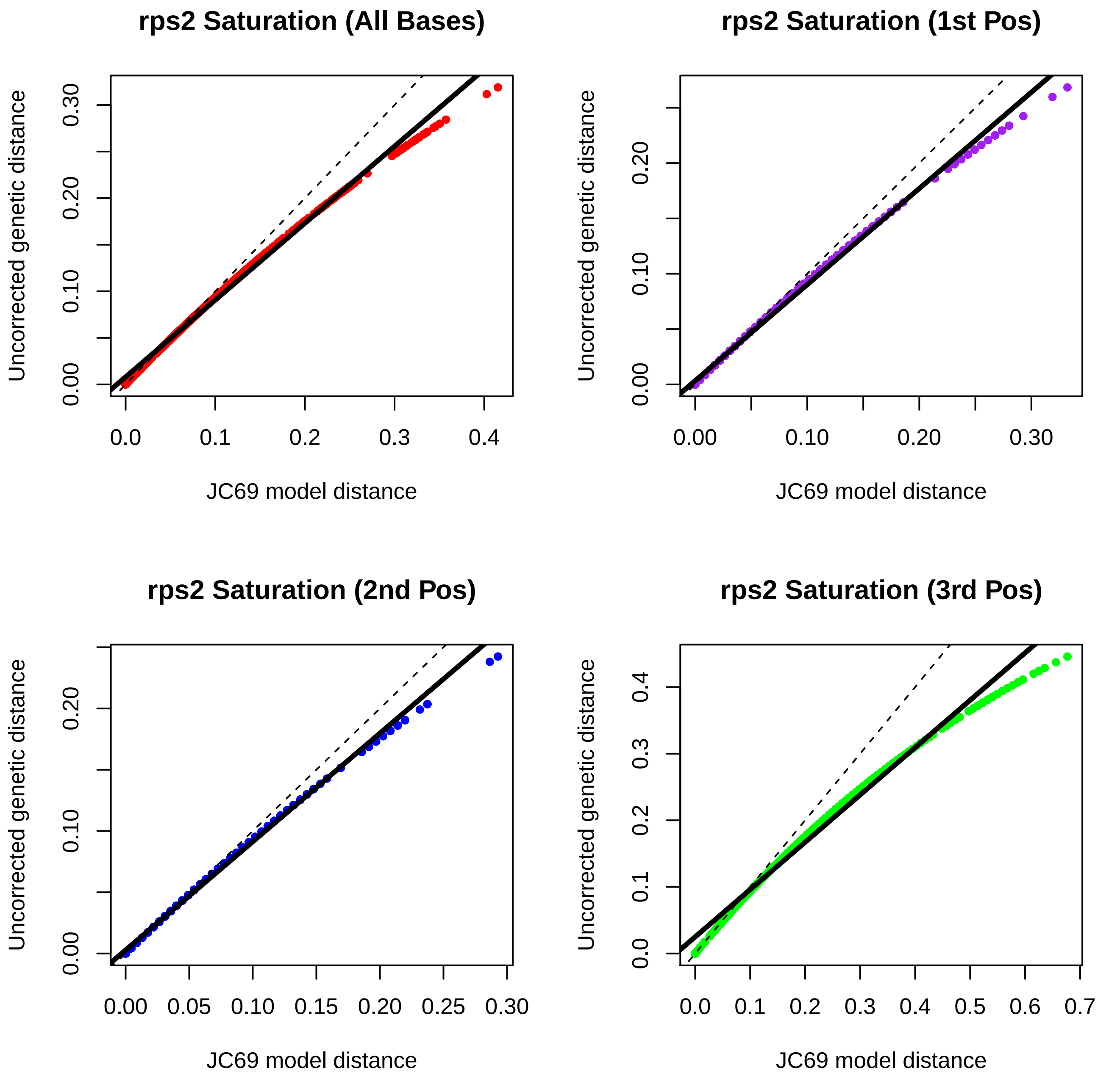
rps2 Saturation (All Bases)
rps2 Saturation (1st Pos)
rps2 Saturation (2nd Pos)
rps2 Saturation (3rd Pos)
Uncorrected genetic distance
JC69 model distance
0.00
0.10
0.20
0.30
0.0
0.1
0.2
0.3
0.4
0.05
0.15
0.25
0.5
0.6
0.7

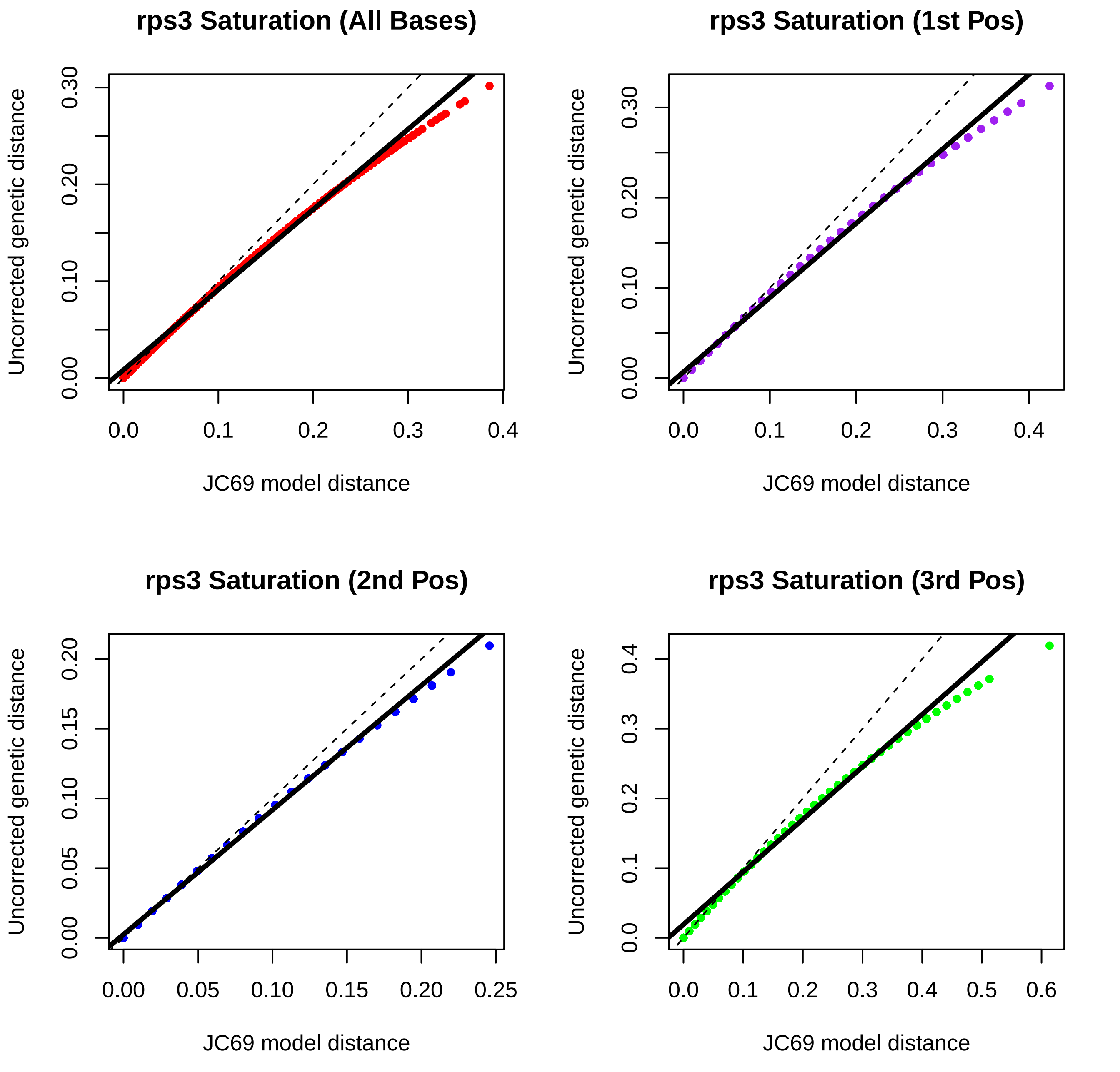

rps3 Saturation (All Bases)
Uncorrected genetic distance
0.00
0.10
0.20
0.30
0.0
0.1
0.2
0.3
0.4
JC69 model distance
rps3 Saturation (1st Pos)
Uncorrected genetic distance
0.00
0.10
0.20
0.30
0.0
0.1
0.2
0.3
0.4
JC69 model distance
rps3 Saturation (2nd Pos)
Uncorrected genetic distance
0.00
0.05
0.10
0.15
0.20
0.00
0.05
0.10
0.15
0.20
0.25
JC69 model distance
rps3 Saturation (3rd Pos)
Uncorrected genetic distance
0.0
0.1
0.2
0.3
0.4
0.0
0.1
0.2
0.3
0.4
0.5
0.6
JC69 model distance

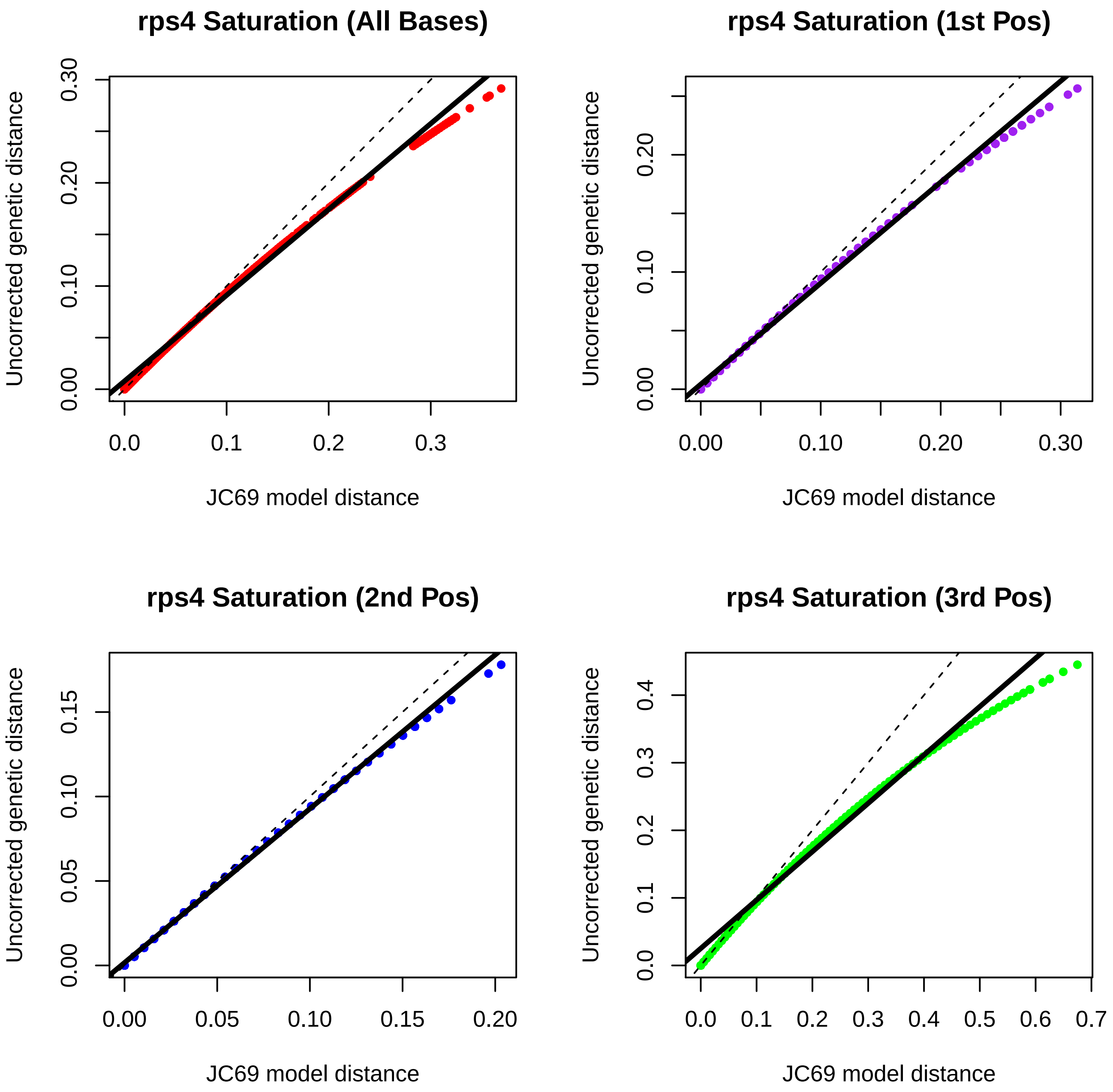
rps4 Saturation (All Bases)
rps4 Saturation (1st Pos)
rps4 Saturation (2nd Pos)
rps4 Saturation (3rd Pos)
Uncorrected genetic distance
JC69 model distance
0.00
0.10
0.20
0.30
0.0
0.1
0.2
0.3
0.05
0.15
0.4
0.5
0.6
0.7

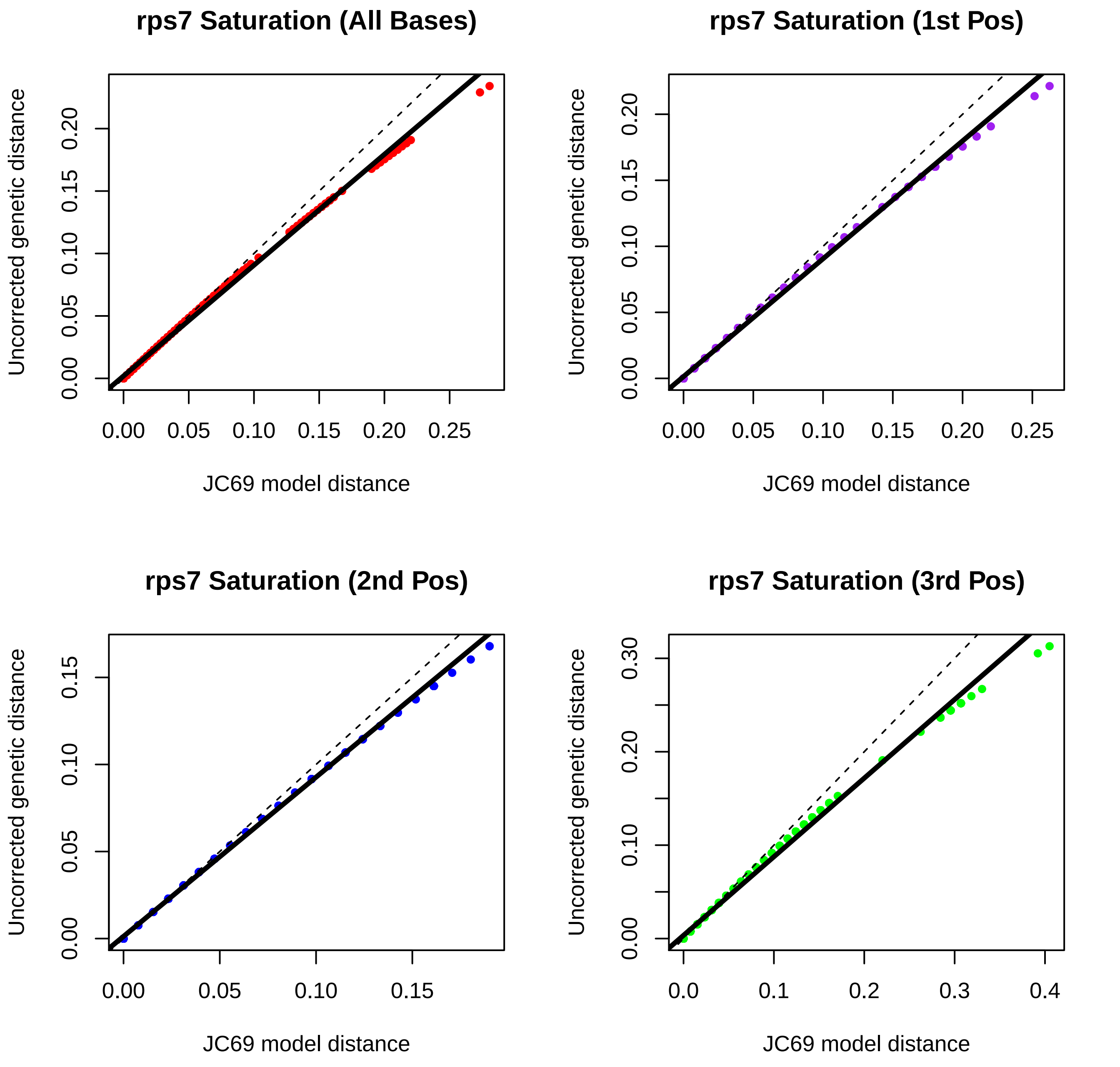
rps7 Saturation (All Bases)
rps7 Saturation (1st Pos)
rps7 Saturation (2nd Pos)
rps7 Saturation (3rd Pos)
Uncorrected genetic distance
JC69 model distance
0.00
0.05
0.10
0.15
0.20
0.25
0.30
0.0
0.1
0.2
0.3
0.4

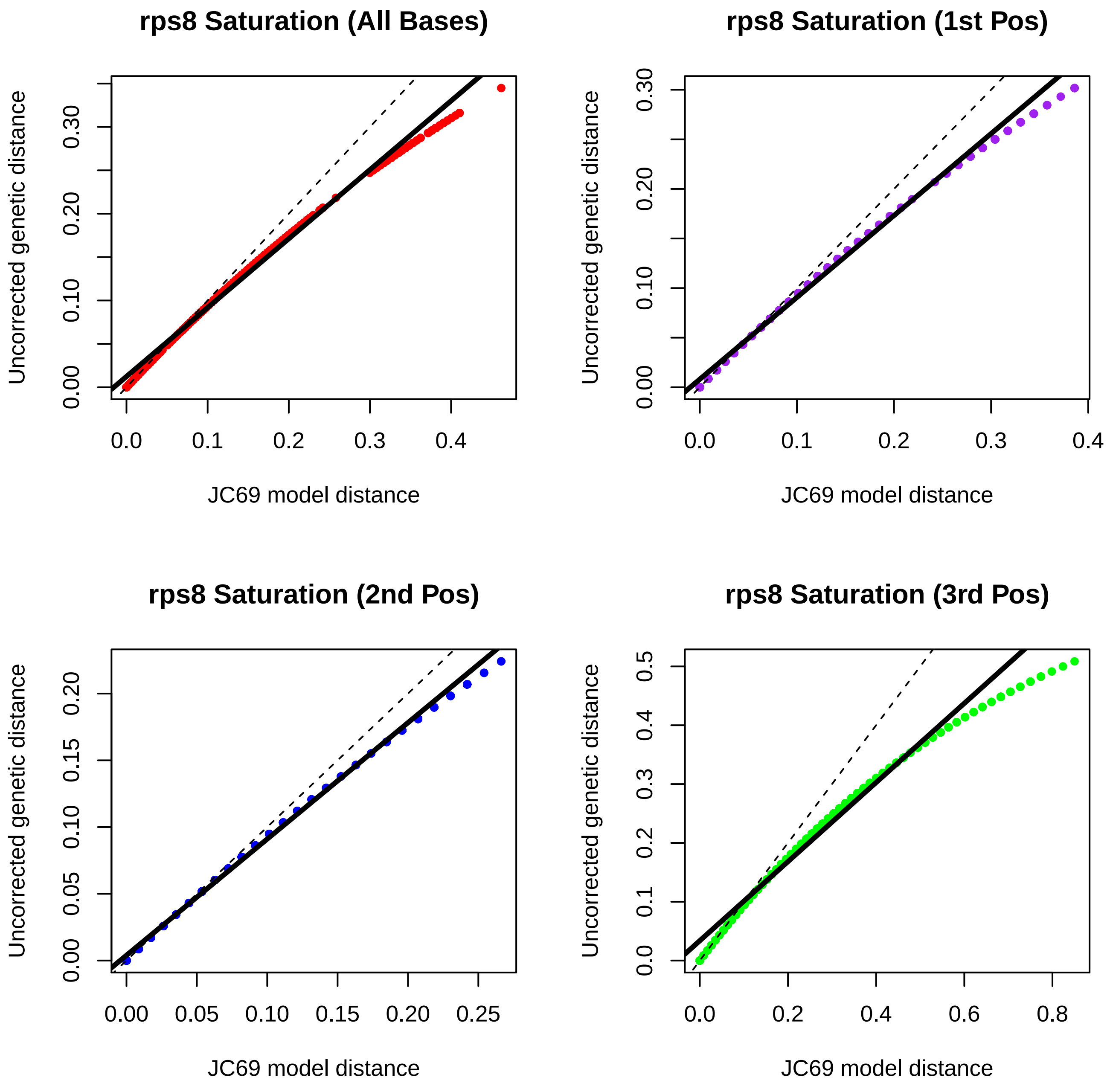
rps8 Saturation (All Bases)
rps8 Saturation (1st Pos)
rps8 Saturation (2nd Pos)
rps8 Saturation (3rd Pos)
Uncorrected genetic distance
JC69 model distance
0.00
0.10
0.20
0.30
0.0
0.1
0.2
0.3
0.4
0.05
0.15
0.25
0.5
0.6
0.8

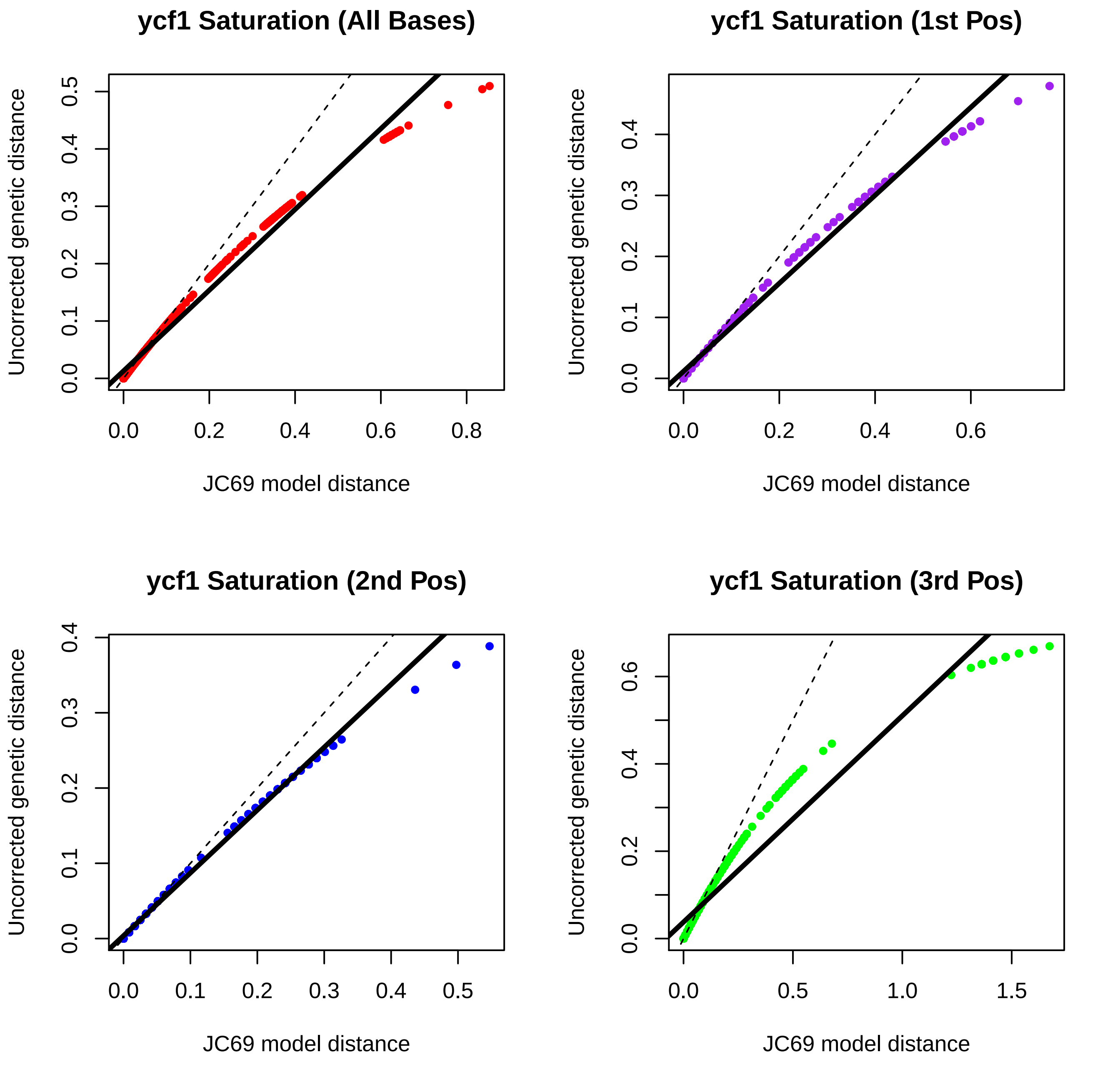

ycf1 Saturation (All Bases)
ycf1 Saturation (1st Pos)
ycf1 Saturation (2nd Pos)
ycf1 Saturation (3rd Pos)
Uncorrected genetic distance
JC69 model distance
0.0
0.1
0.2
0.3
0.4
0.5
0.6
0.8
1.0
1.5

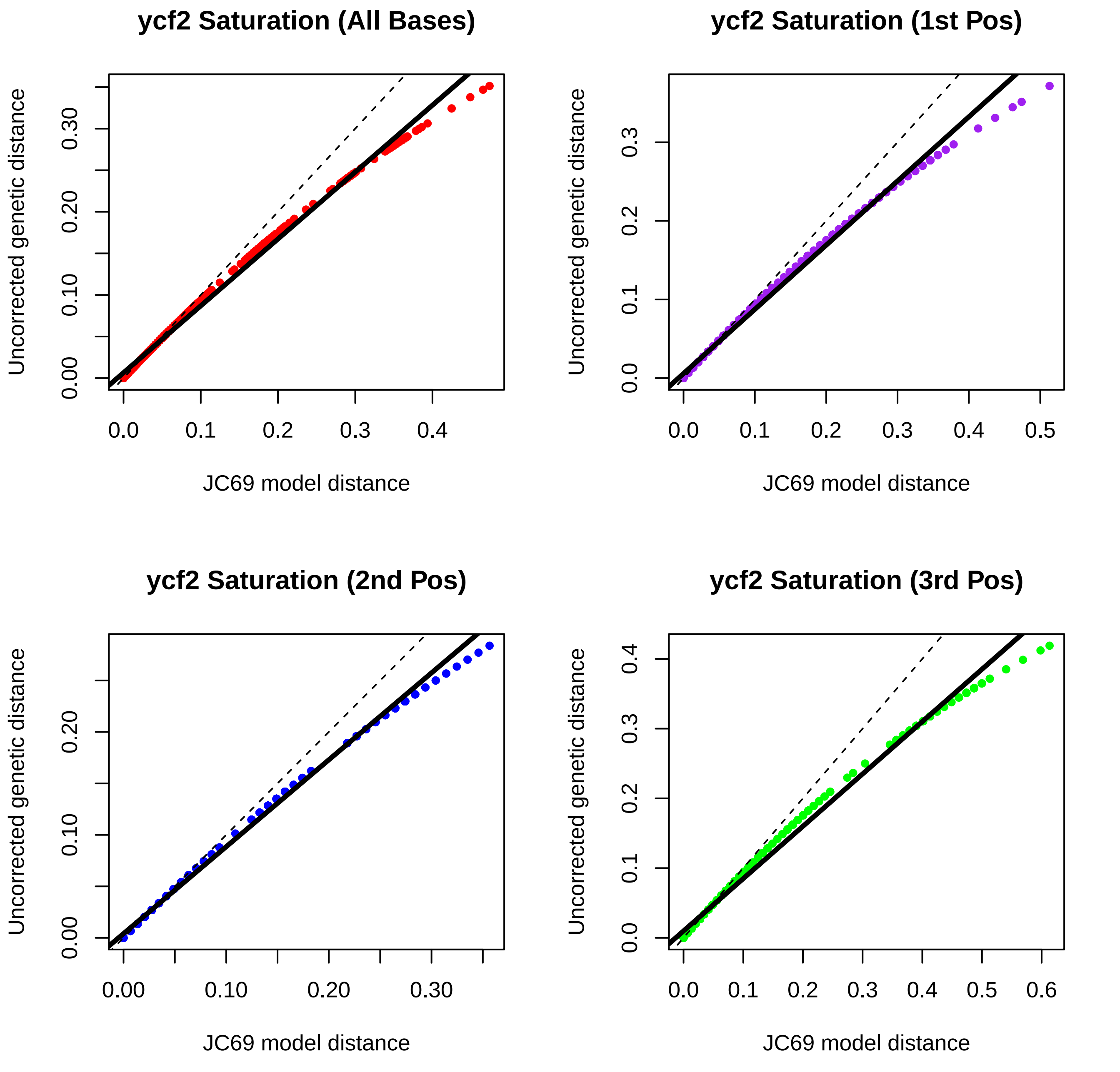

ycf2 Saturation (All Bases)
ycf2 Saturation (1st Pos)
ycf2 Saturation (2nd Pos)
ycf2 Saturation (3rd Pos)
Uncorrected genetic distance
JC69 model distance
0.00
0.10
0.20
0.30
0.0
0.1
0.2
0.3
0.4
0.5
0.6

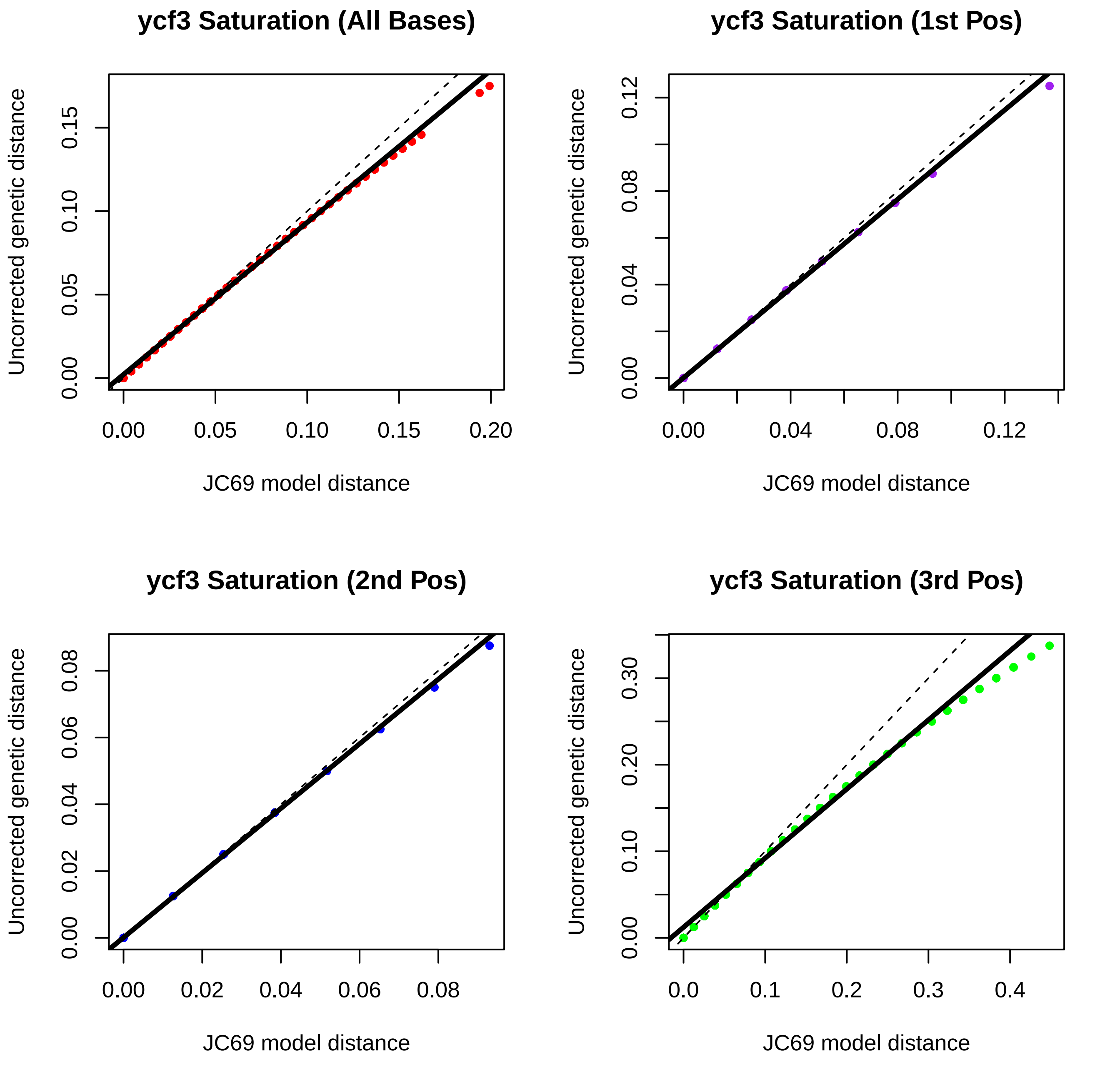
ycf3 Saturation (All Bases)
ycf3 Saturation (1st Pos)
ycf3 Saturation (2nd Pos)
ycf3 Saturation (3rd Pos)
Uncorrected genetic distance
JC69 model distance
0.00
0.05
0.10
0.15
0.20
0.04
0.08
0.12
0.02
0.06
0.0
0.1
0.2
0.3
0.4
0.00
0.10
0.20
0.30

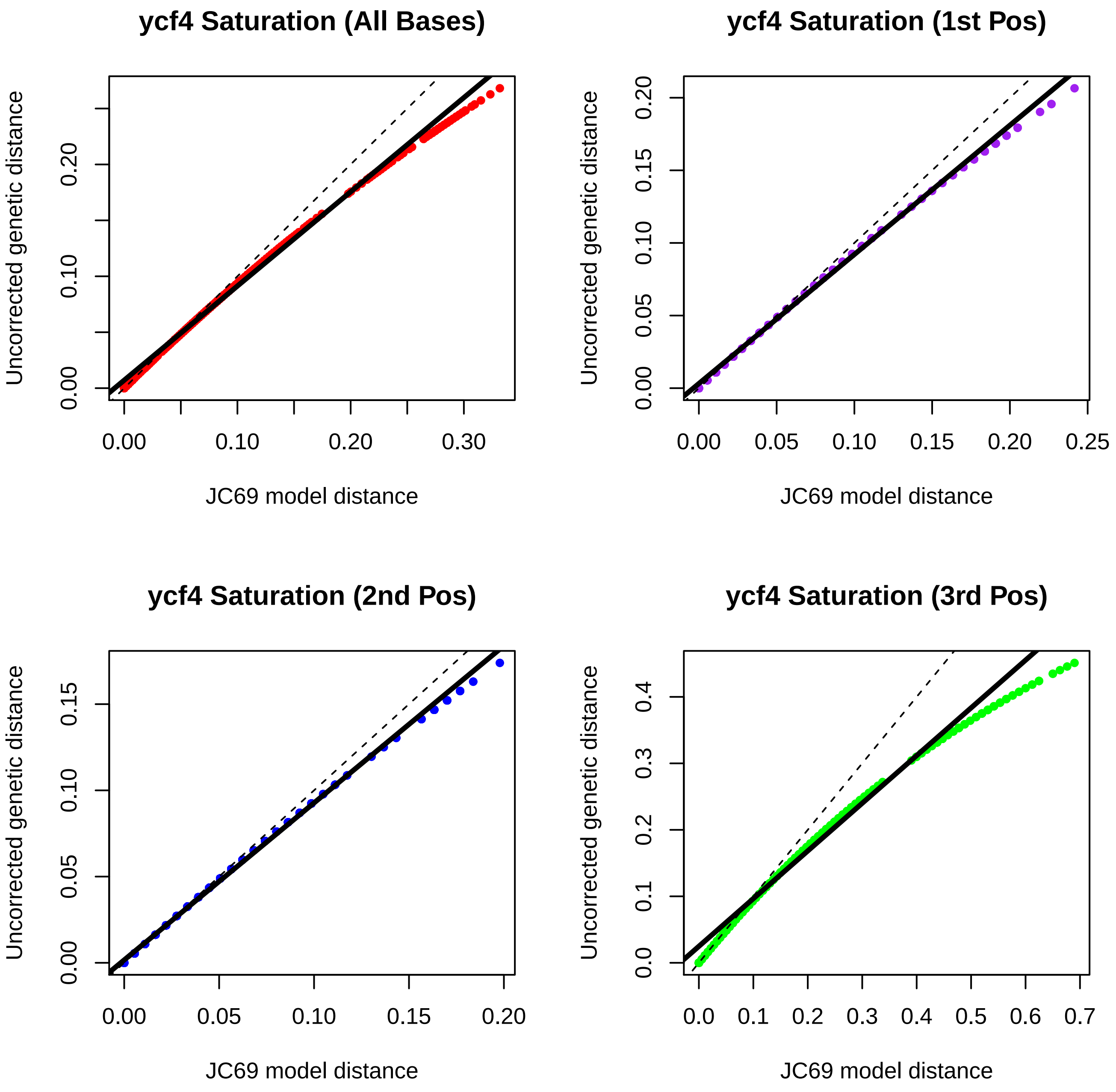
ycf4 Saturation (All Bases)
Uncorrected genetic distance
0.00
0.10
0.20
JC69 model distance
0.00
0.10
0.20
0.30
ycf4 Saturation (1st Pos)
Uncorrected genetic distance
0.00
0.05
0.10
0.15
0.20
JC69 model distance
0.00
0.05
0.10
0.15
0.20
0.25
ycf4 Saturation (2nd Pos)
Uncorrected genetic distance
0.00
0.05
0.10
0.15
JC69 model distance
0.00
0.05
0.10
0.15
0.20
ycf4 Saturation (3rd Pos)
Uncorrected genetic distance
0.0
0.1
0.2
0.3
0.4
JC69 model distance
0.0
0.1
0.2
0.3
0.4
0.5
0.6
0.7
