## Supplementary material for "Characterizing gene tree conflict in plastome-inferred phylogenies": Table S5

**Full Data**

| | Dependent variable: Total Concordant/Total Discordant | | | |
|---|---|---|---|---|
| | *logistic* | | | |
| | AA | AA BS > 70 | Nuc | Nuc BS > 70 |
| Length | 0.0002*** | 0.0002*** | 0.0002*** | 0.0002*** |
| | (0.00004) | (0.00004) | (0.00002) | (0.00002) |
| Tree_Length | 0.589*** | 0.701*** | 1.306*** | 1.649*** |
| | (0.046) | (0.063) | (0.104) | (0.127) |
| Root-to-tip Variance | -41.119*** | -74.377*** | -310.777*** | -504.933*** |
| | (5.551) | (10.768) | (34.320) | (43.698) |
| Constant | -2.012*** | -2.968*** | -1.618*** | -2.732*** |
| | (0.069) | (0.093) | (0.101) | (0.127) |
| Observations | 79 | 79 | 79 | 79 |
| Log Likelihood | -383.756 | -249.108 | -341.953 | -312.632 |
| Akaike Inf. Crit. | 775.513 | 506.217 | 691.906 | 633.263 |
| *Note:* | *p<0.1; **p<0.05; ***p<0.01 | | | |

**Supplementary Table 5:** logistic regression output for models including all predictors across both datasets both not considering and considering (BS > 70) support. Parameters are not transformed i.e. they represent the estimated effect of the predictor on log odds. Quantities in brackets are standard errors.
