## Supplementary material for "Characterizing gene tree conflict in plastome-inferred phylogenies": Table S6

**Full Data**

| | Dependent variable: Total Concordant/Total Discordant | | | |
|---|---|---|---|---|
| | *glm: quasibinomial* | | | |
| | *link = logit* | | | |
| | AA | AA BS > 70 | Nuc | Nuc BS > 70 |
| Length | $0.0002^{**}$ | $0.0002^{*}$ | $0.0002^{***}$ | $0.0002^{***}$ |
| | (0.0001) | (0.0001) | (0.00004) | (0.00004) |
| Tree_Length | $0.589^{***}$ | $0.701^{***}$ | $1.306^{***}$ | $1.649^{***}$ |
| | (0.123) | (0.186) | (0.222) | (0.266) |
| Root-to-tip Variance | $-41.119^{***}$ | $-74.377^{**}$ | $-310.777^{***}$ | $-504.933^{***}$ |
| | (14.787) | (31.532) | (73.500) | (91.484) |
| Constant | $-2.012^{***}$ | $-2.968^{***}$ | $-1.618^{***}$ | $-2.732^{***}$ |
| | (0.183) | (0.273) | (0.215) | (0.266) |
| Dispersion | 7.096 | 8.575 | 4.587 | 4.383 |
| Observations | 79 | 79 | 79 | 79 |
| *Note:* | $^{*}$p<0.1; $^{**}$p<0.05; $^{***}$p<0.01 | | | |

**Supplementary Table 6:** quasibinomial logistic regression output for models with all predictors across all datasets, both not considering and considering (BS > 70) support. Parameters are not transformed i.e. they represent the estimated effect of the predictor on log odds. Dispersion gives the estimated quasibinomial dispersion parameter.
