## Supplementary material for "Characterizing gene tree conflict in plastome-inferred phylogenies": Table S7

**Reduced Data**

| | Dependent variable: Total Concordant/Total Discordant | | | |
|---|---|---|---|---|
| | *logistic* | | | |
| | AA | AA BS > 70 | Nuc | Nuc BS > 70 |
| Length | 0.002*** | 0.002*** | 0.001*** | 0.001*** |
| | (0.0001) | (0.0002) | (0.0001) | (0.0001) |
| Tree_Length | 0.772*** | 0.677*** | 1.138*** | 1.138*** |
| | (0.068) | (0.083) | (0.145) | (0.145) |
| Root-to-tip Variance | -98.944*** | -76.416*** | -234.793*** | -234.793*** |
| | (15.412) | (20.333) | (55.806) | (55.806) |
| Constant | -2.546*** | -3.557*** | -1.894*** | -1.894*** |
| | (0.083) | (0.111) | (0.113) | (0.113) |
| Observations | 75 | 75 | 75 | 75 |
| Log Likelihood | -257.248 | -156.195 | -253.832 | -253.832 |
| Akaike Inf. Crit. | 522.496 | 320.391 | 515.665 | 515.665 |
| *Note:* | *p<0.1; **p<0.05; ***p<0.01 | | | |

**Supplementary Table 7:** logistic regression results for models with all predictors on datasets excluding influential and outlier observations, both considering and not considering (BS > 70) support. Parameters are not transformed i.e. they represent the estimated effect of the predictor on log odds. Quantities in brackets are standard errors.
