## Supplementary material for "Characterizing gene tree conflict in plastome-inferred phylogenies": Table S8

**Reduced Data**

| | Dependent variable: Total Concordant/Total Discordant | | | |
|---|---|---|---|---|
| | *glm: quasibinomial* *link = logit* | | | |
| | AA | AA BS > 70 | Nuc | Nuc BS > 70 |
| Length | 0.002*** | 0.002*** | 0.001*** | 0.001*** |
| | (0.0003) | (0.0002) | (0.0001) | (0.0001) |
| Tree_Length | 0.772*** | 0.677*** | 1.138*** | 1.138*** |
| | (0.129) | (0.094) | (0.242) | (0.242) |
| Root-to-tip Variance | -98.944*** | -76.416*** | -234.793** | -234.793** |
| | (29.067) | (22.918) | (93.245) | (93.245) |
| Constant | -2.546*** | -3.557*** | -1.894*** | -1.894*** |
| | (0.156) | (0.126) | (0.189) | (0.189) |
| Dispersion | 3.557 | 1.27 | 2.792 | 2.792 |
| Observations | 75 | 75 | 75 | 75 |
| *Note:* | *p<0.1; **p<0.05; ***p<0.01 | | | |

**Supplementary Table 8:** quasibinomial logistic regression output for both datasets excluding influential observations and outlier genes, both not considering and considering (BS > 70) support. Parameters are not transformed i.e. they represent the estimate effect of the predictor on log odds. Quantities in brackets are standard errors. Dispersion gives the estimated quasibinomial dispersion parameter.
