## Supplementary material for "Characterizing gene tree conflict in plastome-inferred phylogenies": Table S9

**Full Data**

| | Dependent Variable: Total Concordant/Total Discordant | | | |
|---|---|---|---|---|
| | *logistic* | | | |
| | AA | AA BS > 70 | Nuc | Nuc BS > 70 |
| Length | 0.0002*** | 0.0002*** | 0.0001*** | 0.0001*** |
| | (0.00004) | (0.00003) | (0.00002) | (0.00001) |
| Tree_Length | 0.266*** | 0.223*** | 0.567*** | 0.412*** |
| | (0.027) | (0.032) | (0.063) | (0.068) |
| Constant | -1.756*** | -2.575*** | -1.187*** | -1.867*** |
| | (0.060) | (0.076) | (0.087) | (0.097) |
| Observations | 79 | 79 | 79 | 79 |
| Log Likelihood | -430.988 | -295.026 | -390.554 | -395.112 |
| Akaike Inf. Crit. | 867.975 | 596.053 | 787.109 | 796.225 |
| *Note:* | *p<0.1; **p<0.05; ***p<0.01 | | | |

**Supplementary Table 9:** logistic regression output for models of alignment length and tree length across all datasets both not considering and considering (BS > 70) support. Parameters are not transformed i.e. they represent the effect of the predictor on log odds. Quantities in brackets are standard errors.
