## Supplementary material for "Characterizing gene tree conflict in plastome-inferred phylogenies": Table S10

**Full Data**

| | Dependent Variable: Total Concordant/Total Discordant | | | |
|---|---|---|---|---|
| | *glm: quasibinomial* | | | |
| | *link = logit* | | | |
| | AA | AA BS > 70 | Nuc | Nuc BS > 70 |
| Length | 0.0002** | 0.0002** | 0.0001*** | 0.0001*** |
| | (0.0001) | (0.0001) | (0.0001) | (0.00004) |
| Tree_Length | 0.266*** | 0.223*** | 0.567*** | 0.412** |
| | (0.076) | (0.072) | (0.166) | (0.176) |
| Constant | -1.756*** | -2.575*** | -1.187*** | -1.867*** |
| | (0.168) | (0.173) | (0.229) | (0.249) |
| Dispersion | 7.827 | 5.175 | 6.862 | 6.621 |
| Observations | 79 | 79 | 79 | 79 |
| *Note:* | *p<0.1; **p<0.05; ***p<0.01 | | | |

**Supplementary Table 10:** quasibinomial logistic regression output for models of alignment length and tree length across all datasets both not considering and considering (BS > 70) support. Parameters are not transformed i.e. they represent the effect of the predictor on log odds. Quantities in brackets are standard errors. Dispersion gives the estimated quasibinomial dispersion parameter.
