## Supplementary material for "Characterizing gene tree conflict in plastome-inferred phylogenies": Table S11

**Reduced Data**

| | Dependent Variable: Total Concordant/Total Discordant | | | |
|---|---|---|---|---|
| | *logistic* | | | |
| | AA | AA BS > 70 | Nuc | Nuc BS > 70 |
| Length | $0.002^{***}$ | $0.002^{***}$ | $0.001^{***}$ | $0.001^{***}$ |
| | (0.0001) | (0.0002) | (0.00005) | (0.00005) |
| Tree_Length | $0.291^{***}$ | $0.277^{***}$ | $0.509^{***}$ | $0.414^{***}$ |
| | (0.029) | (0.035) | (0.067) | (0.075) |
| Constant | $-2.390^{***}$ | $-3.414^{***}$ | $-1.580^{***}$ | $-2.516^{***}$ |
| | (0.077) | (0.105) | (0.095) | (0.113) |
| Observations | 77 | 77 | 77 | 77 |
| Log Likelihood | -296.628 | -177.738 | -279.551 | -248.075 |
| Akaike Inf. Crit. | 599.256 | 361.476 | 565.103 | 502.151 |
| *Note:* | $^{*}p<0.1$; $^{**}p<0.05$; $^{***}p<0.01$ | | | |

**Supplementary Table 11:** logistic regression output for models of alignment length and tree length on reduced datasets excluding outlier genes and influential observations. Parameters are not transformed, i.e. they represent the effect of the predictor on log odds. Quantities in brackets are standard errors.
