## Supplementary material for "Characterizing gene tree conflict in plastome-inferred phylogenies": Table S12

**Reduced Data**

| | Dependent Variable: Total Concordant/Total Discordant | | | |
|---|---|---|---|---|
| | *glm: quasibinomial* | | | |
| | *link = logit* | | | |
| | AA | AA BS > 70 | Nuc | Nuc BS > 70 |
| Length | $0.002^{***}$ | $0.002^{***}$ | $0.001^{***}$ | $0.001^{***}$ |
| | (0.0003) | (0.0002) | (0.0001) | (0.0001) |
| Tree_Length | $0.291^{***}$ | $0.277^{***}$ | $0.509^{***}$ | $0.414^{***}$ |
| | (0.059) | (0.046) | (0.121) | (0.127) |
| Constant | $-2.390^{***}$ | $-3.414^{***}$ | $-1.580^{***}$ | $-2.516^{***}$ |
| | (0.158) | (0.137) | (0.171) | (0.190) |
| Dispersion | 4.25 | 1.684 | 3.277 | 2.841 |
| Observations | 77 | 77 | 77 | 77 |
| *Note:* | $^{*}p<0.1$; $^{**}p<0.05$; $^{***}p<0.01$ | | | |

**Supplementary Table 12:** Quasibinomial logistic regression output for models of alignment length and tree length across all datasets both not considering and considering (BS > 70) support. Parameters are not transformed, i.e. they represent the estimated effect of the predictor on the log odds. Quantities in brackets are standard errors. Dispersion gives the estimated quasibinomial dispersion parameter.
